# Molecular characterization reveals genomic and transcriptomic subtypes of metastatic urothelial carcinoma

**DOI:** 10.1101/2021.03.17.435757

**Authors:** J. Alberto Nakauma-González, Maud Rijnders, Job van Riet, Michiel S. van der Heijden, Jens Voortman, Edwin Cuppen, Niven Mehra, Sandra van Wilpe, Sjoukje F. Oosting, L. Lucia Rijstenberg, Hans M. Westgeest, Ellen C. Zwarthoff, Ronald de Wit, Astrid A.M. van der Veldt, Harmen J. G. van de Werken, Martijn P. J. Lolkema, Joost L. Boormans

## Abstract

**Background:** Molecular characterization of primary urothelial carcinoma (UC) revealed molecular subtypes with different genomic, transcriptomic, and clinicopathological characteristics, which might guide therapeutic decision making. A comprehensive molecular characterization of metastatic UC (mUC), however, is currently lacking in the literature. Because of the lethality of mUC, with few therapeutic options available for patients, a multi-omics characterization of mUC could aid to improve patient selection for new and existing therapies.

**Methods:** To define the molecular landscape of mUC and to identify potential targets for therapy, we performed whole genome DNA sequencing on fresh-frozen metastatic tumor biopsies of 116 mUC patients, and mRNA sequencing on 90 matched biopsies.

**Results:** Hierarchical clustering based on mutational signatures revealed two major genomic subtypes. The most prevalent subtype (67%) consisted almost exclusively of tumors with high APOBEC mutagenesis. APOBEC mutagenesis was detected in 91% of the samples, and appeared to be an ongoing process in mUC based on analysis of eight patients from whom serial biopsies were obtained during treatment. Contrary to the overall distribution of mutations, APOBEC associated mutations occurred throughout the genome, and independently of predicted accessible or transcribed genomic regions, suggesting that these mutations were generated during replication. Transcriptomic analysis revealed five mRNA-based subtypes: two luminal subtypes (40%), a stroma-rich (24%), basal/squamous (23%), and non-specified subtype (12%). The transcriptomic subtypes were different regarding driver gene alterations (e.g. *ELF3* and *TSC1*), gene amplifications (*NECTIN4* and *PPARG*), pathway activity, and immune cell infiltration. By integrating the genomic and transcriptomic data, potential therapeutic options per transcriptomic subtype and individual patient were proposed.

**Conclusions:** This study expands our knowledge on the molecular landscape of mUC, and serves as a reference for subtype-oriented and patient-specific research on the etiology of mUC, and for novel drug development.

**Trial registration:** The mUC cohort studied here is part of the Netherlands nationwide study of the center for personalized cancer treatment consortium (CPCT-02 Biopsy Protocol, NCT01855477), and the Drug Rediscovery Protocol (DRUP Trial, NCT02925234).

## Background

Urothelial cancer (UC) is a molecularly and clinically heterogeneous disease. Non-muscle invasive bladder cancer (NMIBC) is characterized by excellent survival but high recurrence rates, whereas muscle-invasive bladder cancer (MIBC) has high metastatic potential and poor patient outcome despite aggressive local and systemic treatment [1]. Comprehensive molecular profiling of UC has been restricted to NMIBC [2] and localized MIBC [3]. At the genomic level, NMIBC is characterized by frequent *FGFR3* and *PIK3CA* mutations, whereas *TP53* mutations are uncommon [1]. In MIBC, *TP53* is the most commonly mutated gene [4]. *The Cancer Genome Atlas* (TCGA) initiative molecularly characterized 412 chemotherapy-naïve primary MIBC patients and found that a subgroup of patients had high Apolipoprotein B mRNA Editing Catalytic Polypeptide-like (APOBEC) signature mutagenesis and high mutational burden. The patients in this subgroup had an excellent 5-year overall survival rate of 75% [3]. At the transcriptomic level, MIBC can be stratified into basal and luminal subtypes. A recent study proposed a consensus molecular classification of MIBC, consisting of six subtypes: basal/squamous, luminal non-specified, luminal papillary, luminal unstable, neuroendocrine-like (NE-like), and stroma-rich [5]. These subtypes included distinct genomic alterations and clinical and pathological characteristics, which might guide therapeutic decision making.

A comprehensive multi-omics characterization of mUC has not yet been performed. A previous study reported the clonal evolution of mUC by whole-exome sequencing (WES) in a cohort of 32 chemotherapy-treated patients, and showed that APOBEC mutagenesis was clonally enriched in chemotherapy-treated mUC [6].

Expanding the knowledge on the molecular characteristics of mUC is crucial for more robust and accurate patient stratification and for rational drug development paths that will eventually improve the outcome of this lethal cancer. In the present study, we conducted a comprehensive genomic and transcriptomic analysis of freshly obtained metastatic biopsies of 116 mUC patients, with the aim of identifying key molecular insights into tumorigenesis and defining molecular subtypes of mUC.

## Methods

### Patient cohort and study procedures

Between 07 June 2012 up to and including 28 February 2019, patients with advanced or mUC (n = 210) from 23 Dutch hospitals (Fig. 1a, Fig. S1) who were scheduled for 1^st^ or 2^nd^ line palliative systemic treatment were included in the Netherlands nationwide study of the Center for Personalized Cancer Treatment (CPCT) consortium (CPCT-02 Biopsy Protocol, NCT01855477 [7]) and the Drug Rediscovery Protocol (DRUP Trial, NCT02925234), which aimed to analyze the cancer genome and transcriptome of patients with advanced cancer. The CPCT-02 and DRUP study protocols were approved by the medical ethics review board of the University Medical Center Utrecht and the Netherlands Cancer Institute, respectively. Patients eligible for inclusion were those aged ≥ 18 years old, with locally advanced or mUC, from whom a histological tumor biopsy could be safely obtained, and whom had an indication for initiation of a new line of systemic treatment with anti-cancer agents. Written informed consent was obtained from all participants prior to inclusion in the trial; the studies comply with all relevant ethical regulations. Tumor biopsies and matched normal blood samples were collected following a standardized procedure described by the Hartwig Medical Foundation (HMF; https://www.hartwigmedicalfoundation.nl; [7]). Whole genome sequencing (WGS) was successfully

**Figure 1.**
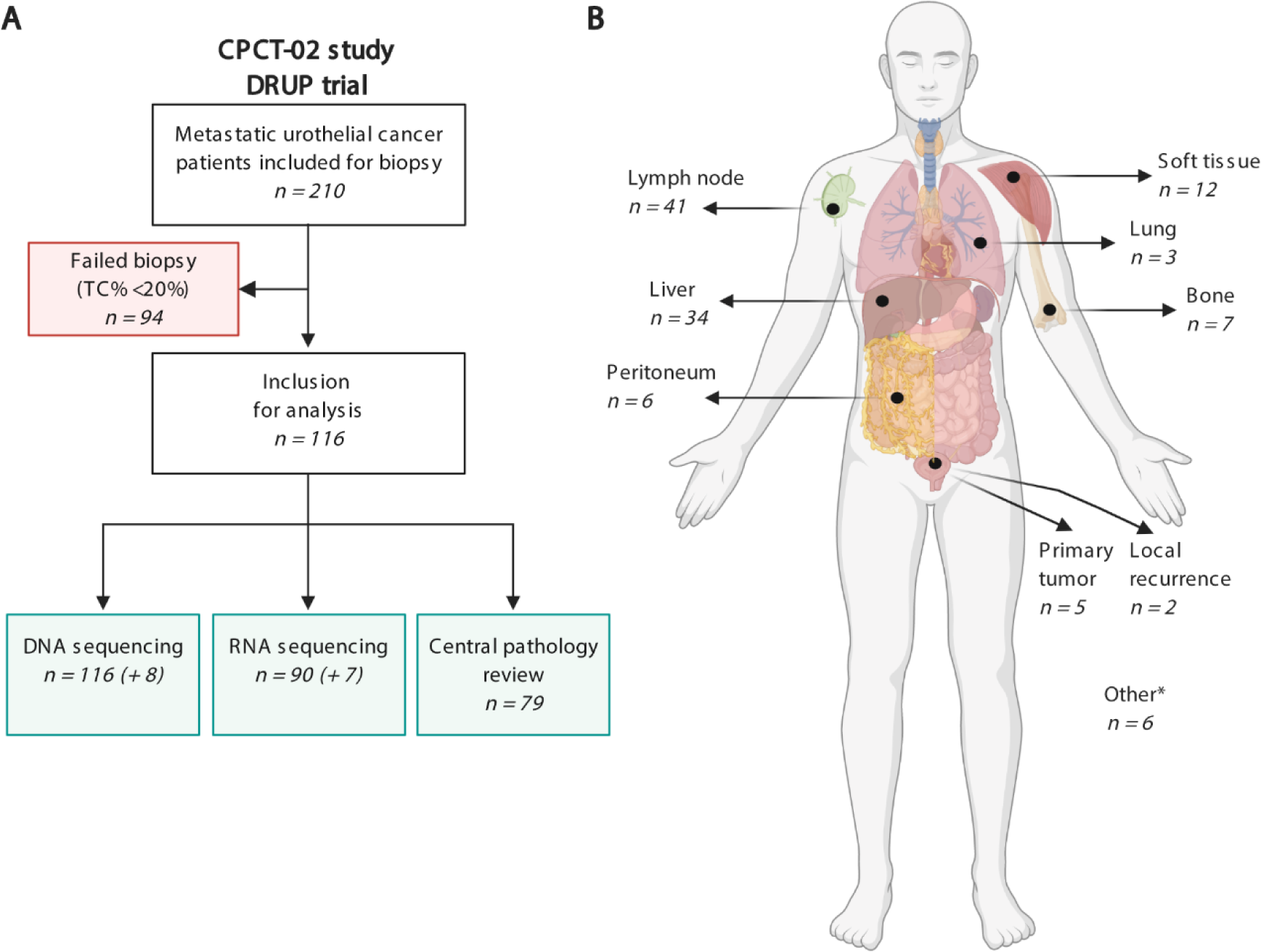
Overview of the study design and biopsy sites of 116 patients with metastatic urothelial cancer. a) Flowchart of patient inclusion. Patients with advanced or metastatic urothelial cancer who were scheduled for systemic palliative treatment were selected from the prospective Center for Personalized Cancer Treatment (CPCT-02) patient cohort and the Drug Rediscovery Protocol (DRUP Trial (n = 210). Patients were excluded if the tumor cell percentage in the biopsy was <20%, resulting in WGS data and RNA-seq for 116 and 90 patients, respectively. Tissue slides of 79 patients were available for central pathology review (primary tumor and/or metastatic biopsy). DNA +8 and RNA +7 indicate the numbers of patients from whom a second biopsy was obtained at disease progression. b)Overview of the number of biopsies per site analyzed by WGS. * Other biopsy sites included abdominal or pelvic masses (n = 3), adrenal gland (n = 1), and brain (n = 1), or unspecified biopsy site (n = 1).

performed on DNA from freshly obtained biopsies from metastatic sites in 116 mUC patients (124 samples), and matched RNA-sequencing (RNA-seq) data was available for 90 patients (97 samples; Fig. 1a). Patient characteristics are described in Table S1.1. Biopsies were obtained from a safely accessible site, including lymph nodes, liver, bone and other organs (Fig. 1b). In five patients, a tumor biopsy was obtained from the primary bladder or upper urinary tract tumor as no safely accessible metastatic lesion was present. In two patients, a biopsy was obtained from a local recurrence after cystectomy and nephrectomy, respectively (Table S1.2). Sequential biopsies of a metastatic lesion taken at the time of clinical or radiological disease progression from eight patients were additionally sequenced. This study extends the pan-cancer analysis of Priestley *et al*., 2019, in which WGS (but not RNA-seq) data of 72 mUC patients included in the current cohort were initially analyzed (Table S1.2).

### Central pathology review

Tumor tissue slides for central pathological revision of the diagnosis of UC was available for 79/116 patients. Hematoxylin and eosin (H&E) stained slides from primary tumor tissues (cystectomy and transurethral resection specimens of the bladder, n = 23 patients), metastatic tumor biopsies (n = 15 patients), or both (n = 41 patients) were requested from the Nationwide Network and Registry of Histo- and Cytopathology in the Netherlands (PALGA) [8]. Tissue slides and corresponding pathological reports were provided anonymously. All patient materials used for central pathology review were obtained within the CPCT-02 biopsy protocol, the DRUP trial, or during routine patient care, and the use of these materials for research purposes was approved by the medical ethics review board of the Erasmus University Medical Center, Rotterdam, the Netherlands (MEC-2019-0188). H&E slides were reviewed by an expert genitourinary pathologist (LLB), and used for re-evaluation of the diagnosis and description of aberrant histology (Tables S1.3 and S1.4). Tumors were classified as pure UC (n = 66), or predominant UC with variant histology (n = 9 squamous, n = 3 neuro-endocrine, n = 1 micropapillary UC), and pure squamous cell bladder carcinomas (n = 3). In patients for whom both the primary and the metastatic tumor biopsy was available for review, the highest grade (WHO 1973 classification) was assigned, and presence of aberrant histology in one of the tissue samples was considered as positive.

### Whole-genome sequencing and analysis

#### Whole-genome DNA sequencing, alignment and data processing

Sufficient amount of DNA (50-200 ng) was extracted from fresh-frozen tumor tissue and blood samples following standard protocols from Qiagen. DNA was fragmented by sonication for NGS Truseq library preparation and sequenced paired-end reads of 2x150 bases with the Illumina HiSeqX platform. Alignment, somatic alterations, ploidy, sample purity and copy numbers estimations were performed as previously described [7]. WGS was aligned to the human reference genome GRCH37 with BWA-mem v.0.7.5a [9], and duplicate reads were marked for filtering. Indels were realigned using GATK IndelRealigner v3.4.46 [10].

Recalibration of base qualities for single nucleotide variants (SNVs) and small insertions and deletions (Indels) was performed with GATK BQSR [11], and SNV and Indel variants were evaluated with Strelka v.1.0.14 [12] using matched blood WGS as normal reference (Table S1.5). Somatic mutations were further annotated with Ensembl Variant Effect Predictor (VEP, v99, cache 99_GRCh37) [13] using GENCODE v33 in combinations with the dbNSFP plugin v3.5 hg19 [14] for gnomAD [15] population frequencies. SNVs, Indels and multiple nucleotide variants (MNVs) variants were removed if the following filters were not passed: default Strelka filters (PASS-only), gnomAD exome (ALL) allele frequency < 0.001, gnomAD genome (ALL) < 0.005 and number of reads < 3. In addition, structural variants (SVs) and copy number changes were estimated using GRIDSS, PURPLE and LINX suit v2.25 [16]. SVs that passed the default QC filters (PASS-only) and Tumor Allele Frequency (TAF) ≥ 0.1 were annotated as “somatic SVs” if there was overlap with coding region. Mean read coverages of tumor and reference samples were estimated using Picard Tools v1.141 (CollectWgsMetrics) based on GRCh37 (https://broadinstitute.github.io/picard/). Genomic and coding tumor mutational burden (TMB; mutations per megabase pair (Mbp)) were calculated considering SNVs, Indels and MNVs (Table S1.5). The total number of somatic mutations in coding region was divided by 28.71 Mbp (protein-coding region size) and in the whole genome by 2,858.67 Mbp (genomic alignment size).

#### Detection of driver genes using dN/dS ratios

Cancer driver genes under strong positive selection were detected with dNdScv v0.0.0.9 [17]. This method uses 192 mutation rates representing all combinations in trinucleotide context. Mutation rates of each gene were corrected by the global mutation rate. The ratio of non-synonymous over synonymous mutations was calculated with maximum-likelihood methods, and statistical significance was estimated. Genes with either qglobal_cv ≤ 0.05 or qallsubs_cv ≤ 0.05 were considered drivers of mUC (Table S1.6 and 1.7).

#### Detection and characterization of recurrent copy number alterations

Ploidy and copy number alterations (CNAs) were estimated as described by Priestley *et al.* 2019; and following the pipeline described by van Dessel *et al.*, 2019. Recurrent focal and broad CNAs were estimated with GISTIC2.0 v2.0.23 [19]. CNAs were classified as shallow or deep according to the threshold in GISTIC2 calls.

Significant recurrent focal CNAs were identified when q ≤ 0.05 and annotated with genes overlapping these regions, which were considered drivers (Table S1.7 and 1.8).

#### APOBEC enrichment and mutagenesis

For each sample, the total number of C>T or C>G (G>A or G>C) mutations was calculated (C*^mut (C>T,C>G)^*). From these mutations, the total number of APOBEC mutations was estimated by counting all mutations in TCW (WGA) context (TCW*^Cmut^*), where W = A or T. The total number of TCW (WGA) motifs and total C (G) nucleotides in the hg19 reference genome were also estimated (TCW*^context^* and C*^context^*, respectively). Using this information and following Roberts *et al.*, 2013, a contingency table was constructed; one-sided Fisher’s exact test was applied to calculate the overrepresentation of APOBEC mutations. P-values were Benjamini-Hochberg corrected. Tumors with adjusted p-values lower than 0.01 were considered APOBEC enriched.

The magnitude of APOBEC enrichment *E* was estimated as [20]

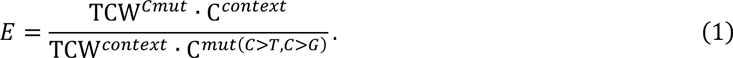

APOBEC enriched tumors (always *E* > 1) were classified as high APOBEC mutagenesis when *E* ≥ 2, and as low APOBEC mutagenesis when *E* < 2. Tumors without APOBEC enrichment were considered tumors with no APOBEC mutagenesis (Table S1.9).

It has been shown that mutations caused by APOBEC3A and APOBEC3B are distinguishable at the tetra- nucleotide context [21]. Mutations in the YTCA (Y = T or C) context have been related to APOBEC3A, while mutations in the RTCA (R = G or A) context are attributed to APOBEC3B. Fold enrichment of C>T and C>G mutations in the tetra YTCA and RTCA context was calculated with equation (1), using the corresponding tetra- nucleotide context (Table S1.9).

#### Clonality fraction estimation

Mutations start as a single copy in the DNA, and multiple copies of the mutated nucleotide may appear if affected by CNAs events. Correcting for tumor purity and CNA, the number of copies *nSNV* of each SNV was calculated as follows [22]

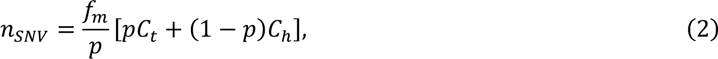

where *fm* is the relative frequency of the mutant variant reads, *p* is the tumor purity, *Ct* is the local copy number affecting a particular SNV, and *Ch* is the healthy copy number (two for autosomes and one for allosomes).

Equation (2) is equivalent to the cancer cell fraction (CCF) with *nSNV* ≍ 1 in haploid and heterozygous-diploid regions; i.e., the fraction of tumor cells carrying a particular mutation. For regions with CNAs, *nSNV* > 1, we must estimate the fraction of cancer cells carrying a particular SNV. As described previously [23], we assume that all SNVs are present in the major copy number *CM*; hence *nSNV* ≤ *CM* will include mutations that were acquired after copy number change events or present only in the minor copy number. Given the number of reference and mutant reads, and assuming binomial distribution, we estimated the expected number of allelic copies (*nchr*) carrying the observed SNV resulting from *fm* values when the mutation is present in 1, 2, 3,…, *Nchr* allelic copies. In some cases (sequencing noise) *nSNV* > *CM*, which was corrected with *Nchr* = *max*(*CM, nSNV*). We also corrected each *fm* value with normal cell contamination – multiplying it by *p*. The resulting estimated *nchr* with the maximum likelihood serves to calculate the CCF as *nSNV*/*nchr*.

Dirichlet process from DPClust v2.2.8 (https://github.com/Wedge-lab/dpclust) with 250 iterations and 125 burn in iterations was applied to the CCF distribution to estimate the fraction of clonal and subclonal SNVs per tumor. Multiple distributions (clusters) were obtained, representing different cancer cell populations. The mean of the distributions was used to classify clusters of SNVs as clonal or subclonal. Clusters of SNVs with mean distribution > 0.8 were considered clonal (Table S1.5).

#### Mutational signatures and genomic subtypes

The mutational pattern of each sample was established by categorizing SNVs according to their 96- trinucleotide context. The contribution of each of the 67 mutational signatures from COSMIC v3 (as deposited on May 2019) [24] was subsequently estimated with MutationalPatterns v1.4.2 (Table S1.10) [25]. To reduce the noise attributed to mutational signatures with very low contribution, mutational signatures were grouped into 26 proposed etiology categories (Table S1.11) derived from Alexandrov *et al.,* 2020, Petljak *et al.,* 2019, Angus *et al.,* 2019 and Christensen *et al.,* 2019. All 26 proposed etiology contributions were used, and hierarchical clustering was applied on 1-Pearson’s correlation coefficient, 80% resampling and 1,000 iterations using ConsensusClusterPlus v1.48.0 [30]. Considering average stability of each cluster and the cluster size (favoring large clusters) after each partition, samples were grouped into five distinct clusters.

Independently, mutational patterns were deconvoluted to estimate *de novo* mutational signatures. Non-negative Matrix Factorization from the NMF R package v0.21.0 was used with 1000 iterations [31]. Evaluating different metrics provided by the NMF R package (high cophenetic correlation coefficient, high dispersion coefficient, high silhouette consensus, high sparseness basis and low sparseness coefficients), seven *de novo* signatures were recovered from the mutational patterns. Cosine similarity was applied to compare the *de novo* signatures with mutational signatures from COSMIC v3.

#### Detection of chromothripsis

Genomic catastrophic-like events, such as chromothripsis, were detected with Shatterseek v0.4 [32] using default parameters. Absolute copy numbers (as derived by PURPLE) were rounded to the nearest integer; only structural variants with TAF ≥ 0.1 at either end of the breakpoint were considered, and chrY was excluded.

Applying the filters suggested by Cortés-Ciriano, et al. [32], we identified 220 chromothripsis events suggesting an enrichment in mUC compared to primary UC from the PCAWG dataset [31] (76% vs 48%, p = 0.011).

Inspecting manually these events, we concluded that a more stringent filter should be used to reduce the rate of false positive events. The following filters were applied and chromothripsis was considered when: a) the number of intra-chromosomal structural variants ≥ 25; b) the maximum number of oscillating CN segments with two states ≥ 7 or with three states ≥ 14; c) the size of the chromothripsis event ≥ 20 Mbp; d) random distribution of breakpoints *p* ≤ 0.05; and e) *chromosomal breakpoint enrichment p* ≤ 0.05. All 220 chromothripsis events are summarized in Table S1.12, indicating those detected by the stringent filtering, which we refer to as chromothripsis throughout the text.

#### MicroSatellite Instabilty (MSI) status

As previously described [7], MSI status was determined by estimating the MSI score as the number of indels (length < 50 bp) per Mbp occurring in homopolymers of five or more bases, dinucleotide, trinucleotide and tetranucleotide sequences of repeat count above five. Tumors with MSI score > 4 were considered MSI positive (Table S1.13).

#### Detection of homologous recombination (HR) deficiency

The Classifier for Homologues Recombination Deficiency (CHORD; v2.0) with default parameters was used to identify tumors with HR proficiency and deficiency [33]. Four samples had very high number of indels corresponding with MSI samples and were discarded for the HR deficiency analysis (Table S1.14).

#### Detection of kataegis

Following the method described by van Dessel *et al.* 2019, kataegis events were estimated using all SNVs. Each chromosome was divided into segments (maximum 5000 segments) of five or more consecutive SNVs.

Segments were considered a kataegis event when the mean intermutational distance was ≤ 2000 bp (Table S1.15). Events were considered APOBEC-driven when >60% of mutations were C>T or C>G mutations in TCW context.

#### Mutational load across genomic regions

The genome was divided in regions (bins) of one Mbp size. The number of SNVs was counted in each bin, and the mean number of SNVs was estimated from the entire cohort. These values represented the average SNVs/Mbp reflecting the mutational load in each genomic region. The average SNVs/Mbp was smoothed by applying a moving average with k = 9 bins. This approach was used per sample and for mean values from the entire cohort.

#### Genomic alteration of oncogenic pathways

Eleven oncogenic pathways were analyzed for somatic alterations. The list of genes was modified from Sanchez-Vega *et al.,* 2018 and Leonard, 2001 (Table S1.16). Altered pathways were defined when at least one of the pathway-genes was affected by any somatic mutation (SNV, Indel, MNV, SV or deep CNA; excluding synonymous mutations).

#### Inventory of clinically-actionable somatic alterations and putative therapeutic targets

Current clinical relevance of somatic alterations in relation to putative treatment options or resistance mechanisms and trial eligibility was determined based upon the following databases: CiViC [36] (Nov. 2018), OncoKB [37] (Nov. 2018), CGI [38] (Nov. 2018) and the iClusion (Dutch) clinical trial database (Sept. 2019, Rotterdam, the Netherlands). The databases were aggregated and harmonized using the HMF knowledgebase- importer (v1.7; https://github.com/hartwigmedical/hmftools/tree/master/knowledgebase-importer).

Subsequently, we curated the linked putative treatments and selected treatments for which level A (biomarker for approved therapy or in guidelines) or level B (biomarker on strong biological evidence or used in clinical trials) evidence was available. Genomic alterations that confer resistance to certain therapies were excluded from the analysis. Treatment strategies including anti-hormonal therapy (as used for breast and prostate cancer), surgical resection, or radioiodine uptake therapy were excluded. Furthermore, closed trials (according to www.clinicaltrials.gov), and trials with only pediatric patients or patients with hematological malignancies were excluded (Table S1.17). The data base was complemented with *FGFR3* and *NTRK2* gene fusions (at RNA level) and patients with MSI high and HR deficient tumors. On-label treatments included chemotherapy (cisplatin, gemcitabine, doxorubicin, mitomycin, and valrubicin) and the FGFR3 inhibitor erdafitinib. Off-label treatments included treatments that are on-label for other tumor types (FDA approved drugs according to the US national cancer institute; https://www.cancer.gov/about-cancer/treatment/drugs/cancer-type), and treatments available in clinical trials or basket trials. When patients had more than one possible treatment, on-label treatment was the preferred treatment, followed by on-label treatments for other tumor types.

#### DNA accessibility estimation (ChIPseq)

All available ChIPseq data for healthy urinary bladder (H3K4me1, H3K4me3, H3K36me3 and H3K27ac) were downloaded from the ENCODE portal (https://www.encodeproject.org) to our local server. The *bed.gz* files were imported with *narrowPeak* format for analysis. The signal of each experiment was divided in regions of one Mbp, and a moving average with k = 9 bins was applied. The scale of the signal was normalized; hence the sum of all regions in a chromosome is one. This step was taken to compensate for the bias observed in peak intensity signals across different chromosomes, possible due to technical issues in the ChIPseq technology, e.g. hyper-ChIPable regions or mappability [39].

High DNA accessible regions (open chromatin) were determined as such if the ChIPseq signal value of the region was above the median. Otherwise, the region was considered as low DNA accessible (condensed chromatin). This procedure was applied on each chromosome.

### Whole-transcriptome sequencing and analysis

#### RNA-sequencing, alignment and data pre-processing

Total RNA was extracted using the QIAGEN QIAsymphony kit (Qiagen, FRITSCH GmbH, Idar-Oberstein, Germany). Samples with a minimum of 100 ng total RNA were sequenced according to the manufacturer’s protocols. Paired-end sequencing of RNA was performed on the Illumina NextSeq 550 platform (2x75bp) and Illumina NovaSeq 6000 platform (2x150bp).

Prior to alignment, samples were visually inspected with FastQC v0.11.5. Sequence adapters (Illumina TruSeq) were trimmed using Trimmomatic v0.39 [40] at the following settings: ILLUMINACLIP:adapters.fa:2:30:10:2:keepBothReads MINLEN:36. The trimmed paired-end reads were aligned to the human reference (GRCh37) using STAR v2.7.1a [41] with genomic annotations from GENCODE hg19 release 30 [42]. Multiple lanes and runs per sample were aligned simultaneously and given respective read- group identifiers for use in downstream analysis to produce two BAM files per sample, consisting of genome- and transcriptome-aligned reads respectively.

STAR was performed using the following command: STAR --genomeDir <genome> --readFilesIn <R1> <R2> --readFilesCommand zcat --outFileNamePrefix <outPrefix> --outSAMtype BAM SortedByCoordinate --outSAMunmapped Within --chimSegmentMin 12 -- chimJunctionOverhangMin 12 --chimOutType WithinBAM --twopassMode Basic --twopass1readsN -1 -- runThreadN 10 --limitBAMsortRAM 10000000000 --quantMode TranscriptomeSAM --outSAMattrRGline <RG> After alignment, duplicate reads were marked and alignment quality metrics (flagstat) were generated using Sambamba v0.7.1 [43]. For each genome-aligned sample, the uniformity of read distributions across transcript lengths was assessed using tin.py v2.6.6 [44] from the RseQC library v3.0.0 [45].

FeatureCounts v1.6.3 [46] was applied to count the number of overlapping reads per gene using genomic annotations from GENCODE (hg19) release 30 [42]; only primary (uniquely mapped) reads were counted per exon and summarized per gene: featureCounts -T 50 -t exon -g gene_id --primary -p -s 2 -a <gencode> -o <output> <genomic BAMs> RSEM v1.3.1 [47] was applied to quantify RNA expression into transcripts per million (TPM) values using transcript annotations from GENCODE (hg19) release 30 [42]: rsem-calculate-expression --bam --paired-end --strand-specific --alignments -p 8 <transcriptome BAM> <RSEM Index> <output>

#### Transcriptome expression data mapped to genomic regions

MultiBamSummary from deepTools v1.30.0 [48] was used to read BAM files and estimate number of reads in genomic regions with a size of one Mbp. The average raw read count per Mbp was calculated, and a moving average with k = 9 bins was applied. The scale of the read counts was normalized following the method for DNA accessibility regions, and high transcriptional regions were defined as such when the expression value of one region was above the median. This procedure was applied on each chromosome.

#### Transcriptomic subtypes: clustering samples by RNA-seq data

Several methods have been proposed to classify bladder cancer into transcriptomic subtypes. In an attempt to standardize the molecular profiling of bladder cancer, a consensus molecular classification was proposed for MIBC based on RNA-seq data from 1750 patients [5]. This classifier was developed strictly for MIBC and is not directly applicable to mUC [49]. Furthermore, this classifier was developed for samples derived from the same organ carrying transcriptomic contamination of normal urothelial cells. In this study, biopsies were obtained from metastatic sites leading to contamination with normal cells from multiple different organs for which no correction was applied in the consensus classifier. Therefore, it was mandatory to perform *de novo* subtyping in this study, which is described below.

Multiple methods were explored to correct for the bias of biopsy site, including batch-correction with DESeq2 [50], and a tissue-aware correction method developed by the Genotype-Tissue Expression (GTEx) project [51]. In both cases, transcripts from liver tissue were very dominant and clustered together in one stable cluster.

The tissue-specific transcript removal method described above was successfully able to correct for organ-specific transcripts, and as a result samples were clustered based on transcriptomic features rather than biopsy site.

Transcripts were normalized using DESeq2 v1.24.0 [50] with variance stabilizing transformation. Only highly expressed mRNA with base mean above 100 was kept. The top 50% most variably expressed genes (6,410 transcripts) were used for clustering. To reduce the ‘transcriptomic noise’ introduced by normal cells of the tissue from which the biopsy was taken, these transcripts were identified and excluded. Samples were grouped according to their biopsy site: liver (n = 31), lymph node (n = 30), bone (n = 5), other (n = 23) and unknown (n=1). Differential expression analysis was performed to compare tumors from a specific biopsy site (liver, lymph node and bone) against all other tumors using DESeq2 with Wald test p-value estimation. Tissue-specific transcripts with log2 Fold Change (log2FC) > 1.0 and Benjamini-Hochberg corrected p-value < 0.05 were considered differentially expressed and identified as tissue-specific (Table S1.18). A total of 689 transcripts were tissue-specific, and were removed from the data set.

The remaining 5,721 transcripts were grouped by hierarchical clustering with 1-Pearson’s correlation coefficient, 80% resampling and 1,000 iterations using ConsensusClusterPlus v1.48.0 [30]. The mean cluster consensus value was obtained as a measure of cluster stability. Increasing the number of clusters will increase the stability by creating smaller clusters. Taking this into account, the criteria for selecting five clusters was based on cluster stability and cluster size by not allowing clusters with <5 samples (Table S1.19). Patients with primary upper tract tumors did not cluster together as was observed for biopsy sites, instead they were distributed across all different transcriptomic clusters (Fig. 5), suggesting that their influence on the clustering was negligible.

To identify transcripts that contribute most to each cluster, we followed the same strategy used to identify tissue-specific transcripts. The top five transcripts with the highest log2FC and with Benjamini-Hochberg adjusted p-values lower than 1x10^-5^ were identified as the most overexpressed genes per cluster. Other differentially expressed genes were included for their clinical relevance (*TGFB3*, *DDR2*, *PDGFRA*, *CD274* and *TGFBR1*). All differentially expressed genes per cluster with adjusted p < 1x10^-5^ and log2FC > 1 are listed in Table S1.20.

To compare our classification system developed for mUC with the consensus classifier, all samples were classified into one of six molecular classes identified in MIBC. All normalized transcripts (excluding biopsy specific transcripts) were used as input for the consensus classifier of primary MIBC (v1.1.0) [5].

#### Phenotypic markers and signature score

Marker genes for basal (*CD44, CDH3, KRT1, KRT14, KRT16, KRT5, KRT6A, KRT6B, KRT6C*), squamous (*DSC1, DSC2, DSC3, DSG1, DSG2, DSG3, S100A7, S100A8*), luminal (*CYP2J2, ERBB2, ERBB3, FGFR3, FOXA1, GATA3, GPX2, KRT18, KRT19, KRT20, KRT7, KRT8, PPARG, XBP1, UPK1A, UPK2*), neuroendocrine (*CHGA, CHGB, SCG2, ENO2, SYP, NCAM1*), cancer-stem cell (*CD44, KRT5, RPSA, ALDH1A1*), EMT (*ZEB1, ZEB2, VIM, SNAI1, TWIST1, FOXC2, CDH2*) and claudin (*CLDN3, CLDN7, CLDN4, CDH1, SNAI2, VIM, TWIST1, ZEB1, ZEB2*) were used for signature scores [3]. Stroma (*FAP*), interferon, and CD8+ effector T cell (*IFNG, CXCL9, CD8A, GZMA, GZMB, CXCL10, PRF1, TBX21*) markers were also included [52]. All normalized expression values were median centered, and the mean expression of each group of genes was defined as signature score.

#### Pathway activity score

Transcriptionally activated genes by the eleven canonical pathways analyzed in this study were used to estimate pathway activity score. All normalized expression values were median centered, and the mean expression of each group of genes was defined as activity score. Activity score was estimated for the TGFβ pathway (*ACTA2, ACTG2, ADAM12, ADAM19, CNN1, COL4A1, CCN2, CTPS1, RFLNB, FSTL3, HSPB1, IGFBP3, PXDC1, SEMA7A, SH3PXD2A, TAGLN, TGFBI, TNS1, TPM1*) [53], cell cycle pathway (*MKI67, CCNE1, BUB1, BUB1B, CCNB2, CDC25C, CDK2, MCM4, MCM6, MCM2*) [52], WNT pathway (*EFNB3, MYC, TCF12, VEGFA*) [53], Notch pathway (*HES1, HES5, HEY1*) [54], PI3K pathway (*AGRP, BCL2L11, BCL6, BNIP3, BTG1, CAT, CAV1, CCND1, CCND2, CCNG2, CDKN1A, CDKN1B, ESR1, FASLG, FBXO32, GADD45A, INSR, MXI1, NOS3, PCK1, POMC, PPARGC1A, PRDX3, RBL2, SOD2, TNFSF10*) [55], hippo pathway (*TAZ, YAP1*) [56], p53 pathway (*CDKN1A, RRM2B, GDF15, SUSD6, BTG2, DDB2, GADD45A, PLK3, TIGAR, RPS27L, TNFRSF10B, TRIAP1, ZMAT3, BAX, BLOC1S2, PGF, POLH, PPM1D, PSTPIP2, SULF2, XPC*) [57], Nrf2 pathway (*GCLM, NQO1, PHGDH, PSAT1, SHMT2*) [58], MYC pathway (*TFAP4, BMP7, CCNB1, CCND2, CCNE1, CDC25A, CDK4, CDT1, E2F1, GATA4, HMGA1, HSP90AA1, JAG2, CDCA7, LDHA, MCL1, NDUFAF2, MTA1, MYCT1, NPM1, ODC1, SPP1, PIN1, PTMA, PRDX3, PRMT5, DNPH1, TFRC, EMP1, PMEL, C1QBP*) [59], RTK-RAS pathway (*SPRY2, SPRY4, ETV4, ETV5, DUSP4, DUSP6, CCND1, EPHA2, EPHA4*) [60] and JAK-STAT pathway (*IRGM, ISG15, GATA3, FCER2, THY1, NFIL3, ARG1, RETNLB, CLEC7A, CHIA, OSM, BCL2L1, CISH, PIM1, SOCS2, GRB10*) [61].

#### Pathway enrichment analysis

All differentially expressed genes with Benjamini-Hochberg adjusted p < 0.05 and absolute log2FC > 1 in each transcriptomic subtype were used for pathway enrichment analysis. Using reactomePA v1.34 [62], the top ten (sorted by Benjamini-Hochberg adjusted p-value) up- and down-regulated pathways were selected.

#### Immune cell infiltration

To quantify immune cell fractions in each sample, we analyzed RSEM read counts of all transcripts with immunedeconv v2.0.3 [63] using the quanTIseq method [64].

#### Detection of gene fusions

In-frame gene fusions were detected at DNA level by the GRIDSS, PURPLE, LINX suite v2.25 [16], and reported relevant if they appear in the ChimerDB 4.0 (Table S1.21) [65]. At RNA level, Arriba v2.0.0 (https://github.com/suhrig/arriba/) was used to infer gene fusion events with the option to discard known false positives from a list provided by Arriba. High confidence fusions were retained, and only events where at least one transcript is protein coding were kept (Table S1.22). Gene fusions previously identified by other studies, mostly from the TCGA data, were identified with ChimerDB 4.0 [65]. All “deletion/read-through” events were discarded as possible false positives unless they were supported by the ChimerDB 4.0 database. Medium confidence fusions were included in the final list if one of the fused genes appeared in a high confidence fusion event.

#### mRNA editing

Jalili*, et al.* 2017 [67] identified hotspot mutations in the mRNA of *DDOST* and *CYFIP1* that are targeted by APOBEC3A. The genomic position of these hotspot mutations reveals hairpin loop structures that are the ideal substrate for APOBEC3A. Due to the short life-time of mRNA molecules, the presence of these hotspot mutations reflects ongoing APOBEC mutagenesis. The proportion of C>U mutations in chr1:20981977 and chr15:22999350 were estimated to identify the RNA-editing activity of APOEBC3A.

#### APOBEC mutation rate and APOBEC expression in tumors with multiple sequential biopsies

A second metastatic tumor biopsy was taken in eight patients from the same (n = 5) or a different (n = 3) metastatic lesion, and analyzed by WGS (n = 8) and RNA-seq (n = 7). Patients with high APOBEC mutagenesis (n = 5) tumors were all treated with pembrolizumab (one patient received consecutive lines of chemotherapy and pembrolizumab). Two out of three patients with low APOBEC mutagenesis tumors were treated with chemotherapy. All patients, except two, received systemic pre-treatment.

Each patient’s first and second biopsies shared a high proportion of mutations (SNVs, Indels and MNVs), confirming the clonal relation of the sampled sites. Dirichlet process from the DPClust v2.2.8 R package (https://github.com/Wedge-lab/dpclust) with 250 iterations and 125 burn in iterations was applied to the CCF distribution of paired-biopsies to estimate the subclonal (clusters) composition of each tumor. All unique mutations in each biopsy were considered a subclone; only subclones with >5% of SNVs were considered relevant. Small populations of subclones (<5% of SNVs) were merged to the nearest subclone. The evolutionary tree was reconstructed following the *sum rule* [66].

The CCF of somatic mutations in the branches was lower than that in the trunk, suggesting that these mutations are recently acquired mutations. To compare APOBEC mutagenesis between patients, the rate of novel APOBEC associated mutations was estimated. Only unique SNVs from the second biopsy were kept, as these somatic alterations probably correspond to new mutations acquired during the time frame between the biopsies. As the time elapsed between the first and the second biopsy varied between tumors, we normalized the number of recent APOBEC associated mutations by dividing the total over the number of days elapsed between the biopsies. The value estimated is proportional to the mutation rate of APOBEC associated mutations (mutations per day).

For seven tumors, RNA-seq data was available. Expression of *APOBEC3A* and *APOBEC3B* per patient represents the mean normalized expression of the paired biopsies.

#### Analysis of the *The Cancer Genome Atlas* primary bladder cancer cohort

To compare the genomic and transcriptomic landscapes of mUC with primary bladder cancer, publicly available data of the TCGA bladder cancer cohort, including somatic mutations detected by Mutect (SNVs and Indels) of 412 tumors, GISTIC copy number changes at gene level of 410 tumors, and RNA-seq (HTSeq counts; Affymetrix SNP6 arrays) data available for 410 tumors were analyzed. Some samples had very few mutations, and only tumors with total SNVs > 50 were considered in this analysis (367/412). The same method applied on our mUC cohort was applied on the TCGA data to deconvolute mutational signatures and to identify genomic subtypes. Twelve genomic subtypes were identified, but several of them formed small groups with very specific mutational signature patterns, including one sample with very high POLE signature. All genomic subtypes with < 1% of the total cohort were grouped together in GenS0. We compared the genomic subtypes between mUC and the TCGA cohort using cosine similarity.

Transcript counts were normalized with DESeq2 [50] following the same procedure used for the mUC cohort. All tumors were from primary UC, and organ-specific transcripts were not discarded. The consensus classifier of primary MIBC [5] was applied to infer the transcriptomic subtype of each tumor.

### Code availability

All custom code and scripts are available at https://bitbucket.org/ccbc/dr31_hmf_muc/ and https://github.com/hartwigmedical/.

### Statistical analysis

Several statistical tests were used in this study: Fisher’s exact test, T-test, binomial test, Wald test for differential expression analysis and logistic regression analysis, Wilcoxon signed-rank test, Wilcoxon rank-sum test, Kruskal-Wallis test, and tests performed by dNdScv [17] and GISTIC2 [19]. In cases of multiple testing, p-values were Benjamini-Hochberg corrected. The appropriate statistical test is mentioned in the text when describing significance values. All statistical analyses were performed using the statistical computing and graphics platform R v3.6.1 [69].

## Results

### Genomic landscape of mUC

Analysis of WGS (mean coverage 106 X) and matched blood samples (mean coverage 38 X) identified a median of 20,634 SNVs, 1,018 Indels and 175 MNVs (Fig. S2a). SNVs were more frequent in coding regions (7.63 SNVs per Megabase pair; SNVs/Mbp) than in the whole genome (7.22 SNVs/Mbp; Wilcoxon signed-rank test p = 0.0024; Supplementary Fig. 2b). However, Indels and MNVs were less frequent in coding regions (Wilcoxon signed-rank test p < 0.001 and p = 0.0072, respectively). Analysis of all SNVs revealed that 68% of all SNVs were clonal with a median of 74% per tumor, and that 91% of the tumors were enriched for APOBEC associated mutations (73% high and 18% low enrichment of APOBEC mutagenesis; Fig. 2a). The mean contribution of APOBEC COSMIC signatures (SBS2 and SBS13) in tumors with high APOBEC mutagenesis enrichment was 52% *versus* 15% in tumors with low APOBEC mutagenesis.

**Figure 2.**
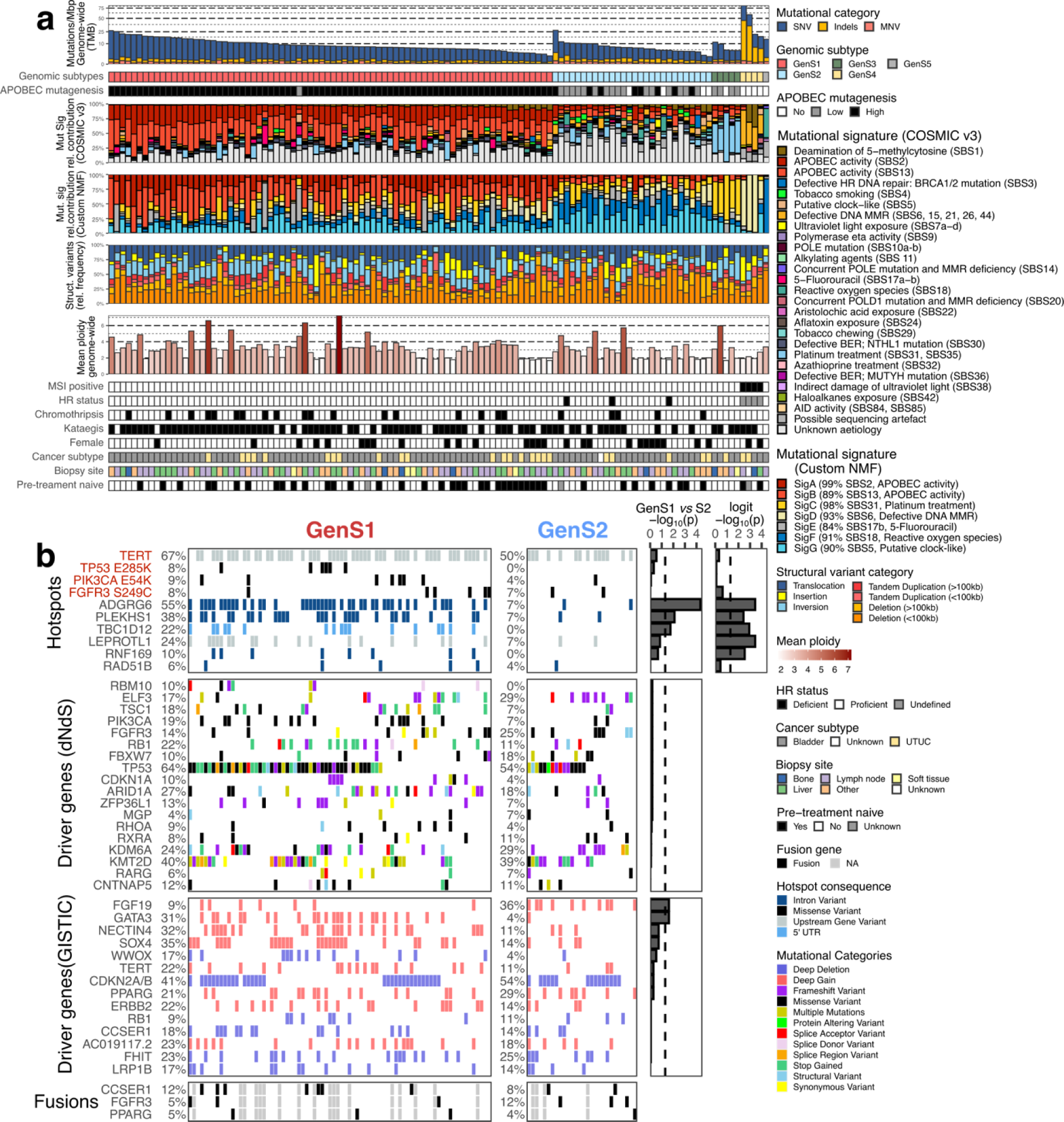
Genomic landscape of metastatic urothelial carcinoma stratified by genomic subtypes. a) Whole-genome sequencing data from biopsy samples of metastatic urothelial carcinoma were classified into genomic subtypes by hierarchical consensus clustering of the relative contribution of COSMIC v3 mutational signatures [24] grouped by etiology. The genomic features are displayed from top to bottom as follows: Genome-wide TMB; Genomic subtype (GenS1-5); APOBEC enrichment analysis showing tumors with no-, low- and high-APOBEC mutagenesis; Mutational signatures grouped by etiology, except for APOBEC activity for which both signatures are shown separately; Relative contribution of seven *de novo* (custom) mutational signatures by deconvolution of SNVs in 96 tri-nucleotide context with NMF; Relative frequency of different types of structural variants; Mean ploidy; Tumors with MSI; HR deficiency status; Samples with at least one chromothripsis event; Samples with at least one kataegis event; Female patients; Origin of primary tumor; Metastatic site from which a biopsy was obtained; Treatment-naïve patients. b)Overview of recurrent hotspot mutations, driver genes and gene fusions for the genomic subtypes GenS1 and GenS2. Name of genes affected by hotspot mutations in >5% of samples are displayed in red when the hotspot had a COSMIC id. Significantly mutated genes were estimated by dNdScv [17]; all genes with q < 0.05 were considered driver genes. Recurrent focal copy number changes were estimated by GISTIC2 [19]; genes in genomic regions with q < 0.05 were considered significant. Only affected genes present in >10% of the samples are shown. Gene fusions were detected from RNA-seq data. Benjamini-Hochberg adjusted p-values of Fisher’s exact test (for hotspot mutations and GISTIC2) and of logistic regression analysis corrected by mutational load (driver genes by dNdScv) are shown on the right to reflect the significance of the difference between GenS1 and GenS2. An additional logistic (logit) regression analysis was performed on hotspot mutations to show the linear relation with the number of APOBEC associated mutations. Bars beyond the dashed line (-log10(0.05)) are statistically significant. Abbreviations: TMB = tumor mutational burden; Mbp = mutations per mega base pair; NMF = non-Negative Matrix Factorization; MSI = microsatellite instability; HR = Homologous Recombination; UTUC = upper tract urothelial carcinoma.

Genes harboring more mutations in their coding sequence than expected by random chance were analyzed with dNdScv; the analysis revealed 18 significantly mutated genes (Table S1.6). These genes resembled those reported for primary UC [3], although mUC lesions did harbor more somatic mutations in *TP53* than numbers reported in TCGA (non-synonymous mutations and indels; 60% *vs* 49%, Fisher’s exact test p = 0.021, Fig. S3).

SVs were common with a median of 259 (40,297 in total) per tumor. Deletion was the most frequent type of SV with a median of 92 per tumor (Fig. S2d). The genes most frequently affected by SVs were *CCSER1* (13%) and *AHR* (12%; Fig. S3). Chromothripsis, a complex event that produces SVs in which chromosomes are shattered and rearranged, was detected in 20% of the tumors (Fig. S4).

Chromosomal arm and focal CNA were analyzed with GISTIC2. This revealed frequent deletion of chromosome 9 and amplification of chromosome 20 (Fig. S5a). In total, 49 genomic regions were significantly altered by focal CNAs which included several oncogenic genes (Table S1.8). The most frequently amplified genes were *SOX4* (28%), *GATA3* (22%), *PPARG* (22%), and ERBB2 (19%); the most frequently deleted genes were *CDKN2A/B* (43%), *FHIT* (24%), *CCSER1* (17%) and *LRP1B* (17%), and also resembled those reported in primary UC [3] (Fig. S3).

Hotspot mutations in driver genes concerned *FGFR3* S249C (8%), *PIK3CA* E54K (7%), and *TP53* E285K (5%). Hotspot mutations in the *TERT* promoter were present in 62% of the tumors (Fig. S3; Table S1.23). Still, *TERT* expression did not differ between tumors with and without hotspot mutations (Fig. S6b), in line with a previous report [68]. However, differential gene expression analysis showed that tumors with hotspot mutations in the *TERT* promoter had downregulation of genes related to the muscle contraction pathway (Fig. S6a, Table S1.24). Furthermore, hotspot mutations were identified in non-coding regions of *ADGRG6 (40%)*, *PLEKHS1 (28%)*, *LEPROTL1* (18%), and *TBC1D12* (15%; Fig. S3) with no apparent association with gene expression and minimal transcriptomic effect (Fig. S6a-b). The hotspot areas of *ADGRG6*, *PLEKHS1* and *TBC1D12* form hairpin loop structures in the DNA with specific tri-nucleotide sequences frequently mutated by APOBEC enzymes (Fig. S6c). Unlike other known driver genes affected by hotspot mutations (*TERT*, *FGFR3*, *PIK3CA* and *TP53*), these genes were not significantly affected by other somatic mutations in the coding region or by CNAs, suggesting that hotspot mutations in *ADGRG6*, *PLEKHS1* and *TBC1D12* are likely passenger hotspots attributed to APOBEC activity as theoretically predicted [69].

Gene fusion analysis performed at the transcriptomic level (Table S1.22) detected 1394 gene fusions, of which 10% were also reported in the TCGA cohort [65]. Seventy-six percent of all individual genes found involved in fusion events have previously been implicated in fusions [65]. *FGFR3* gene fusions were present in seven out of 90 samples with only one *FGFR3*-*TACC3* fusion. Four *PPARG* fusions were detected, of which two *PPARG-TSEN2* fusions were confirmed at DNA level (Table S1.21). Other putative fusion events in cancer-related genes were found in *CCSER1* (n = 9), *ERBB4* (n = 5), *RB1* (n = 4), *MDM2* (n = 4), *TERT* (n = 3) and *STAG2* (n = 3).

A stratification based on the proposed etiology of SNV COSMIC signatures using unsupervised consensus clustering [30] revealed two major genomic subtypes (GenS; Fig. 2; Fig. S7). GenS1 (67%) contained almost exclusively tumors with high APOBEC mutagenesis, which was reflected by a large contribution of APOBEC signatures SBS2 and SBS13. In addition, we performed deconvolution of SNV patterns by non-negative matrix factorization (NMF; Fig. S8), which confirmed APOBEC signatures as the main source of mutations in GenS1, with high contribution of *de novo* mutational signatures SigA (0.99 cosine similarity with APOBEC signature SBS2) and SigB (0.89 cosine similarity with APOBEC signature SBS13). GenS2 (24%) aggregated predominantly tumors with low APOBEC mutagenesis (16 out of 28), and was characterized by signatures of unknown etiology. *De novo* mutational signatures SigF (0.91 cosine similarity with SBS18 COSMIC signature) and SigG (0.90 cosine similarity with SBS5 COSMIC signature) were dominant in GenS2. Analysis of the TCGA cohort (WES data) showed that GenS1 and GenS2 were also the two major genomic subtypes in localized UC (Fig. S9a- b). The other three smaller subtypes (9% of the present cohort) were related to the platinum treatment signature (GenS3), the defective DNA mismatch repair (MMR) signature and microsatellite instability (MSI, GenS4), and the reactive oxygen species signature (GenS5), which was characterized by *de novo* sigF in > 95%.

The origin of somatic driver mutations was independent of the genomic subtypes, although amplifications of *GATA3* and *FGF19* were enriched in GenS1 and GenS2, respectively (Fig. 2b). Hotspot mutations occurred more frequently in GenS1. In particular, *ADGRG6*, *PLEKHS1* and *TBC1D12* were significantly more often mutated in GenS1. However, these hotspot mutations are potentially irrelevant byproducts caused by APOBEC mutagenesis as logistic regression analysis showed a correlation between APOBEC mutational load (C>T and C>G mutations in TCW context) and occurrence of these hotspot mutations (Fig. 2b).

Other genomic differences between GenS1 and GenS2 (Fig. S3, S10 and S11) included higher SNVs/Mbp in GenS1 and higher Indels/Mbp in GenS2, which was also observed in the TCGA cohort (Fig. S9c). All three tumors with homologous recombination (HR) deficiency identified were of subtype GenS2. Clinical characteristics such as sex, cancer subtype (bladder or upper tract UC), and pre-treatment status did not differ between GenS1 and GenS2. Thus, despite that two very different etiologies lead to UC development, the two mutagenic processes lead to similar profiles of somatically affected driver genes.

### APOBEC mutagenesis is an active process that generates new mutations in mUC

In tumors with high APOBEC mutagenesis, the mean ploidy and the number of genes affected by CNA were higher than in tumors without APOBEC mutagenesis (Wilcoxon rank-sum test p = 0.01 and p < 0.001, respectively; Fig. S12e-f). This phenomenon may indicate genomic instability in APOBEC-driven mUC tumors.

APOBEC enzyme expression analysis revealed neither significant differences between GenS1 and GenS2 (Fig. S12a), nor between tumors with and without APOBEC mutagenesis (Fig. S12c). To further investigate the role of APOBEC mutagenesis in mUC, we analyzed WGS data of eight tumors from patients who had undergone serial biopsies, and reconstructed their evolutionary paths (Fig. 3a). Significantly mutated genes (dNdScv), kataegis events and hotspot mutations displayed in the trunk represent clonal alterations that are fixed in the tumor and are present in all cancer cells of both biopsies. Alternatively, mutations displayed in the branches represent subclonal alterations that are present exclusively in one biopsy, or present in both biopsies but are found only in a fraction of cancer cells (e.g. patient 1 in Fig. 3a). A lower cancer cell fraction in the branches of than the trunk of the evolutionary trees (Fig. 3b) suggests that mutations from branches corresponding to the second biopsy might be novel and not widely spread in the cancer cell population. The rate of novel APOBEC mutations (number of APOBEC mutations divided by the number of days elapsed between serial biopsies) was calculated using only mutations from branches corresponding to the second biopsy. This analysis showed that tumors with high APOBEC mutagenesis accumulated more novel APOBEC mutations than other tumors (Fig. 3d, Wilcoxon rank-sum test p = 0.036). In line with this, we observed that the APOBEC mutational signature was still present in the second biopsy of these patients, together suggesting that APOBEC mutagenesis is ongoing in these samples (Fig. 3c). We further confirmed ongoing APOBEC mutagenesis by analyzing hotspot mutations in mRNA. The frequency of hotspot mutations in *DDOST* (mRNA) was enriched in tumors with high APOBEC mutagenesis and in GenS1 compared to GenS2, with up to 15% of mRNA molecules mutated in one single sample (Fig. S13a).

**Figure 3.**
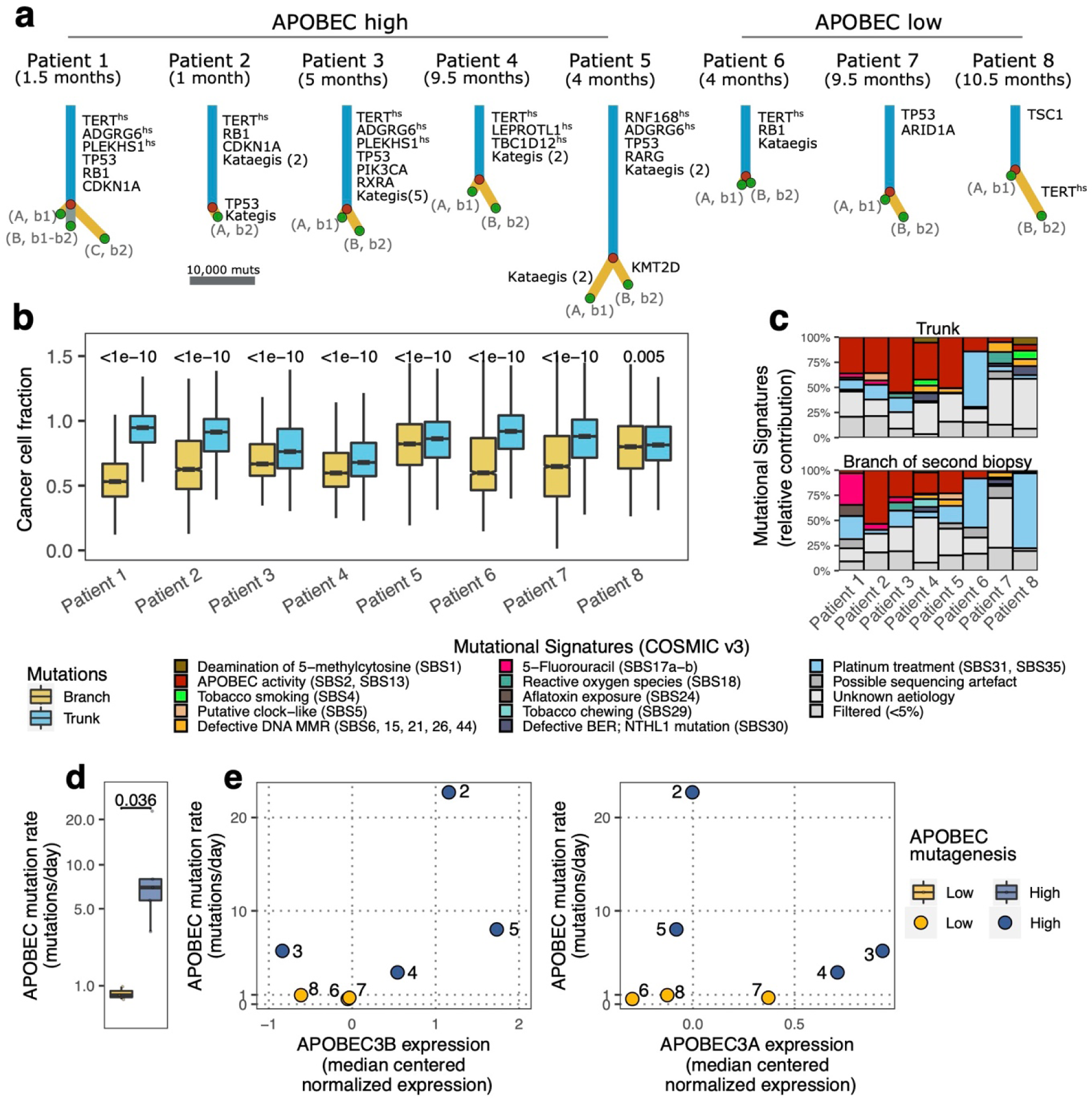
Cancer evolution of eight tumors from metastatic urothelial carcinoma patients with serial biopsies of metastatic lesions. a) Evolutionary trees from eight tumors (five with high APOBEC and three with low APOBEC mutagenesis) from which two biopsies were obtained (interval depicted in months) were reconstructed from single nucleotide variants. Significantly mutated genes (dNdScv), kataegis (number of events in parenthesis) and hotspot sites (hs) are shown, and their locations are indicated (trunk or branch). Branches represent subclonal populations (A, B or C), indicating their presence in the first or second biopsy (b1 or b2). For patient 1, a subclonal population is present in both biopsies. The cancer cell fraction of each single nucleotide variant was calculated and clustered using DPClust for paired-biopsies. The evolutionary tree was reconstructed using the *sum rule* [66]. b) Boxplots comparing the cancer cell fraction of somatic mutations from the trunk and branches. Wilcoxon rank-sum test was applied and p-values were Benjamini-Hochberg corrected. c) COSMIC v3 mutational signatures calculated from the trunk and from the branch exclusive to the second biopsy. d) The APOBEC mutation rate from novel (recent) mutations in the second biopsy was compared between low and high APOBEC mutagenesis tumors. Wilcoxon rank-sum test was applied. e)APOBEC mutation rate is displayed as a function of *APOBEC3A* and *APOBEC3B* median centered normalized expression. APOBEC expression was estimated as the mean expression of both biopsies per tumor. Numbers indicate patient number. RNA-seq was not available for patient 1.

Several studies have linked the expression of APOBEC3A/3B to APOBEC mutagenesis in UC [3, 70]. When comparing the estimated relative expression of APOBEC3A and 3B in the same tumor, we observed differential expression of these enzymes (Fig. 3e). Some tumors had high levels of *APOBEC3A* expression while the expression of *APOBEC3B* was low – or *vice versa*. The correlation between the expression of APOBEC3A and APOBEC3B was poor (Fig S13b), which may explain the lack of differential expression of individual APOBEC enzymes between tumors with different levels of APOBEC mutagenesis (Fig. S12c). To further investigate the link between the occurrence of APOBEC mutations and APOBEC gene expression, we considered the expression of both APOBEC enzymes and calculated an APOBEC score (sum of the median centered normalized expression of *APOBEC3A* and *3B*). It appeared that tumors with high APOBEC mutagenesis had a higher APOBEC score than other tumors (Wilcoxon rank-sum test p = 0.012; Fig. S12d). The APOBEC score was also higher in GenS1 compared to GenS2 (Wilcoxon rank-sum test p < 0.001; Fig. S12b). This analysis confirmed a link between APOBEC gene expression and APOBEC mutations in mUC. To further validate this result, the fold enrichment of C>T and C>G mutations in the tetra YTCA (related to APOBEC3A) and RTCA (related to APOBEC3B) context was calculated for the entire cohort (Fig. S13c). We found that both APOBEC3A and APOBEC3B contribute to APOBEC associated mutations (fold enrichment is above 1.0), but APOBEC3A appears to be the main contributor in mUC, as was reported previously [21, 71].

### APOBEC associated mutations are randomly distributed across the genome in mUC

The substrate of APOBEC enzymes is single-stranded DNA (ssDNA) [20], this has led to the hypothesis that APOBEC enzymes are mainly active during replication or in open chromatin and transcriptionally active genomic regions [72, 73]. As our cohort contained predominantly tumors with APOBEC mutagenesis, and WGS data of these tumors was available, we had the unique opportunity to explore the mutational consequences of APOBEC mutagenesis across the genome.

The total number of SNVs/Mbp varied across the genome, and non-APOBEC mutations followed the same pattern (Fig. 4a). The frequency of non-APOBEC mutations decreased as the predicted DNA accessibility and overall gene expression level increased (Fig. 4b). In contrast, the frequency of APOBEC mutations was constant across the genome, demonstrating that APOBEC mutagenesis was likely independent of genomic regions (Fig. 4a-c). The ratio of APOBEC mutations between high and low transcriptional regions also suggests an equal distribution of APOBEC associated mutations across the genome (Fig S14). This ratio was close to 0.5 (0.45- 0.55) in tumors with high APOBEC mutagenesis. The equal distribution of APOBEC mutations across genomic regions supported the hypothesis that these mutations had been generated during replication, when APOBEC enzymes have equal access to ssDNA across the genome [73].

**Figure 4.**
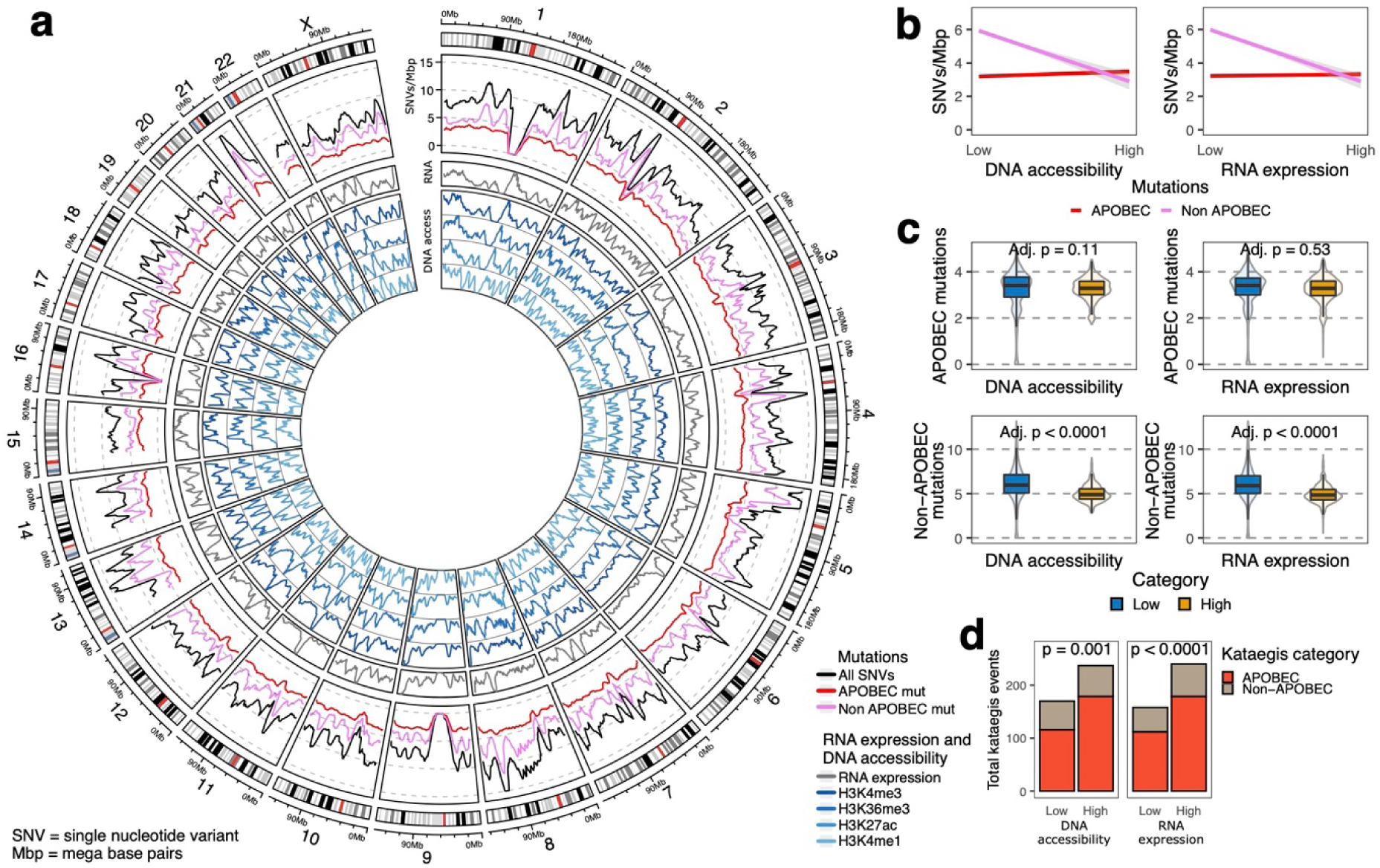
Differences in the load of APOBEC associated mutations between high and low DNA accessibility regions in metastatic urothelial carcinoma genomes. a) WGS data (n = 116) was analyzed to estimate the mean number of single nucleotide variants in windows of one mega base pair across the entire genome. The Circos plot shows from outer to inner circles: the genomics ideogram from chromosome 1 to X where the centrosomes are indicated in red; Mutational load of APOBEC and non-APOBEC associated mutations; Average RNA counts (expression) from 90 tumors with RNA-seq data; DNA accessibility estimation from different ChIPseq experiments in normal urothelial samples derived from the ENCODE [74]. Peaks represent highly accessible DNA. b) Linear regression (with 95% confidence interval) of mutational load per mega base pairs for APOBEC and non-APOBEC associated mutations with DNA accessibility and expression data. c) The number of APOBEC and non-APOBEC associated mutations were compared between high and low RNA expression and DNA accessibility regions. The distributions are shown as boxplots and as violin plots. T-test was applied and p-values were Benjamini-Hochberg corrected for multiple testing. Here, in b) and in c), results from H3K4me1 ChIPseq were used. Using other ChIPseq experiments showed similar results. d) Frequency of kataegis events (n = 116) in high and low DNA accessibility or in high and low RNA expression regions. P-values of binomial test are shown for each comparison.

Localized hypermutation events (kataegis) were present in 70% of the samples (Fig. 2a); which was more frequent when APOBEC mutagenesis was high (Fig. S14). This higher frequency confirmed a link between kataegis and APOBEC activity [75], and we therefore expected to find kataegis events scattered across the genome. However, our data suggested that kataegis was more likely to happen in regions with high DNA accessibility and high transcriptional activity (Fig. 4d). Thus, while general APOBEC mutagenesis seemed to occur primarily during replication, kataegis-like APOBEC events seemed to occur more frequently at transcribed loci.

In summary, APOBEC mutagenesis was an ongoing process in mUC that equally affected the whole genome, and seemed to be triggered by both APOBEC3A and APOBEC3B. Tumors with APOBEC mutagenesis were genomically less stable and displayed more kataegis events.

### Transcriptomic subtypes of mUC

The consensus classifier of primary MIBC stratifies organ-confined UC of the bladder into six transcriptomic subtypes [5]. Unlike primary bladder tumor samples, metastatic biopsies are derived from different organs with some degree of normal non-urothelial cell contamination. Using the consensus classifier would lead to misclassification of samples when no correction for organ-specific transcripts is applied. It would also limit the detection of new phenotypic subtypes if they existed in mUC, which is crucial in this study as the transcriptomic phenotypes of mUC are unknown. Therefore, we performed *de novo* subtyping for mUC samples (see Methods section for details).

High-quality RNA-seq data was available for 90 (97 samples) out of 116 patients (Fig. S15). To reduce the bias introduced by biopsy location, we filtered the organ-specific transcripts prior to hierarchical consensus clustering (Fig. S16). Five transcriptomic subtypes were identified (Fig. 5). We did not observe a neuroendocrine subtype in our cohort, and noticed that the neuroendocrine signature score was equally low in all mUC subtypes (Fig. S17a). Several phenotypic markers were used to establish the phenotype of each subtype (Fig. S17a), and are described below.

**Figure 5.**
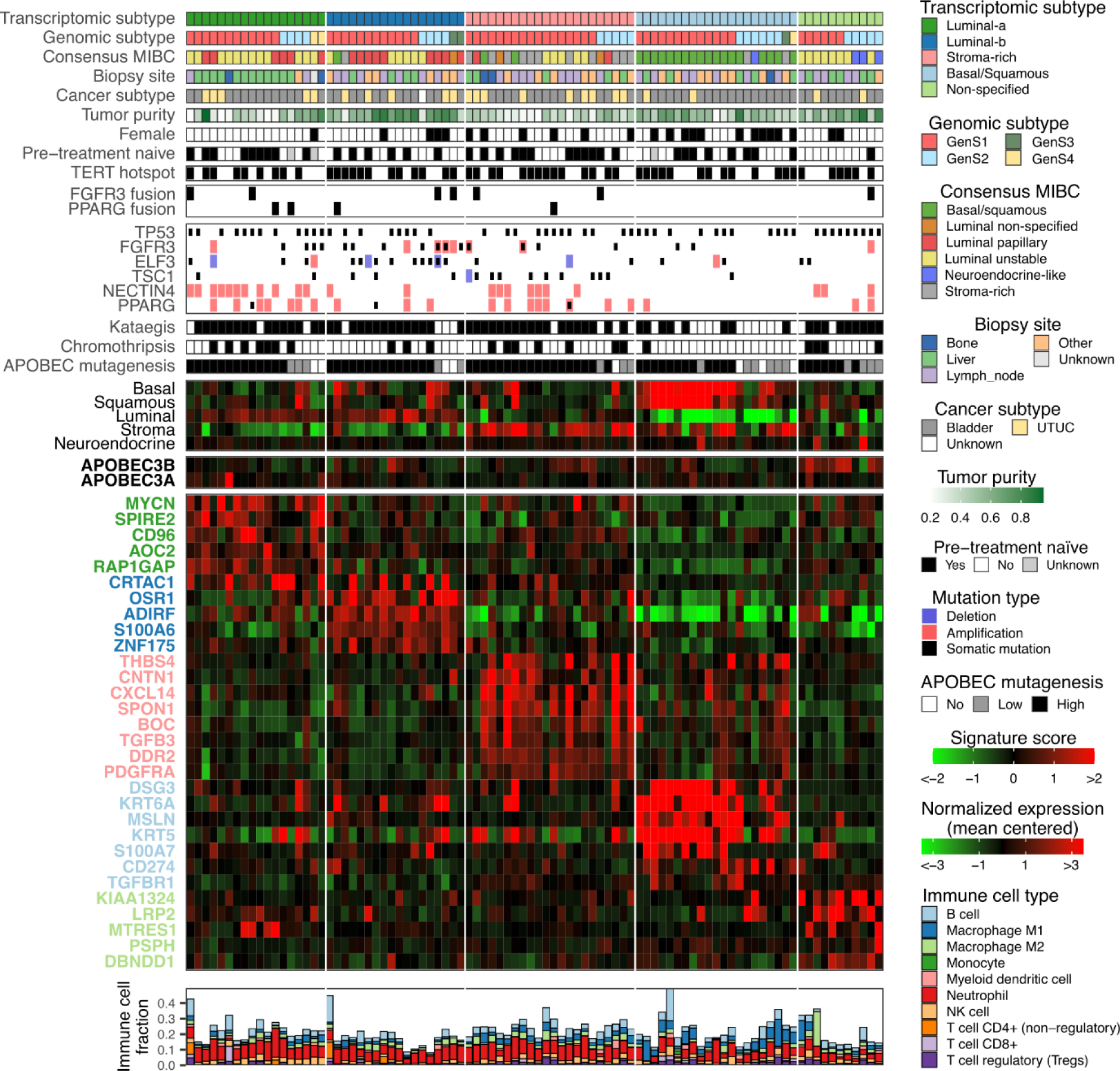
Genomic and transcriptomic characteristics of patients with metastatic urothelial carcinoma stratified by mRNA subtypes. Transcriptomic profiles of 90 metastatic urothelial carcinoma samples were clustered using ConsensusClusterPlus [30]. Five transcriptomic subtypes were identified: luminal-a, luminal-b, stroma, basal/squamous and non-specified phenotype. From top to bottom: Transcriptomic subtypes; Genomic subtypes (GenS1-4); Transcriptional subtypes based on the consensus classifier of primary MIBC [5]; Metastatic site from which a biopsy was obtained; Site of origin of primary tumor (UTUC = upper tract urinary cancer); Estimated tumor cell percentage; Female patients; Pre-treatment naïve patients; Tumors with hotspot mutations in *TERT* promoter; Tumors with gene fusions detected by RNA-seq; Tumors with alterations in selected genes; Tumors with one or more kataegis events; Tumors with one or more chromothripsis events; APOBEC enrichment analysis showing tumors with no-, low- and high-APOBEC mutagenesis; *APOBEC3B* and *APOBEC3A* expression; Signature score (mean expression of genes related to each phenotype) of basal, squamous, luminal, stroma and neuroendocrine markers; Top overexpressed genes in each mRNA subtype; Immune cell fractions estimated with immunedeconv [63], using the quanTIseq method [64].

The luminal subtypes represented 51% of the tumors in the TCGA cohort *versus* 40% in the present cohort (p = 0.061, Fig. S9d). In contrast to the consensus classifier of primary MIBC, we identified two and not three luminal subtypes. The luminal mUC tumors were mostly identified as luminal-papillary and luminal-unstable according to the consensus classifier of primary MIBC. All luminal tumors together exhibited similarities with the consensus-based luminal subtypes [5] with respect to high luminal signature scores, high expression of *PPARG* and *GATA3* (Fig. S16d), and frequent alterations in *FGFR3* and *KDM6A* (Fig. S17). In contrast, the individual luminal mUC subtypes lacked a high TMB; high APOBEC mutation load, as described for the luminal unstable subtype; high stromal signature score, as described for the luminal non-specified subtype; and frequent CDKN2A alterations, as described for the luminal papillary and unstable subtypes. Looking into the characteristics of the two luminal mUC subtypes, we noted that the luminal-a subtype had high expression of *MYCN*, one of the MYC oncogene family members that regulates different species of RNA [76], and high expression of *CD96* (Fig. 5). This subtype had low tumor purity, a high fraction of NK cells, a low clonal fraction (interpreted as high heterogeneity), and relatively high expression of *FGFR3* and *NECTIN4* (Fig. S17). *NECTIN4* was amplified in 61% of these tumors. The luminal-b subtype had high tumor purity, a low number of SVs, a low fraction of NK cells, high expression of *FGFR3* and *S100A6* (Fig. S17), and a higher proportion of *ELF3* (56%) and *FGFR3* (50%) DNA alterations compared to all other subtypes (Fig. S18; Fisher’s exact test p = 0.0023 and p = 0.0053, respectively).

The stroma-rich subtype, had the highest level of stroma signature score. Genes known to be associated with stromal content and cancer-associated fibroblasts (*THBS4*, *CNTN1*, *CXCL14* and *BOC*) [77–80] were differentially expressed (Fig. 5). This subtype was highly concordant with the stratification of the consensus classifier of primary MIBC: 79% of tumors identified as stroma-rich by the consensus classifier of MIBC were in the stroma-rich subtype of mUC. However, the stroma-rich subtype was more prevalent in the present mUC cohort than in the TCGA cohort (24% *vs* 9%, Fisher’s exact test p < 0.001, Fig. S9d). Tumors of the stroma-rich subtype showed high expression of *TGFB3*, a ligand of the TGF-β pathway (Fig. 5), low tumor purity, a high signature score for epithelial to mesenchymal transition (Fig. S17), high expression of various collagens (Table S1.20), and a higher rate of *TSC1* DNA alterations (45% of the tumors, Fig. 5) compared to the rest of the cohort (Fisher’s exact test p < 0.001).

The basal/squamous subtype was also highly concordant to the similarly named cluster of the consensus classifier for MIBC; 86% of tumors identified as basal/squamous by the consensus MIBC were in this group. Yet, the prevalence of this subtype was lower in the present cohort than in the TCGA cohort (23% *vs* 37%, Fisher’s exact test p = 0.013; Fig. S9d). This subtype was characterized by high expression of basal/squamous markers (*DSG3*, *KRT5, KRT6A* and *S100A7*), highest levels of basal and squamous signature scores, and enrichment in females (52%, Fisher’s exact test p = 0.0043). *TGFBR1,* a receptor of the TGF-β pathway; *CD274*, the gene that encodes PD-L1; and *MSLN*, a tumor-associated antigen, were highly expressed in this subtype (Fig. 5). *NECTIN4* amplifications were not found, and *NECTIN4* expression level was low (Fig. S17). In line with a previous study [81], the expression of adipogenesis regulatory factor (*ADIRF*) was low (Fig. 5). The immune cell compartment consisted of a large fraction of M1 macrophages and a low fraction of neutrophils (Fig. S17b).

This subtype was less affected by kataegis and chromothripsis events than the other subtypes (Fig. 5, Fisher’s exact test p = 0.0006 and p = 0.019, respectively).

The majority of samples in the non-specified subtype was identified as luminal unstable according to the consensus classifier of primary MIBC. However, key markers of luminal phenotypes such as the luminal signature score, and *PPARG* and *GATA3* expression were relatively low, and genomic instability (high TMB, high APOBEC mutagenesis) was not observed in this subtype (Fig. 5 and S17). This subtype had overexpression of *KIAA1324;* a diagnostic biomarker in different cancer subtypes [82], and of *LRP2* (Fig. 5),. Furthermore, it had a high score of claudin markers, a low fraction of neutrophils, high numbers of Indels, high numbers of SVs, high levels of *APOBEC3B* expression (Fig. S16), and this subtype was enriched for patients who were previously treated with chemotherapy (Fisher’s exact test p = 0.023, Fig. 5).

In summary, transcriptomic profiling revealed that mUC can be stratified into five transcriptomic subtypes, of which the stroma-rich and basal/squamous subtypes are highly concordant to primary MIBC subtypes. Both luminal subtypes showed some concordance with the luminal MIBC subtypes, however the individual luminal subtypes in mUC and MIBC differed. The phenotype of the non-specified mUC subtype did not match any of the phenotypes of the consensus classifier established for MIBC. A complete overview of driver genes, fusion genes and hotspot mutations per transcriptomic subtype is presented in Fig. S18.

### Altered canonical signaling pathways in different transcriptomic subtypes

Several canonical pathways involved in cell growth, proliferation and survival [34] were altered at the DNA level (Fig. 6a). Of all subtypes, the luminal-a subtype showed most alterations in the Myc (72%) and TGF-β (72%) pathways. In contrast, only 5% of basal/squamous tumors had TGF-β pathway alterations. Perturbations in the TGF-β pathway were mainly driven by alterations in *TGFBR2*, *SMAD4*, *SMAD2* and *TGFBR1*. The most altered genes per pathway are displayed in Fig. S19.

**Figure 6.**
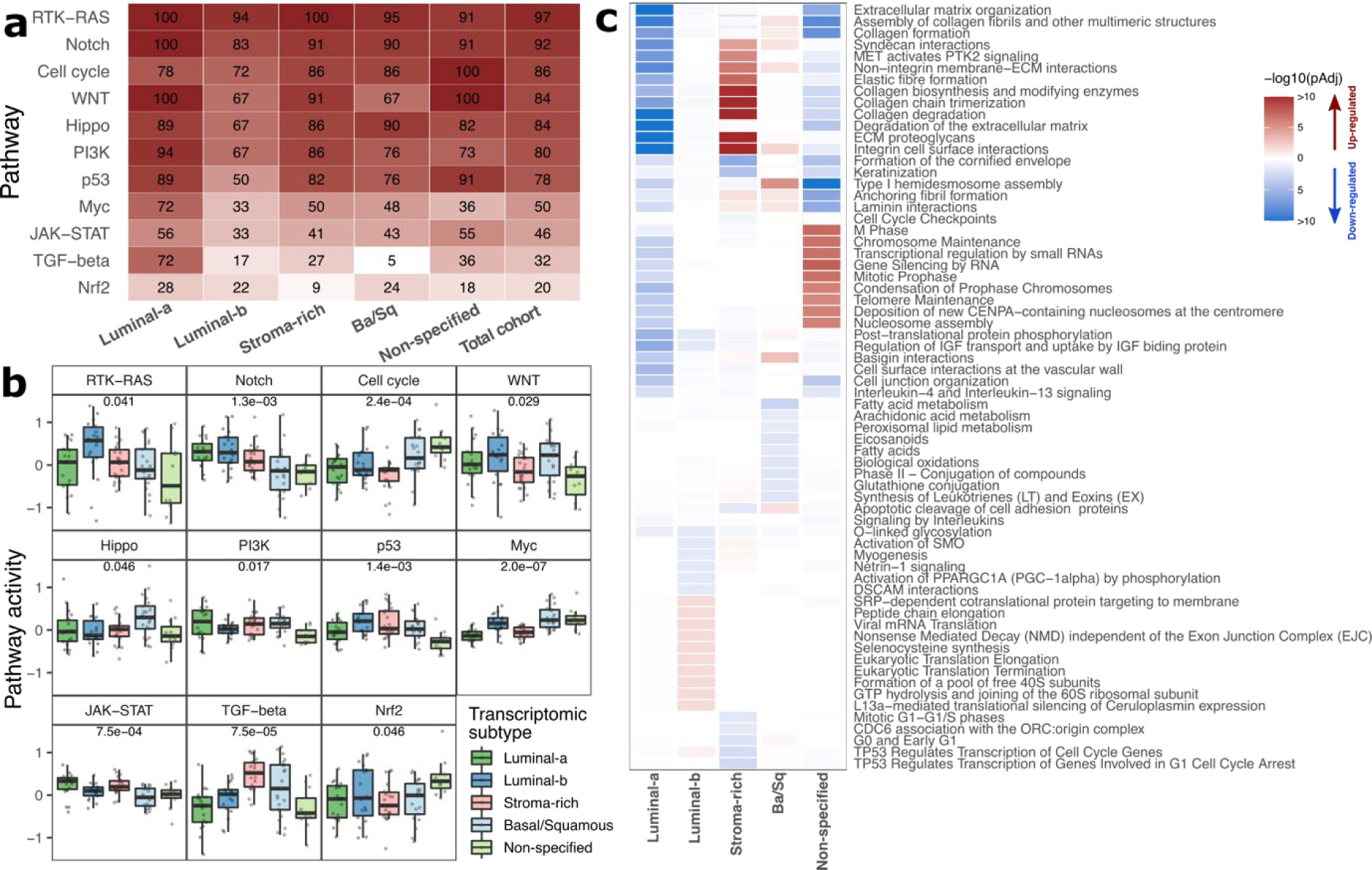
Pathway alterations at genomic and transcriptomic level across mRNA-based subtypes of metastatic urothelial carcinoma. a) The percentage of samples with DNA alterations in 11 canonical pathways is shown for each transcriptomic subtype (90 tumors in total) and for the entire cohort (n = 116). A patient was considered to have an altered pathway when at least one of the pathway-genes was altered either by non-synonymous mutations, structural variants or by deep copy number changes. b) Pathway activity was estimated as the mean expression of downstream genes targeted by each pathway. Only genes that were transcriptionally activated by these pathways were considered. Kruskal-Wallis test p-values were Benjamini-Hochberg corrected. c) Pathways up- (red) or down-regulated (blue) were estimated with reactomePA [62] from RNA-seq data. Only the top ten up- and top ten down-regulated pathways per subtype are shown.

The luminal-b subtype was characterized by fewer alterations in Notch, Cell cycle, Hippo, PI3K, p53, Myc and JAK-STAT pathways. Alterations in the p53 pathway were common in the other subtypes, and most of them were the result of somatic mutations in *TP53* or amplification of *MDM2* (mutually exclusive p = 0.024).

Amplification of *MDM2* has been previously reported in a pan-cancer study [34] as an alternative to *TP53* alterations to inactivate the p53 pathway through direct inhibition of p53 protein [83].

To assess pathway activity across different transcriptomic subtypes, we calculated the mean expression of genes targeted by each pathway as *a proxy* of activity (Fig. 6b). Myc and TGF-β pathway activities were low in the luminal-a subtype, corresponding with high frequencies of pathway alterations at the genomic level (Fig. 6a). The luminal-b subtype showed the highest RTK-RAS and high WNT pathway activity. The stroma-rich subtype had high TGF-β pathway activity. The basal/squamous subtype had high activity of the Hippo, Myc and TGF-β pathways. The non-specified subtype had very low p53 pathway activity and very active cell cycle pathway signaling, two pathways that are usually co-altered [34].

In addition to the 11 oncogenic pathways described above, any pathway up- or down-regulated was analyzed by enriched pathway analysis with ReactomePA (Fig. 6c). Up-regulation of pathways involved in collagen metabolism and extracellular matrix in the stroma-rich subtype corresponded with the stromal phenotype of this subtype [84]. In the non-specified subtype, pathways related to cell cycle and chromosome integrity were up-regulated. Considering as well the high cell cycle pathway activity (Fig. 6b), and high frequency of mutations in the cell cycle pathway (Fig. 6a), this up-regulation may suggest that the non-specified subtype is highly proliferative.

In summary, signaling pathway analysis showed the extent of heterogeneity between the transcriptomic subtypes, reflecting phenotypic characteristics of each group. The most striking difference was observed for the TGF-β pathway, in which genomic alterations greatly affected the luminal-a subtype and pathway activity was heavily reduced.

## Discussion

To the best of our knowledge, we defined, for the first time, molecular subtypes of mUC based on whole genome and transcriptome characteristics of metastatic biopsies of 116 mUC patients. In line with findings reported for primary UC, we identified a central role for APOBEC mutagenesis in mUC. Furthermore, we showed that mUC is a heterogeneous disease with various genomic and transcriptomic subtypes revealing the main mutational processes and phenotypes of this cancer.

The genomic landscape of mUC showed important similarities to that of primary UC. We validated our mUC findings in the TCGA cohort of primary MIBC, showing that aggregating mutational signatures by etiology is a robust approach to identify genomic subtypes. A recent study analyzed archived paraffin-embedded primary or metastatic tumor samples from UC patients who received palliative chemotherapy. By WES analysis, two major genomic subtypes were identified [85]. The GenS2 subtype, enriched with SigG that correlates with COSMIC SBS5, in the present study largely overlapped with the SBS5 subtype reported by Taber *et al.*, 2020. Furthermore, an APOBEC high signature was identified that was similar to GenS1 in our study.

We identified significantly mutated genes similar to those reported for primary UC. Frequent hotspot mutations in the non-coding region of *TERT*, *ADGRG6*, *PLEKHS1*, *LEPROTL1* and *TBC1D12* occurred similarly in NMIBC and MIBC [86, 87]. Evidence suggests that clones with known driver genes emerge early during bladder cancer development and colonize distant areas of the bladder, which may explain the genomic similarity between mUC and primary UC [88].

WGS analysis revealed frequent SVs affecting *AHR* (aryl hydrocarbon receptor) and *CCSER1* (coiled-coil serine rich protein). SVs in *AHR* have not been described in cancer, but other molecular alterations in this gene have been associated with bladder cancer progression [89–91]. *CCSER1* is located in a common fragile site region; thus, it is exposed to chromosomal rearrangements [92]. Altered transcripts created through the deletion of specific exons in *CCSER1* have been associated with oncogenesis [92, 93]; it is unclear, however, if SVs may have similar oncogenic effects in UC.

Previous studies that performed RNA-based subtyping showed that NMIBC is a homogeneous disease primarily of luminal origin (> 90%) and that MIBC is highly heterogeneous with multiple subtypes [5, 94]. Here, we performed *de novo* transcriptomic subtyping of mUC, as the consensus classifier of primary MIBC is not suitable for mUC, and it does not correct for organ-specific transcriptomic contamination. Although this method may also remove transcription related to urothelial adaptation, or to a specific metastatic site, it was mandatory in order to prevent subtyping of tumors to be predominantly based on biopsy site rather than biological differences. We showed that mUC is a heterogeneous disease, with similarities to subtypes described for primary MIBC, specifically regarding the stroma-rich and basal/squamous subtypes. The luminal subtypes overall showed some concordance, although there was not a one to one match with the primary MIBC luminal subtypes. The phenotypic similarity of primary MIBC and mUC suggests that despite ongoing mutagenesis, UC cell behavior does not change significantly during the metastatic process – all subtypes have metastatic potential. However, as matched data from the primary tumor were lacking in this study, we were unable to draw conclusions on specific factors which may drive some clones towards metastasis whilst other remain tissue confined. Furthermore, some patients with primary non-metastatic MIBC, as assessed by cross-sectional imaging, actually have systemic rather than localized disease. In a previous study, lymph node metastases were present in the cystectomy resection specimen of 25% of clinically node-negative patients. In addition, patients with locally advanced bladder tumors (pathological stage T3) have a poor 5-year overall survival rate of only 35% due to rapid onset of metastatic disease, despite radical surgery [95, 96].

In the present cohort, we did not identify a NE-like subtype at the transcriptional level. The prevalence of this subtype in UC is, however low; in the TCGA cohort only 2% of tumors were of the NE-like subtype. Central pathology revision of the metastatic biopsies identified only three NE-like tumors in our cohort (Table S3-4). The non-specified subtype we identified did not express any of the markers used to identify the consensus subtypes of primary MIBC (luminal, basal, squamous, stroma or NE), suggesting rewiring of its transcriptomic profile for adaptation. Studies in various cancers have shown that therapeutic pressure may trigger a phenotype-switching event [97], which could have happened in the non-specified phenotype as it was enriched for patients who had received systemic therapy prior to biopsy. Studies with larger numbers of paired biopsies, obtained before and after treatment, and obtained from the primary and metastatic site would be needed to explore this phenomenon in mUC.

APOBEC mutagenesis was widespread in mUC; the reconstruction of evolutionary paths from sequential biopsies of eight patients indicated that it was an ongoing process, which was confirmed in the entire cohort by RNA-editing of *DDOST* attributed to APOBEC3A. This suggests that mUC is in continuous adaptation by generating novel mutations. A previous study indeed reported accumulating mutations in six patients whose primary tumor and metachronous metastases were analyzed by WES [85]. In our study, the accumulation of new mutations in the sequential biopsy specimen of one of eight patients led to the identification of new therapeutic targets (Fig. S20).

In a previous study [7], the genomic landscape of 85 (72 re-analyzed here) mUC patients was compared with that of other metastatic tumor types. This pan-cancer study concluded that mUC was characterized by high tumor mutational load, with no difference between mUC and primary UC, high CNAs, the highest number of driver genes among all cancer types analyzed, and actionable targets in 75% of the patients. In our study, we identified a potential targetable alteration in the genome of 98% of the patients (Fig. 7a). In line with Priestley *et al.*, 2019, we found that 41% of patients could benefit from on-label therapies, and 63% from therapies approved by the US Food and Drug Administration for other tumor types. Additionally, we identified targets for therapies under investigation in clinical trials including basket trials in 109 of 116 patients. We identified four patients with MSI-high tumors that are potentially sensitive to immune checkpoint inhibitors [98]. HR deficiency, observed in three patients, is a potential target for treatment with poly-ADP ribose polymerase inhibitors and/or double-stranded DNA break-inducing chemotherapy. At the RNA level, targetable *FGFR3* and *NTRK2* gene fusions were identified in eight patients.

**Figure 7.**
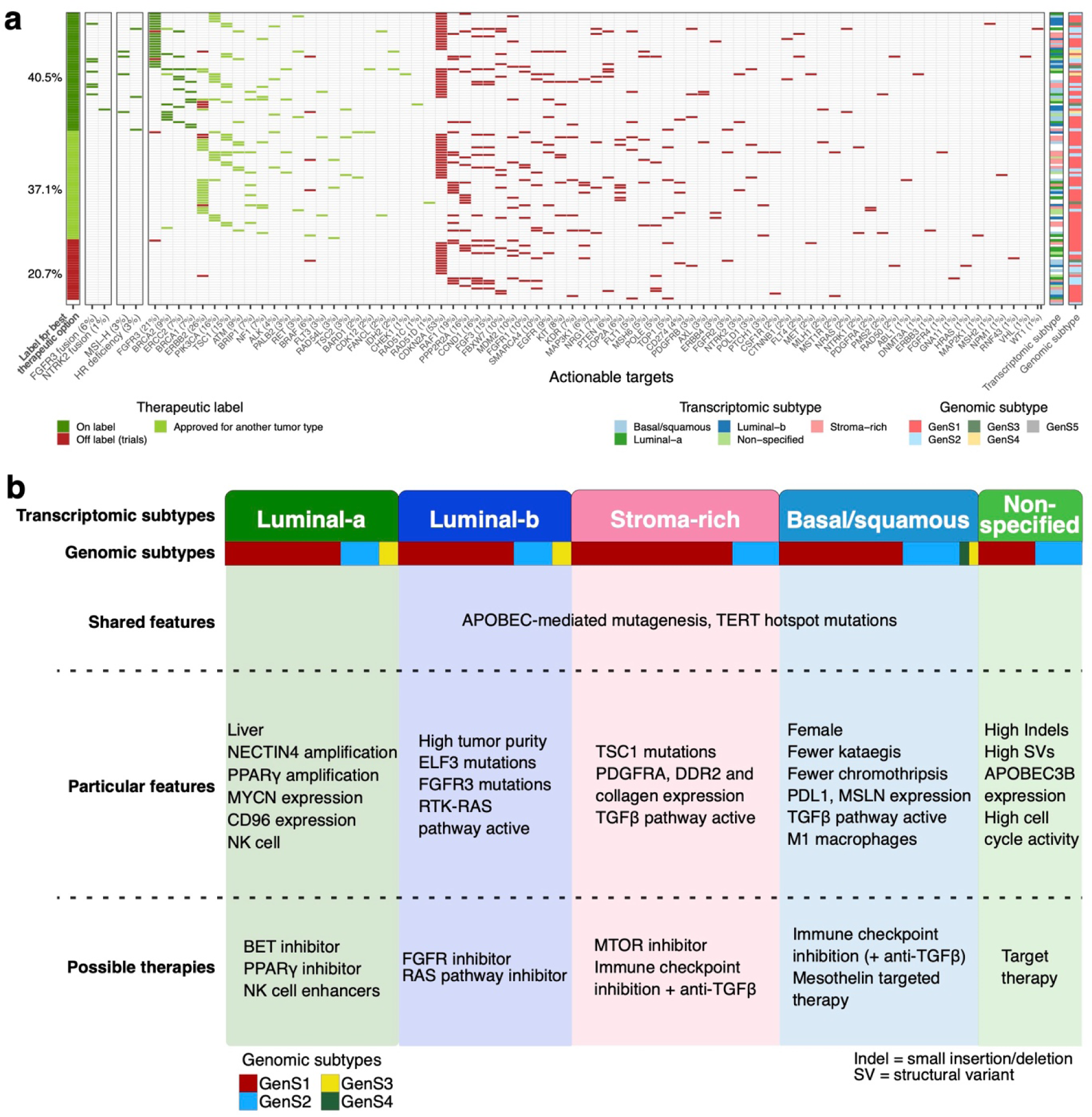
Overview of actionable targets and possible treatments per transcriptomic subtype for metastatic urothelial carcinoma. a) Per patient overview of therapeutic targets based on gene fusions at RNA level (first column), tumors with microsatellite instability high (MSI-H) tumors, or homologous recombination (HR) deficiency (second column), and clinically-actionable genomic alterations for on- and off-label therapies for urothelial carcinoma (third column). On the left side, the therapeutic label for the best treatment option per patient is shown. Bars on the right depict the genomic and the transcriptomic subtype per patient. b) Summary of molecular characteristics found in the present study, and potential therapeutic implications for the treatment of metastatic urothelial carcinoma per transcriptomic subtype. From top to bottom: transcriptomic mUC subtypes, genomic mUC subtypes, shared genomic features among transcriptomic subtypes; unique characteristics per transcriptomic subtype; suggested therapeutic strategies per transcriptomic subtype.

In a previous study, the antibody-drug conjugate enfortumab vedotin targeting NECTIN4 induced objective clinical responses in 44% of mUC patients who experienced disease progression following chemotherapy and anti-PD1/L1 therapy [99]. Currently, preselection for this treatment is not required. However, we found significant variation in the expression of *NECTIN4*, suggesting that patients with tumors of the basal/squamous subtype may be less likely to experience clinical benefit, as no *NECTIN4* amplifications were detected, and *NECTIN4* expression levels were low. Thus, subtype-specific treatment with enfortumab vedotin might result in better risk-benefit ratios. The 23 patients with *HER2* aberrations may be sensitive to HER2 targeting agents; especially some of the newer antibody-drug conjugates with DNA damaging payloads could represent an effective treatment [100, 101].

Based on the identified transcriptomic subtypes we suggested potential therapeutic targets per subtype (Fig. 7b). The luminal-a subtype was characterized by *MYCN* and *PPARGC1B* overexpression. In pre-clinical studies, treatment with a BET- or PPARγ-inhibitor downregulated the expression levels of both genes, and had an antiproliferative effect on tumor cells [102, 103]. The immune cell compartment of tumors of the luminal-a subtype was found rich in NK cells, which could be explained by the large fraction of liver biopsies, which are known to be enriched for NK cells [104]. Thus, other potential treatment strategies comprise of cytokine-mediated stimulation of NK cells and TLR agonists [105].

The luminal-b subtype was enriched for *FGFR3* mutations and had high expression of *FGFR3*, suggesting that this subtype may be susceptible to FGFR inhibitors. This subtype may also be sensitive to RAS pathway inhibitors as the RTK-RAS pathway activity was high [106].

The stroma-rich subtype was characterized by *TSC1* alterations that confer sensitivity to MTOR inhibitors, which have been approved for treatment of several tumor types [37, 38]. Compared with the other subtypes, the stroma-rich subtype displayed the highest TGF-β pathway activity and overexpression of different collagens. Previous studies have shown that TGF-β can stimulate cancer-associated fibroblasts to produce collagens [107, 108]. Other studies found that TGF-β expression was associated with resistance to immune checkpoint inhibition in bladder cancer [52, 109]. Results from pre-clinical studies suggest that addition of a TGF-β inhibitor may improve anti-PD1 efficacy [53].

The basal/squamous subtype has been found associated with high immune cell infiltration (significantly more M1 macrophages) and overexpression of PD-L1, which suggests that patients with tumors of this subtype are likely to benefit from immunotherapy [3]. Since TGF-β pathway activity was also high in this subtype, combination therapy with a TGF-β inhibitor could be of added value. Furthermore, this subtype was characterized by overexpression of mesothelin, a known tumor antigen that is being investigated as a target for antibody-based, vaccine and CAR-T cell therapies in several tumor types [110].

A limitation of this study is the lack of matched primary tumor samples. Despite this, we showed striking genomic and transcriptomic similarities between mUC and what has been reported for primary MIBC. Our results, however, require validation in other independent mUC cohorts. Also, as our studied cohort was heterogeneous regarding pre-treatment history and type of treatment initiated after biopsy collection, we were unable to correlate the characteristics of the molecular subtypes to clinical endpoints such as overall survival. Additional studies in which biopsies are collected from uniformly treated mUC patients would be crucial to be able to properly correlate large scale genomic and transcriptomic data with clinical outcomes.

### Conclusions

By performing WGS and RNA-seq analysis of metastatic sites of 116 mUC patients who participated in a clinical trial, this study contributed to the knowledge on the molecular landscape of mUC, which has important similarities to the molecular landscape of primary UC. The findings reported here serve as a reference for subtype-oriented and patient-specific research on the etiology of mUC and for novel drug development – with the ultimate aim to improve the management of mUC patients.

### List of abbreviations

APOBEC: Apolipoprotein B mRNA Editing Catalytic Polypeptide-like
CCF: Cancer cell fraction
CNA: Copy number alterations
CPCT: The Center for Personalized Cancer Treatment
FC: Fold change
H&E: Hematoxylin and eosin
HMF: Hartwig Medical Foundation
HR: Homologues recombination
Indel: insertion/deletion
Mbp: Megabase pair
MIBC: Muscle invasive bladder cancer
MMR: Mismatch repair
MNV: Multiple nucleotide variants
MSI: Microsatellite instability
mUC: Metastatic urothelial carcinoma
NE-like: Neuroendrocrine-like
NMF: Non-negative matrix factorization
NMIBC: Non-muscle invasive bladder cancer
RNA-seq: RNA sequencing
SNV: Single nucleotide variant
SV: Structural variant
TAF: Tumor allele frequency
TCGA: The Cancer Genome Atlas
TMB: Tumor mutational burden
UC: Urothelial carcinoma
UTUC: Upper tract urothelial carcinoma
VEP: Variant effect predictor
WES: whole-exome sequencing
WGS: Whole-genome sequencing

## Declarations

### Ethics approval and consent to participate

Patients studied here were included in the CPCT-02 Biopsy Protocol (ClinicalTrial.gov no. NCT01855477) and the DRUP Trial (ClinicalTrial.gov no. NCT02925234). The study protocols were approved by the medical ethics review board of the University Medical Center Utrecht and the Netherlands Cancer Institute. Written informed consent was obtained from all participants prior to inclusion in the trials; the studies comply with all relevant ethical regulations.

### Consent for publication

All patients studied here provided consent to report individual (anonymised) patient data.

### Availability of data and materials

WGS data, RNA-seq data and corresponding clinical data have been requested from the HMF and were provided under data request number DR-031. All data are freely available for academic use from the HMF through standardized procedures. Request forms can be found at https://www.hartwigmedicalfoundation.nl [7].

ChIPseq data experiments are freely available through The ENCODE Project Consortium [111] and the Roadmap Epigenomics Consortium [112] on the ENCODE portal (https://www.encodeproject.org) [74]. Files were downloaded with the following identifiers: ENCSR065IQH, ENCSR054BKO, ENCSR632OWD and ENCSR449TNC.

TCGA data for the bladder cancer cohort was downloaded through the portal: https://www.cancer.gov/tcga.

## Author’s contributions

Conceptualization: JAN, MR, HJGvdW, MPJL, JLB; Methodology: JAN, MR, HJGvdW, MPJL, and JLB; Software: JAN, HJGvdW, and JvR; Validation: MPJL, JLB, JvR ; Formal Analysis: JAN and HJGvdW; Investigation: MR, MSvdH, JV, NM, SvW, SO, HMW, ECZ, RdW, AAMvdV; Resources: HJGvdW, MPJL, JLB, MSvdH, JV, EC, NM, SvW, SO, HMW, ECZ, RdW, AAMvdW, MPJL and JLB; Data Curation: JAN, MR, HJGvdW, JvR, and EC ; Writing – Original Draft: JAN, MR, HJGvdW, MPJL and JLB; Writing – Review & Editing: JAN, MR, JvR, MSvdH, JV, EC, NM, SvW, SO, HMW, ECZ, RdW, AAMvdW, HJGvdW, MPJL and JLB; Visualization: JAN and MR; Supervision: JLB, HJGvdW and MPJL; Project Administration: JAN, MR, HJGvdW, MPJL and JLB; Funding Acquisition: JLB, HJGvdW, and MPJL. All authors read and approved the final manuscript.

## Acknowledgements

The Hartwig Medical Foundation and the Center of Personalized Cancer Treatment are acknowledged for making the clinical and genomic data available to the study. We thank all local principal investigators and the nurses of all contributing centers for their help with patient recruitment. We are particularly grateful to all participating patients and their families.

## References

1. Knowles MA, Hurst CD. Molecular biology of bladder cancer: New insights into pathogenesis and clinical diversity. Nat Rev Cancer. 2015;15(1):25–41.

2. Lindskrog SV, Prip F, Lamy P, Taber A, Groeneveld CS, Birkenkamp-Demtröder K, et al. An integrated multi-omics analysis identifies prognostic molecular subtypes of non-muscle-invasive bladder cancer. Nat Commun. 2021;12(1):2301.

3. Robertson AG, Kim J, Al-Ahmadie H, Bellmunt J, Guo G, Cherniack AD, et al. Comprehensive Molecular Characterization of Muscle-Invasive Bladder Cancer. Cell. 2017;171(3):540–556.e25.

4. Giannopoulou A, Velentzas A, Konstantakou E, Avgeris M, Katarachia S, Papandreou N, et al. Revisiting Histone Deacetylases in Human Tumorigenesis: The Paradigm of Urothelial Bladder Cancer. Int J Mol Sci. 2019;20(6):1291.

5. Kamoun A, Reyniès A de, Allory Y, Sjödahl G, Gordon Robertson A, Seiler R, et al. A Consensus Molecular Classification of Muscle-invasive Bladder Cancer. Eur Urol. 2019;77(4):420–33.

6. Faltas BM, Prandi D, Tagawa ST, Molina AM, Nanus DM, Sternberg C, et al. Clonal evolution of chemotherapy-resistant urothelial carcinoma. Nat Genet. 2016;48(12):1490–9.

7. Priestley P, Baber J, Lolkema MP, Steeghs N, de Bruijn E, Shale C, et al. Pan-cancer whole-genome analyses of metastatic solid tumours. Nature. 2019;575(7781):210–6.

8. Casparie M, Tiebosch ATMG, Burger G, Blauwgeers H, Pol A van de, Krieken JHJM van, et al. Pathology Databanking and Biobanking in The Netherlands, a Central Role for PALGA, the Nationwide Histopathology and Cytopathology Data Network and Archive. Cell Oncol. 2007;29(1):19.

9. Li H, Durbin R. Fast and accurate short read alignment with Burrows-Wheeler transform. Bioinformatics. 2009;25(14):1754–60.

10. McKenna A, Hanna M, Banks E, Sivachenko A, Cibulskis K, Kernytsky A, et al. The genome analysis toolkit: A MapReduce framework for analyzing next-generation DNA sequencing data. Genome Res. 2010;20(9):1297–303.

11. Van der Auwera GA, Carneiro MO, Hartl C, Poplin R, del Angel G, Levy-Moonshine A, et al. From fastQ data to high-confidence variant calls: The genome analysis toolkit best practices pipeline. Curr Protoc Bioinforma. 2013;43(SUPL.43):11.10.1–11.10.33.

12. Saunders CT, Wong WSW, Swamy S, Becq J, Murray LJ, Cheetham RK. Strelka: Accurate somatic small- variant calling from sequenced tumor-normal sample pairs. Bioinformatics. 2012;28(14):1811–7.

13. McLaren W, Gil L, Hunt SE, Riat HS, Ritchie GRS, Thormann A, et al. The Ensembl Variant Effect Predictor. Genome Biol. 2016;17(1):122.

14. Liu X, Wu C, Li C, Boerwinkle E. dbNSFP v3.0: A One-Stop Database of Functional Predictions and Annotations for Human Nonsynonymous and Splice-Site SNVs. Hum Mutat. 2016;37(3):235–41.

15. Lek M, Karczewski KJ, Minikel E V., Samocha KE, Banks E, Fennell T, et al. Analysis of protein-coding genetic variation in 60,706 humans. Nature. 2016;536(7616):285–91.

16. Cameron D, Baber J, Shale C, Papenfuss A, Valle-Inclan JE, Besselink N, et al. GRIDSS, PURPLE, LINX: Unscrambling the tumor genome via integrated analysis of structural variation and copy number. bioRxiv preprint. 2019; https://doi.org/10.1101/781013.

17. Martincorena I, Raine KM, Gerstung M, Dawson KJ, Haase K, Van Loo P, et al. Universal Patterns of Selection in Cancer and Somatic Tissues. Cell. 2017;171(5):1029–1041.e21.

18. van Dessel LF, van Riet J, Smits M, Zhu Y, Hamberg P, van der Heijden MS, et al. The genomic landscape of metastatic castration-resistant prostate cancers reveals multiple distinct genotypes with potential clinical impact. Nat Commun. 2019;10(1):1–13.

19. Mermel CH, Schumacher SE, Hill B, Meyerson ML, Beroukhim R, Getz G. GISTIC2.0 facilitates sensitive and confident localization of the targets of focal somatic copy-number alteration in human cancers. Genome Biol. 2011;12(4):R41.

20. Roberts SA, Lawrence MS, Klimczak LJ, Grimm SA, Fargo D, Stojanov P, et al. An APOBEC cytidine deaminase mutagenesis pattern is widespread in human cancers. Nat Genet. 2013;45(9):970–6.

21. Chan K, Roberts SA, Klimczak LJ, Sterling JF, Saini N, Malc EP, et al. An APOBEC3A hypermutation signature is distinguishable from the signature of background mutagenesis by APOBEC3B in human cancers. Nat Genet. 2015;47(9):1067–72.

22. Stephens PJ, Tarpey PS, Davies H, Van Loo P, Greenman C, Wedge DC, et al. The landscape of cancer genes and mutational processes in breast cancer. Nature. 2012;486(7403):400–4.

23. Gundem G, Van Loo P, Kremeyer B, Alexandrov LB, Tubio JMC, Papaemmanuil E, et al. The evolutionary history of lethal metastatic prostate cancer. Nature. 2015;520(7547):353–7.

24. Tate JG, Bamford S, Jubb HC, Sondka Z, Beare DM, Bindal N, et al. COSMIC: The Catalogue Of Somatic Mutations In Cancer. Nucleic Acids Res. 2019;47(D1):D941–7.

25. Blokzijl F, Janssen R, van Boxtel R, Cuppen E. MutationalPatterns: Comprehensive genome-wide analysis of mutational processes. Genome Med. 2018;10(1):33.

26. Alexandrov LB, Kim J, Haradhvala NJ, Huang MN, Tian Ng AW, Wu Y, et al. The repertoire of mutational signatures in human cancer. Nature. 2020;578(7793):94–101.

27. Petljak M, Alexandrov LB, Brammeld JS, Price S, Wedge DC, Grossmann S, et al. Characterizing Mutational Signatures in Human Cancer Cell Lines Reveals Episodic APOBEC Mutagenesis. Cell. 2019;176(6):1282–1294.e20.

28. Angus L, Smid M, Wilting SM, van Riet J, Van Hoeck A, Nguyen L, et al. The genomic landscape of metastatic breast cancer highlights changes in mutation and signature frequencies. Nat Genet. 2019;51(10):1450–8.

29. Christensen S, Van der Roest B, Besselink N, Janssen R, Boymans S, Martens JWM, et al. 5-Fluorouracil treatment induces characteristic T>G mutations in human cancer. Nat Commun. 2019;10(1):1–11.

30. Wilkerson MD, Hayes DN. ConsensusClusterPlus: A class discovery tool with confidence assessments and item tracking. Bioinformatics. 2010;26(12):1572–3.

31. Gaujoux R, Seoighe C. A flexible R package for nonnegative matrix factorization. BMC Bioinformatics. 2010;11(1):367.

32. Cortés-Ciriano I, Lee JJK, Xi R, Jain D, Jung YL, Yang L, et al. Comprehensive analysis of chromothripsis in 2,658 human cancers using whole-genome sequencing. Nat Genet. 2020;52(3):331–41.

33. Nguyen L, Martens J, Hoeck A van, Cuppen E. Pan-cancer landscape of homologous recombination deficiency. Nat Commun. 2020;11:5584.

34. Sanchez-Vega F, Mina M, Armenia J, Chatila WK, Luna A, La KC, et al. Oncogenic Signaling Pathways in The Cancer Genome Atlas. Cell. 2018;173(2):321–337.e10.

35. Leonard WJ. Role of JAK kinases and stats in cytokine signal transduction. Int J Hematol. 2001;73(3):271–7.

36. Griffith M, Spies NC, Krysiak K, McMichael JF, Coffman AC, Danos AM, et al. CIViC is a community knowledgebase for expert crowdsourcing the clinical interpretation of variants in cancer. Nat Genet. 2017;49(2):170–4.

37. Chakravarty D, Gao J, Phillips S, Kundra R, Zhang H, Wang J, et al. OncoKB: A Precision Oncology Knowledge Base. JCO Precis Oncol. 2017;1:1–16.

38. Tamborero D, Rubio-Perez C, Deu-Pons J, Schroeder MP, Vivancos A, Rovira A, et al. Cancer Genome Interpreter annotates the biological and clinical relevance of tumor alterations. Genome Med. 2018;10(1):25.

39. Nakato R, Shirahige K. Recent advances in ChIP-seq analysis: from quality management to whole- genome annotation. Brief Bioinform. 2017;18(2):279–90.

40. Bolger AM, Lohse M, Usadel B. Trimmomatic: a flexible trimmer for Illumina sequence data. Bioinformatics. 2014;30(15):2114–20.

41. Dobin A, Davis CA, Schlesinger F, Drenkow J, Zaleski C, Jha S, et al. STAR: Ultrafast universal RNA-seq aligner. Bioinformatics. 2013;29:15–21.

42. Harrow J, Frankish A, Gonzalez JM, Tapanari E, Diekhans M, Kokocinski F, et al. GENCODE: The reference human genome annotation for the ENCODE project. Genome Res. 2012;22(9):1760–74.

43. Tarasov A, Vilella AJ, Cuppen E, Nijman IJ, Prins P. Sambamba: Fast processing of NGS alignment formats. Bioinformatics. 2015;31(12):2032–4.

44. Wang L, Nie J, Sicotte H, Li Y, Eckel-Passow JE, Dasari S, et al. Measure transcript integrity using RNA- seq data. BMC Bioinformatics. 2016;17:58.

45. Wang L, Wang S, Li W. RSeQC: quality control of RNA-seq experiments. Bioinformatics. 2012;28(16):2184–5.

46. Liao Y, Smyth GK, Shi W. FeatureCounts: An efficient general purpose program for assigning sequence reads to genomic features. Bioinformatics. 2014;30(7):923–30.

47. Li B, Dewey CN. RSEM: Accurate transcript quantification from RNA-seq data with or without a reference genome. BMC Bioinformatics. 2011;12:323.

48. Ramírez F, Ryan DP, Grüning B, Bhardwaj V, Kilpert F, Richter AS, et al. deepTools2: a next generation web server for deep-sequencing data analysis. Nucleic Acids Res. 2016;44(W1):W160–5.

49. Williams SB, Black PC, Dyrskjøt L, Seiler R, Schmitz-Dräger B, Nawroth R, et al. Re: Aurélie Kamoun, Aurélien de Reyniès, Yves Allory, et al. A Consensus Molecular Classification of Muscle-invasive Bladder Cancer. Eur Urol 2020;77:420–33: A Statement from the International Bladder Cancer Network. Vol. 77, European Urology. Elsevier B.V.; 2020. p. e105–6.

50. Love MI, Huber W, Anders S. Moderated estimation of fold change and dispersion for RNA-seq data with DESeq2. Genome Biol. 2014;15(12):550.

51. Paulson JN, Chen CY, Lopes-Ramos CM, Kuijjer ML, Platig J, Sonawane AR, et al. Tissue-aware RNA-Seq processing and normalization for heterogeneous and sparse data. BMC Bioinformatics. 2017;18(1):1– 10.

52. Powles T, Kockx M, Rodriguez-Vida A, Duran I, Crabb SJ, Van Der Heijden MS, et al. Clinical efficacy and biomarker analysis of neoadjuvant atezolizumab in operable urothelial carcinoma in the ABACUS trial. Nat Med. 2019;25(11):1706–14.

53. Mariathasan S, Turley SJ, Nickles D, Castiglioni A, Yuen K, Wang Y, et al. TGFβ attenuates tumour response to PD-L1 blockade by contributing to exclusion of T cells. Nature. 2018;554(7693):544–8.

54. Borggrefe T, Oswald F. The Notch signaling pathway: Transcriptional regulation at Notch target genes. Cell Mol Life Sci. 2009;66(10):1631–46.

55. van Ooijen H, Hornsveld M, Dam-de Veen C, Velter R, Dou M, Verhaegh W, et al. Assessment of Functional Phosphatidylinositol 3-Kinase Pathway Activity in Cancer Tissue Using Forkhead Box-O Target Gene Expression in a Knowledge-Based Computational Model. Am J Pathol. 2018;188(9):1956– 72.

56. Varelas X. The hippo pathway effectors TAZ and YAP in development, homeostasis and disease. Dev. 2014;141(8):1614–26.

57. Fischer M. Census and evaluation of p53 target genes. Oncogene. 2017;36(28):3943–56.

58. Kitamura H, Motohashi H. NRF2 addiction in cancer cells. Cancer Sci. 2018;109(4):900–11.

59. Hartl M. The quest for targets executing MYC-dependent cell transformation. Vol. 6, Frontiers in Oncology. Frontiers Research Foundation; 2016. p. 132.

60. Wagle M-C, Kirouac D, Klijn C, Liu B, Mahajan S, Junttila M, et al. A transcriptional MAPK Pathway Activity Score (MPAS) is a clinically relevant biomarker in multiple cancer types. npj Precis Oncol. 2018;2(1):1–12.

61. Murray PJ. The JAK-STAT Signaling Pathway: Input and Output Integration. J Immunol. 2007;178(5):2623–9.

62. Yu G, He QY. ReactomePA: An R/Bioconductor package for reactome pathway analysis and visualization. Mol Biosyst. 2016;12(2):477–9.

63. Sturm G, Finotello F, Petitprez F, Zhang JD, Baumbach J, Fridman WH, et al. Comprehensive evaluation of transcriptome-based cell-type quantification methods for immuno-oncology. Bioinformatics. 2019;35(14):i436–45.

64. Finotello F, Mayer C, Plattner C, Laschober G, Rieder D, Hackl H, et al. Molecular and pharmacological modulators of the tumor immune contexture revealed by deconvolution of RNA-seq data. Genome Med. 2019;11(1):34.

65. Jang YE, Jang I, Kim S, Cho S, Kim D, Kim K, et al. ChimerDB 4.0: an updated and expanded database of fusion genes. Nucleic Acids Res. 2019;48(D1):D817–24.

66. Dang HX, White BS, Foltz SM, Miller CA, Luo J, Fields RC, et al. ClonEvol: clonal ordering and visualization in cancer sequencing. Ann Oncol. 2017;28(12):3076–82.

67. R Core Team. R Core Team (2017). R: A language and environment for statistical computing. R Found Stat Comput Vienna, Austria URL http://www.R-project.org/. 2017;R Foundation for Statistical Computing.

68. Allory Y, Beukers W, Sagrera A, Flández M, Marqués M, Márquez M, et al. Telomerase reverse transcriptase promoter mutations in bladder cancer: High frequency across stages, detection in urine, and lack of association with outcome. Eur Urol. 2014;65(2):360–6.

69. Buisson R, Langenbucher A, Bowen D, Kwan EE, Benes CH, Zou L, et al. Passenger hotspot mutations in cancer driven by APOBEC3A and mesoscale genomic features. Science. 2019;364(6447):eaaw2872.

70. Glaser AP, Fantini D, Wang Y, Yu Y, Rimar KJ, Podojil JR, et al. APOBEC-mediated mutagenesis in urothelial carcinoma is associated with improved survival, mutations in DNA damage response genes, and immune response. Oncotarget. 2018;9(4):4537–48.

71. Cortez LM, Brown AL, Dennis MA, Collins CD, Brown AJ, Mitchell D, et al. APOBEC3A is a prominent cytidine deaminase in breast cancer. PLoS Genet. 2019;15(12):e1008545.

72. Kazanov MD, Roberts SA, Polak P, Stamatoyannopoulos J, Klimczak LJ, Gordenin DA, et al. APOBEC- Induced Cancer Mutations Are Uniquely Enriched in Early-Replicating, Gene-Dense, and Active Chromatin Regions. Cell Rep. 2015;13(6):1103–9.

73. Hoopes JII, Cortez LMM, Mertz TMM, Malc EPP, Mieczkowski PAA, Roberts SAA. APOBEC3A and APOBEC3B Preferentially Deaminate the Lagging Strand Template during DNA Replication. Cell Rep. 2016;14(6):1273–82.

74. Davis CA, Hitz BC, Sloan CA, Chan ET, Davidson JM, Gabdank I, et al. The Encyclopedia of DNA elements (ENCODE): data portal update. Nucleic Acids Res. 2018;46(D1):D794–801.

75. Nik-Zainal S, Alexandrov LB, Wedge DC, Van Loo P, Greenman CD, Raine K, et al. Mutational processes molding the genomes of 21 breast cancers. Cell. 2012;149(5):979–93.

76. Baluapuri A, Wolf E, Eilers M. Target gene-independent functions of MYC oncoproteins. Nat Rev Mol Cell Biol. 2020;21(5):255–67.

77. Kuroda K, Yashiro M, Sera T, Yamamoto Y, Kushitani Y, Sugimoto A, et al. The clinicopathological significance of Thrombospondin-4 expression in the tumor microenvironment of gastric cancer. Katoh M, editor. PLoS One. 2019;14(11):e0224727.

78. Gu Y, Li T, Kapoor A, Major P, Tang D. Contactin 1: An important and emerging oncogenic protein promoting cancer progression and metastasis. Genes (Basel). 2020;11(8):1–22.

79. Zhang Q, Zhou N, Wang W, Zhou S. A novel autocrine CXCL14/ACKR2 Axis: The achilles’ heel of cancer metastasis? Clin Cancer Res. 2019;25(12):3476–8.

80. Mathew E, Zhang Y, Holtz AM, Kane KT, Song JY, Allen BL, et al. Dosage-dependent regulation of pancreatic cancer growth and angiogenesis by Hedgehog signaling. Cell Rep. 2014;9(2):484–94.

81. Eriksson P, Aine M, Veerla S, Liedberg F, Sjödahl G, Höglund M. Molecular subtypes of urothelial carcinoma are defined by specific gene regulatory systems. BMC Med Genomics. 2015;8(1):25.

82. Dieters-Castator DZ, Rambau PF, Kelemen LE, Siegers GM, Lajoie GA, Postovit LM, et al. Proteomics- derived biomarker panel improves diagnostic precision to classify endometrioid and high-grade serous ovarian carcinoma. Clin Cancer Res. 2019;25(14):4309–19.

83. Haupt Y, Maya R, Kazaz A, Oren M. Mdm2 promotes the rapid degradation of p53. Nature. 1997;387(6630):296–9.

84. Nissen NI, Karsdal M, Willumsen N. Collagens and Cancer associated fibroblasts in the reactive stroma and its relation to Cancer biology. J Exp Clin Cancer Res. 2019;38(1):115.

85. Taber A, Christensen E, Lamy P, Nordentoft I, Prip F, Lindskrog SV, et al. Molecular correlates of cisplatin-based chemotherapy response in muscle invasive bladder cancer by integrated multi-omics analysis. Nat Commun. 2020;11(1):1–15.

86. Wu S, Ou T, Xing N, Lu J, Wan S, Wang C, et al. Whole-genome sequencing identifies ADGRG6 enhancer mutations and FRS2 duplications as angiogenesis-related drivers in bladder cancer. Nat Commun. 2019;10(1):1–12.

87. Bell RJA, Rube HT, Xavier-Magalhães A, Costa BM, Mancini A, Song JS, et al. Understanding TERT promoter mutations: A common path to immortality. Mol Cancer Res. 2016;14(4):315–23.

88. Lawson ARJ, Abascal F, Coorens THH, Hooks Y, O’Neill L, Latimer C, et al. Extensive heterogeneity in somatic mutation and selection in the human bladder. Science. 2020;370(6512):75–82.

89. Shi MJ, Meng XY, Fontugne J, Chen CL, Radvanyi F, Bernard-Pierrot I. Identification of new driver and passenger mutations within APOBEC-induced hotspot mutations in bladder cancer. Genome Med. 2020;12(1):85.

90. Matheus LHG, Dalmazzo SV, Brito RBO, Pereira LA, De Almeida RJ, Camacho CP, et al. 1-Methyl-D- tryptophan activates aryl hydrocarbon receptor, a pathway associated with bladder cancer progression. BMC Cancer. 2020;20(1):869.

91. Yu J, Lu Y, Muto S, Ide H, Horie S. The Dual Function of Aryl Hydrocarbon Receptor in Bladder Carcinogenesis. Anticancer Res. 2020;40(3):1345–57.

92. Kang SU, Park JT. Functional evaluation of alternative splicing in the FAM190A gene. Genes and Genomics. 2019;41(2):193–9.

93. Patel K, Scrimieri F, Ghosh S, Zhong J, Kim MS, Ren YR, et al. FAM190A deficiency creates a cell division defect. Am J Pathol. 2013;183(1):296–303.

94. Lindskrog SV, Prip FF, Lamy P, Taber A, Clarice S, Nordentoft I, et al. An integrated multi-omics analysis identifies clinically relevant molecular subtypes of non-muscle-invasive bladder cancer. 2020;

95. Stein JP, Lieskovsky G, Cote R, Groshen S, Feng AC, Boyd S, et al. Radical cystectomy in the treatment of invasive bladder cancer: Long-term results in 1,054 patients. J Clin Oncol. 2001;19(3):666–75.

96. Mari A, Campi R, Tellini R, Gandaglia G, Albisinni S, Abufaraj M, et al. Patterns and predictors of recurrence after open radical cystectomy for bladder cancer: a comprehensive review of the literature. Vol. 36, World Journal of Urology. Springer Verlag; 2018. p. 157–70.

97. Marine J-C, Dawson S-J, Dawson MA. Non-genetic mechanisms of therapeutic resistance in cancer. Nat Rev Cancer. 2020;20:1–14.

98. Pivot XB, Bondarenko I, Dvorkin M, Trishkina E, Ahn J-H, Im S-A, et al. A randomized, double-blind, phase III study comparing SB3 (trastuzumab biosimilar) with originator trastuzumab in patients treated by neoadjuvant therapy for HER2-positive early breast cancer. J Clin Oncol. 2017;35(15_suppl):509–509.

99. Rosenberg JE, O’Donnell PH, Balar A V., McGregor BA, Heath EI, Yu EY, et al. Pivotal Trial of Enfortumab Vedotin in Urothelial Carcinoma After Platinum and Anti-Programmed Death 1/Programmed Death Ligand 1 Therapy. J Clin Oncol. 2019;37(29):2592–600.

100. Boni V, Sharma MR, Patnaik A. The Resurgence of Antibody Drug Conjugates in Cancer Therapeutics: Novel Targets and Payloads. Am Soc Clin Oncol Educ B. 2020;40:e58–74.

101. Sheng X, Yan X, Wang L, Shi Y, Yao X, Luo H, et al. Open-label, Multicenter, Phase II Study of RC48-ADC, a HER2-Targeting Antibody–Drug Conjugate, in Patients with Locally Advanced or Metastatic Urothelial Carcinoma. Clin Cancer Res. 2021;27(1):43–51.

102. Delmore JE, Issa GC, Lemieux ME, Rahl PB, Shi J, Jacobs HM, et al. BET bromodomain inhibition as a therapeutic strategy to target c-Myc. Cell. 2011;146(6):904–17.

103. Goldstein JT, Berger AC, Shih J, Duke FF, Furst L, Kwiatkowski DJ, et al. Genomic activation of PPARG reveals a candidate therapeutic axis in bladder cancer. Cancer Res. 2017;77(24):6987–98.

104. Peng H, Wisse E, Tian Z. Liver natural killer cells: Subsets and roles in liver immunity. Cell Mol Immunol. 2016;13(3):328–36.

105. Ochoa MC, Minute L, Rodriguez I, Garasa S, Perez-Ruiz E, Inogés S, et al. Antibody-dependent cell cytotoxicity: immunotherapy strategies enhancing effector NK cells. Immunol Cell Biol. 2017;95(4):347–55.

106. Moore AR, Rosenberg SC, McCormick F, Malek S. RAS-targeted therapies: is the undruggable drugged? Nat Rev Drug Discov. 2020;19(8):533–52.

107. Meng XM, Nikolic-Paterson DJ, Lan HY. TGF-β: The master regulator of fibrosis. Nat Rev Nephrol. 2016;12(6):325–38.

108. Borthwick LA, Wynn TA, Fisher AJ. Cytokine mediated tissue fibrosis. Biochim Biophys Acta - Mol Basis Dis. 2013;1832(7):1049–60.

109. van Dijk N, Gil-Jimenez A, Silina K, Hendricksen K, Smit LA, de Feijter JM, et al. Preoperative ipilimumab plus nivolumab in locoregionally advanced urothelial cancer: the NABUCCO trial. Nat Med. 2020;26(12):1839–1844.

110. Lv J, Li P. Mesothelin as a biomarker for targeted therapy. Biomark Res. 2019;7(1):18.

111. Dunham I, Kundaje A, Aldred SF, Collins PJ, Davis CA, Doyle F, et al. An integrated encyclopedia of DNA elements in the human genome. Nature. 2012;489(7414):57–74.

112. Roadmap Epigenomics Consortium, Kundaje A, Meuleman W, Ernst J, Bilenky M, Yen A, et al. Integrative analysis of 111 reference human epigenomes. Nature. 2015;518(7539):317–29.

