## Supplementary figures for "Molecular characterization reveals genomic and transcriptomic subtypes of metastatic urothelial carcinoma"

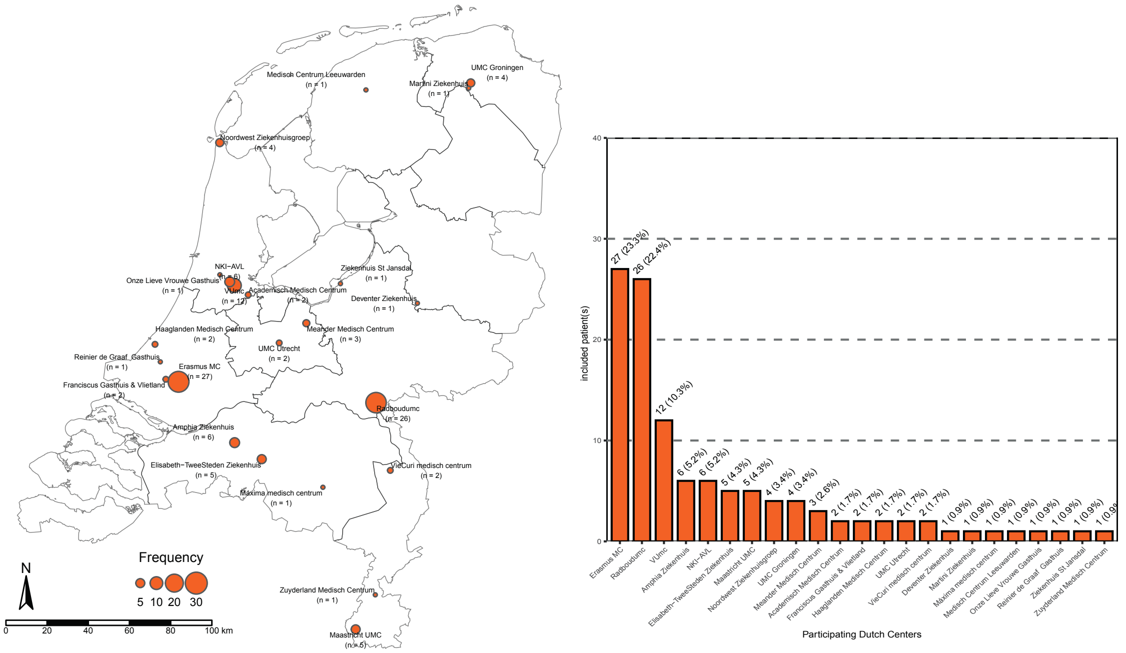


**Figure S1. Participating hospitals and patients with metastatic urothelial carcinoma accrual per hospital**

Overview of Dutch hospitals participating in the study of the Center for Personalized Cancer Treatment (CPCT) consortium, and the number of mUC patients included in the study per hospital.

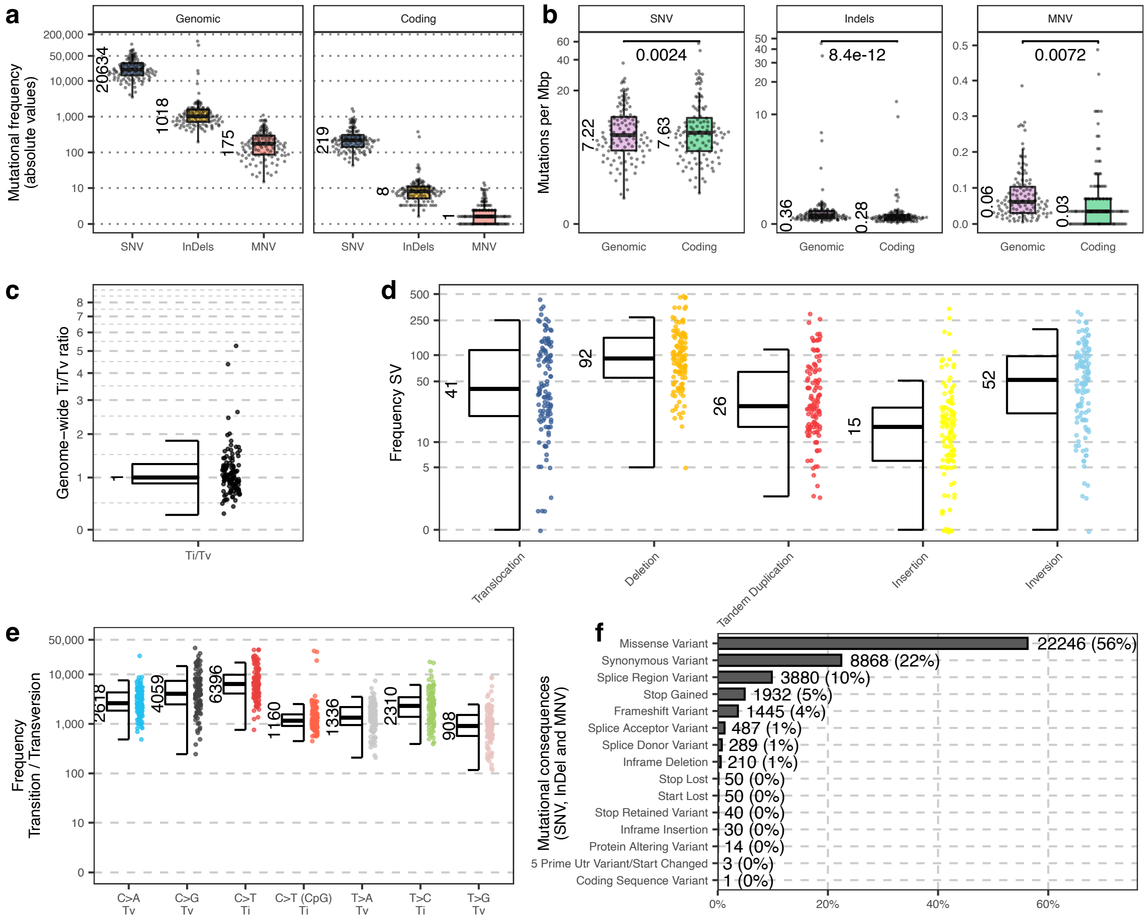


**Figure S2. Overview of the mutational landscape of 116 metastatic urothelial carcinoma samples**

(a) Boxplot of the total number of single nucleotide variants (SNVs), insertion/deletions (Indels) and multiple nucleotide variants (MNVs) at whole-genomic level and in coding region. The median value of the entire cohort is shown next to each boxplot.

(b) Comparison of SNVs, Indels and MNVs per mega base-pair (Mbp) between genome-wide and coding region. The median is shown next to each boxplot. Wilcoxon rank sum test p-values are shown for each comparison.

(c) Boxplot with median value and individual data points of genome-wide ratio of transitions (Ti) over transversions (Tv).

(d) Boxplot and frequency of interchromosomal translocations, deletions, tandem duplications (TD); insertions and inversions are indicated per sample and with median value.

(e) Boxplot of genome-wide SNV types. Transition (Ti) and transversion (Tv), with a special attention for C to T in CpG context are indicated per sample with median value.

(f) Barplot of mutational consequences of genomic variants at protein level over the entire cohort using Ensembl variant effect predictor.

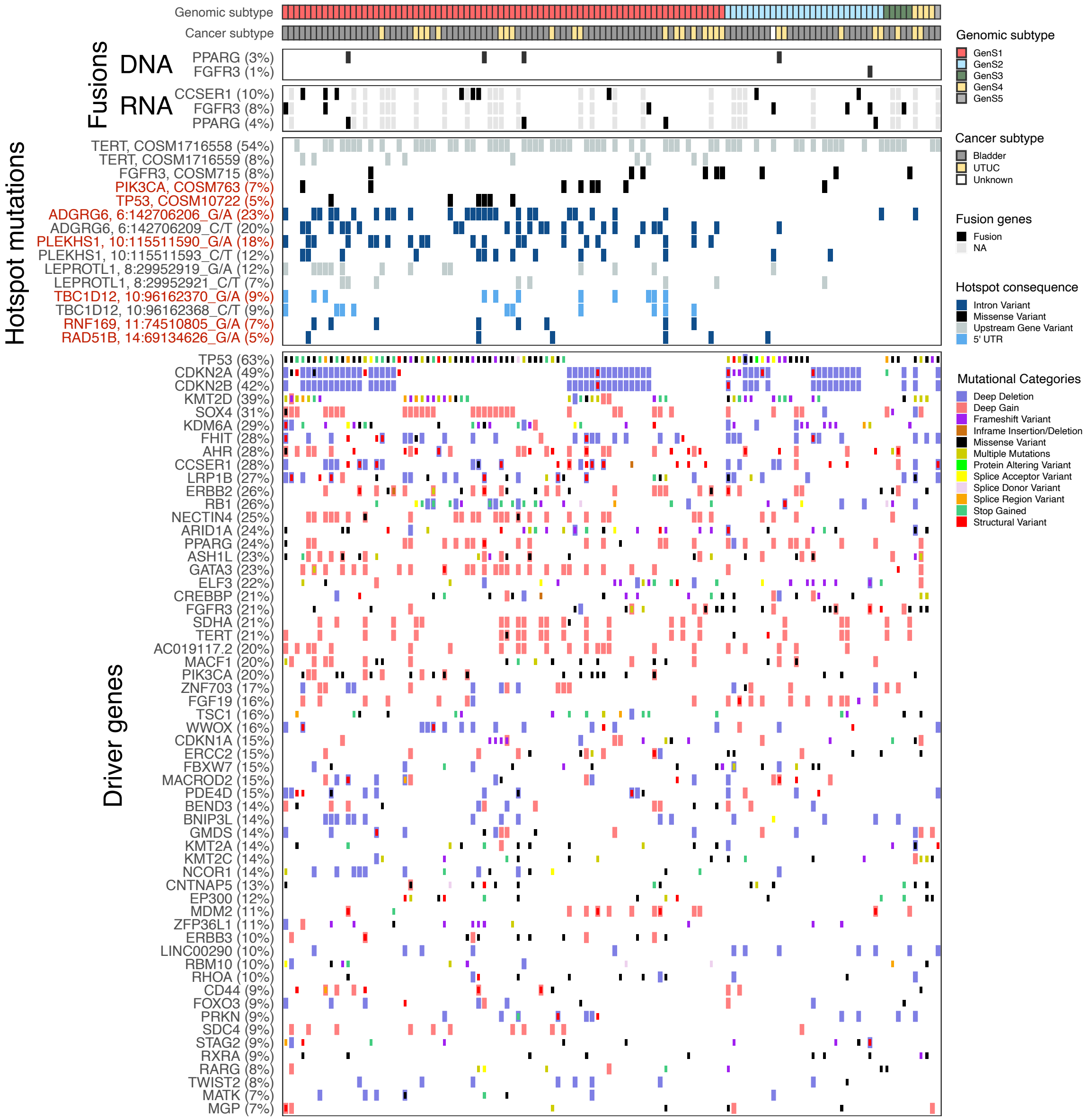


**Figure S3. Gene fusions (DNA- and RNA-based detection), hotspot mutations and driver genes distribution across the genomic subtypes**

Overview of recurrent gene fusion were detected at RNA and DNA level. Samples without RNA-seq are shown in gray. Hotspot mutations that may be related to APOBEC activity (C>T and C>G in TCW context) are marked in red. The oncoplot shows significantly mutated genes estimated with dNdScv [1] and with GISTIC2 [2]. Other known driver genes recurrently mutated were also included. Cancer subtype is indicated.

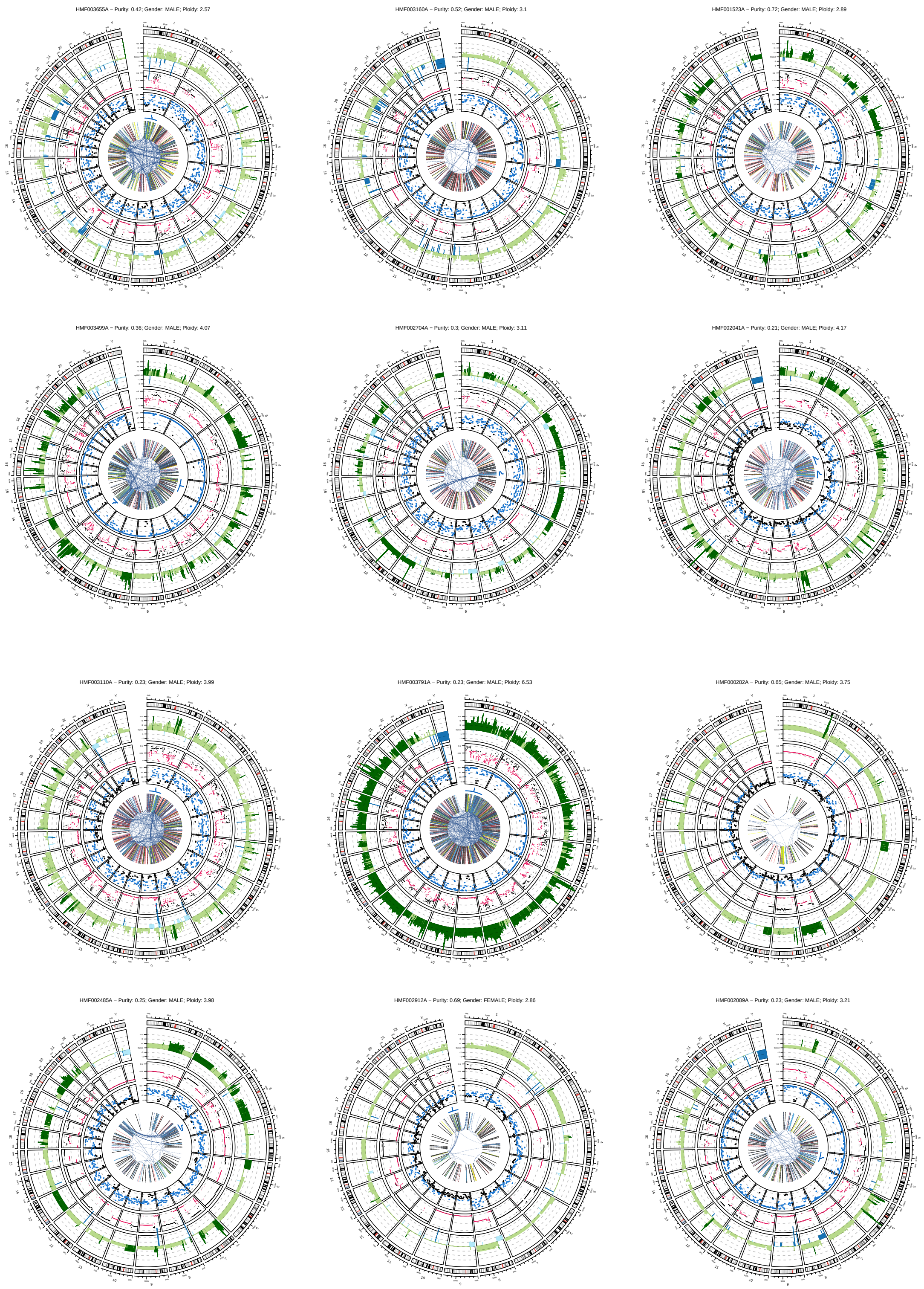


**Circos plot of samples with chromothripsis**

Description continues on next page.

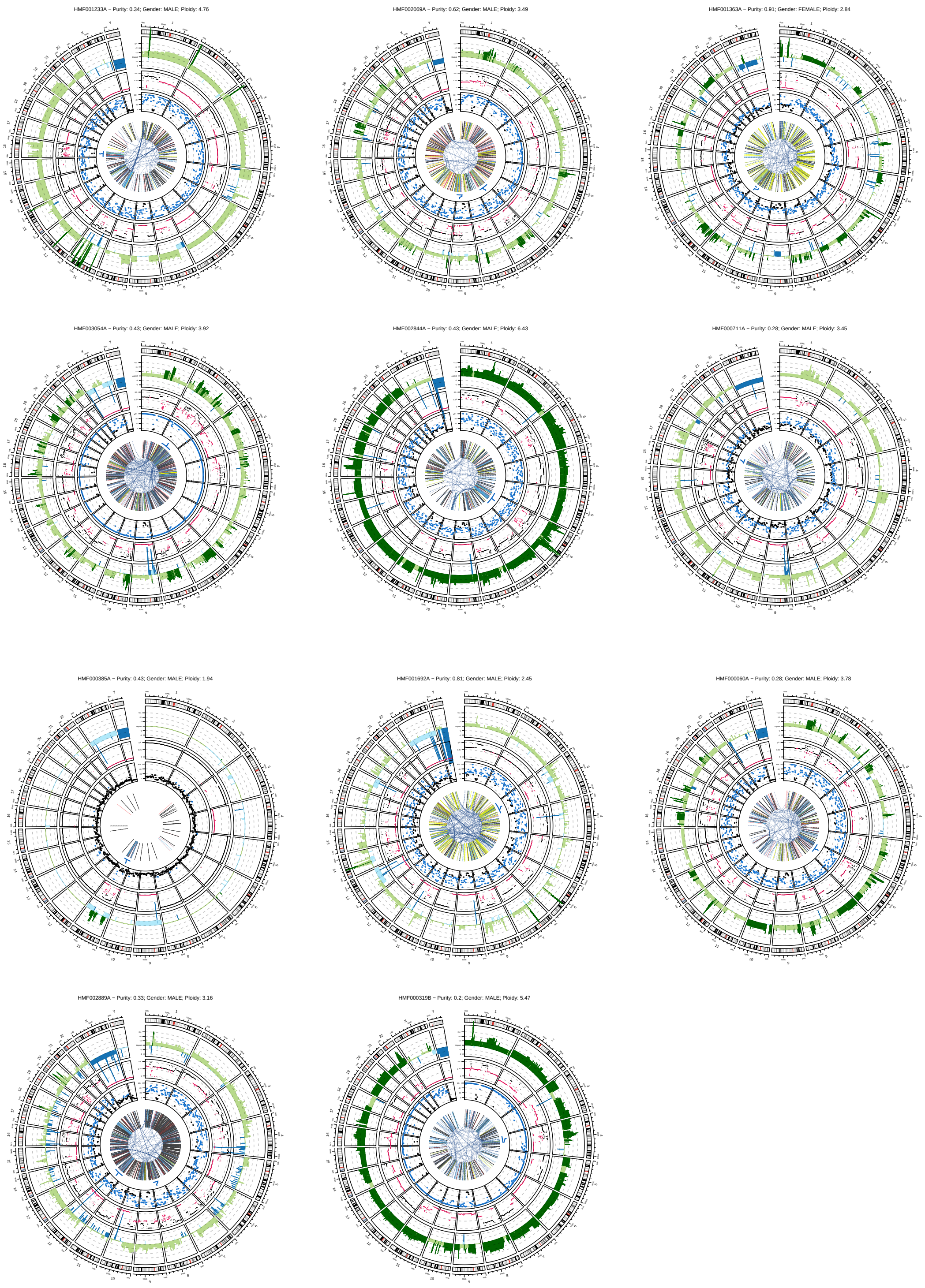


**Figure S4. Circos plot of samples with chromothripsis**
The circos plot shows from outer to inner circles: the genomics ideogram from chromosome 1 to X where the centrosomes are indicated in red; Copy number estimated with GISTIC2 [2]; B allele frequency (red for <0.33 and black for >0.33); Mutational load (number of mutations per 5 Mbp; black for < 20 and blue for >20); Chromothripsis regions in blue lines; Structural variants (break ends in black; deletions in gray, insertions in yellow, inversions in blue and duplications in red).

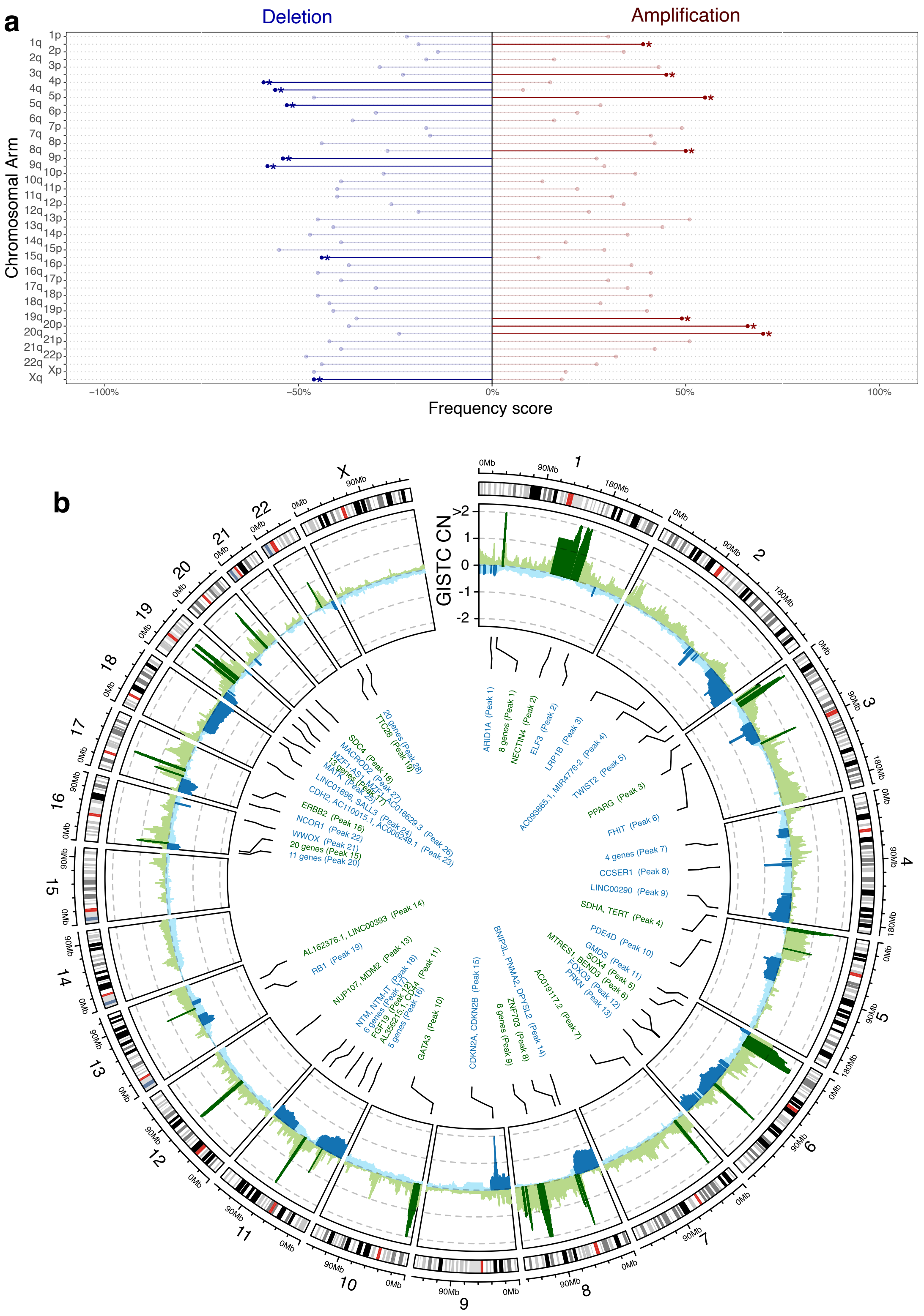

**Figure S5. Recurrent copy number changes at chromosomal arm and gene level calculated with GISTIC2**

(a) Significant chromosomal arm deletions and amplifications (q < 0.05) indicated in darker colors.
(b) Circos plot with significant (q < 0.05) focal copy number changes (narrow peaks). Deleted/amplified genes are shown for each peak.

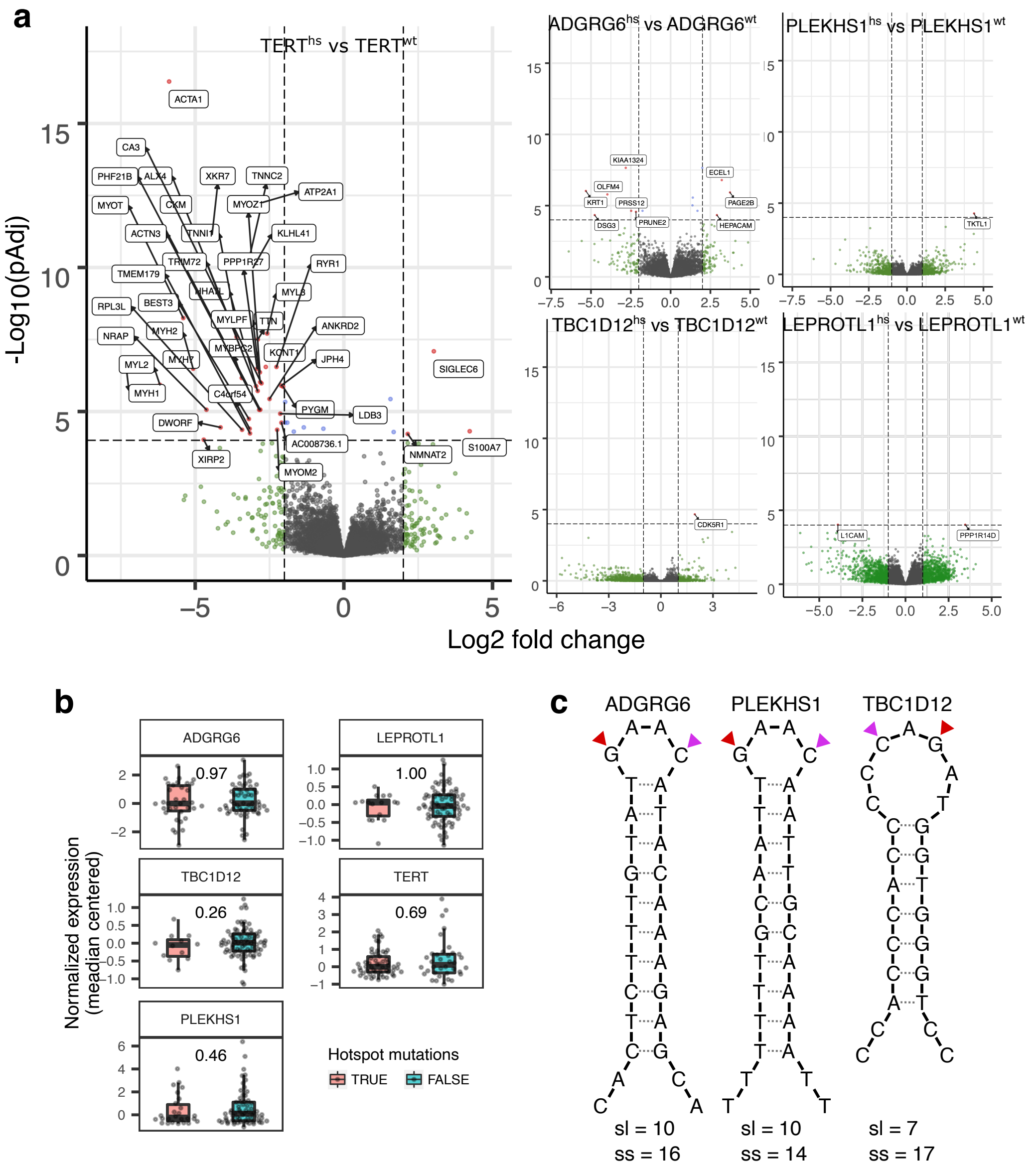


**Figure S6. Consequences of hotspot mutations in non-coding regions of *TERT*, *PLEKHS1*, *TBC1D12* and *ADGRG6***

(a) Volcano plot of differential gene expression analysis between tumors with hotspot mutations ($TERT^{hs}$, $PLEKHS1^{hs}$, $TBC1D12^{hs}$, $ADGRG6^{hs}$ and $LEPROTL^{hs}$) and tumors without hotspot mutations ($TERT^{wt}$, $PLEKHS1^{wt}$, $TBC1D12^{wt}$, $ADGRG6^{wt}$ and $LEPROTL^{wt}$) in non-coding region of *TERT*, *PLEKHS1*, *TBC1D12*, *ADGRG6* and *LEPROTL*. Log2 fold change represents gene expression of tumors with hotspot-mutant gene relative to gene expression of tumors without hotspot mutations.

(b) RNA expression of genes affected by hotspot mutations (*TERT*, *PLEKHS1*, *TBC1D12*, *ADGRG6* and *LEPROTL*) was compared between tumors with and without the mutated hotspot. BH corrected p-values are shown for Wilcoxon rank-sum test.

(c) Scheme of possible hairpin loop structures formed in the regions frequently affected by hotspot mutations in *ADGRG6*, *PLEKHS1* and *TBC1D12*. The most frequent positions of hotpots mutations are indicated with a red triangle, other recurrent positions are indicated with a purple triangle. The most frequent hotspot mutation occurs in APOBEC context (C>T in TCW context). The stem length (sl) and stem strength (ss) for each hairpin loop structure is indicated. The stem strength is calculated as the sum of base pairs, where A-T takes the value of one and C-G takes the value of three [3].

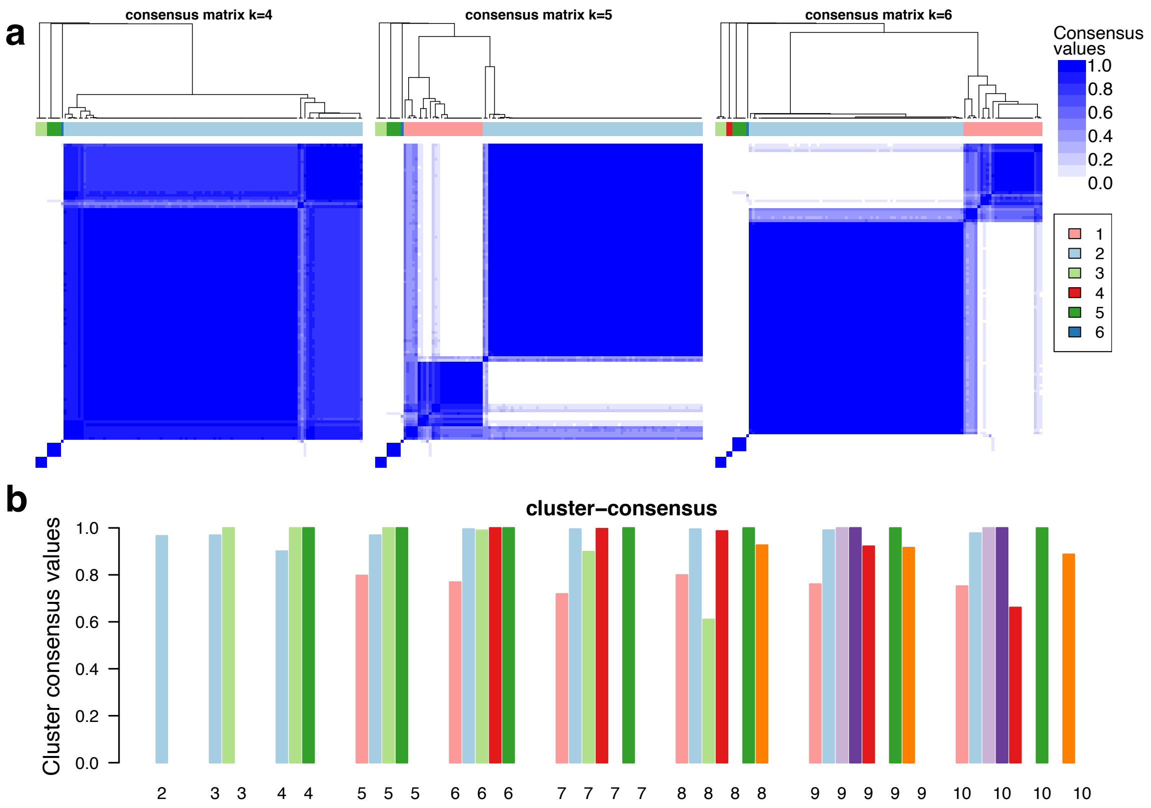


**Figure S7. Consensus matrix of mutational signatures**

(a) Three consensus matrices with k=4-6 estimated from the etiology of mutational signatures.
(b) Mean cluster stability indicated for k=2 to k=10. Stability was calculated by resampling 80% of the data points repeated 1000 times.

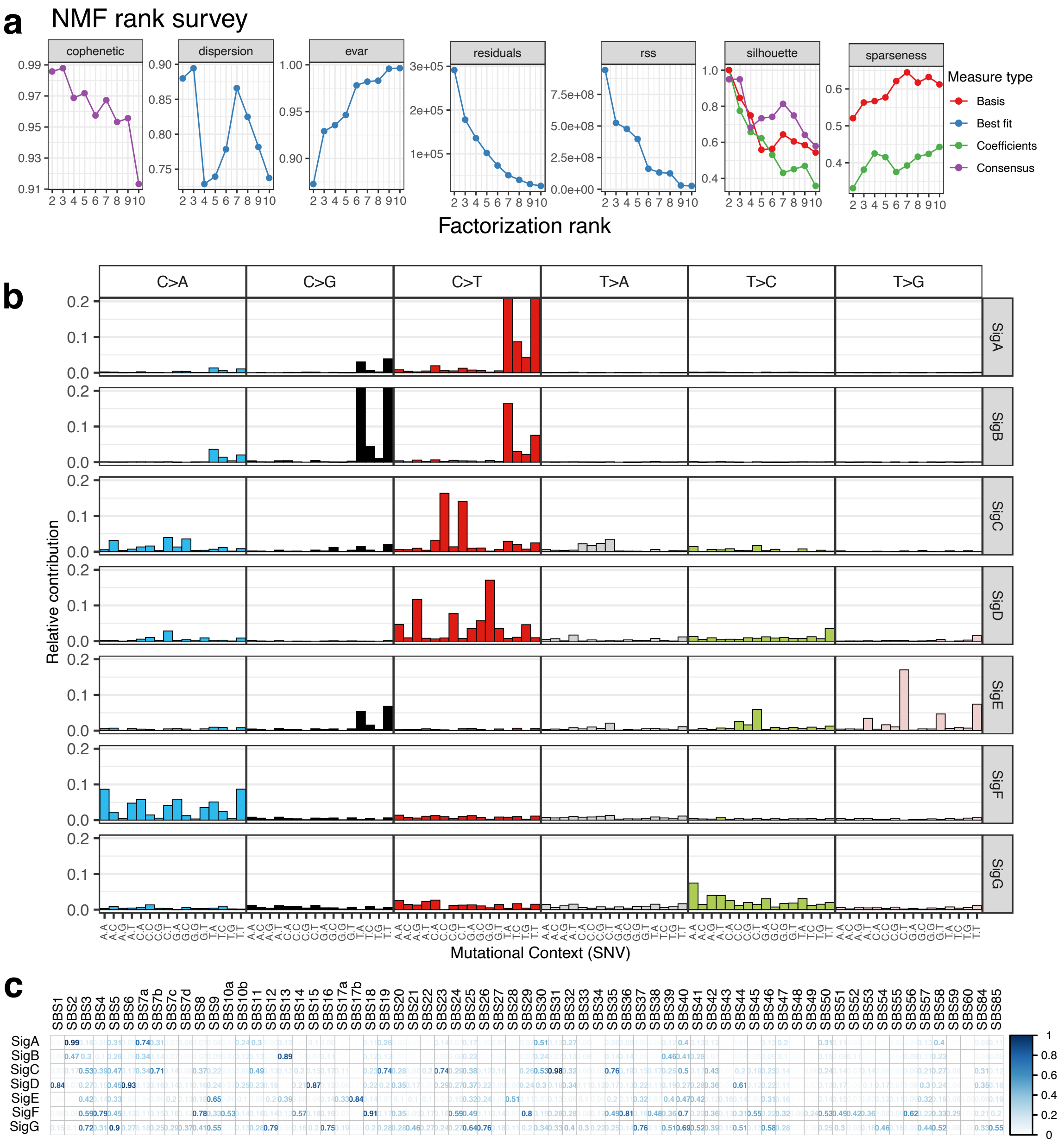


**Figure S8.** ***De novo*** **mutational signatures estimated from all SNVs considering the tri-nucleotide context**

(a) Results from non-negative matrix factorization to estimate *de novo* mutational signatures from k=2 to k=10.

(b) Mutational profile of seven de novo signatures. Mutations are shown in their 96 tri-nucleotide context.

(c) Cosine similarity of *de novo* signatures with all mutational signatures from COSMIC v3. Cosine similarity values are shown for each pairwise comparison and colored in darker blue for higher values.

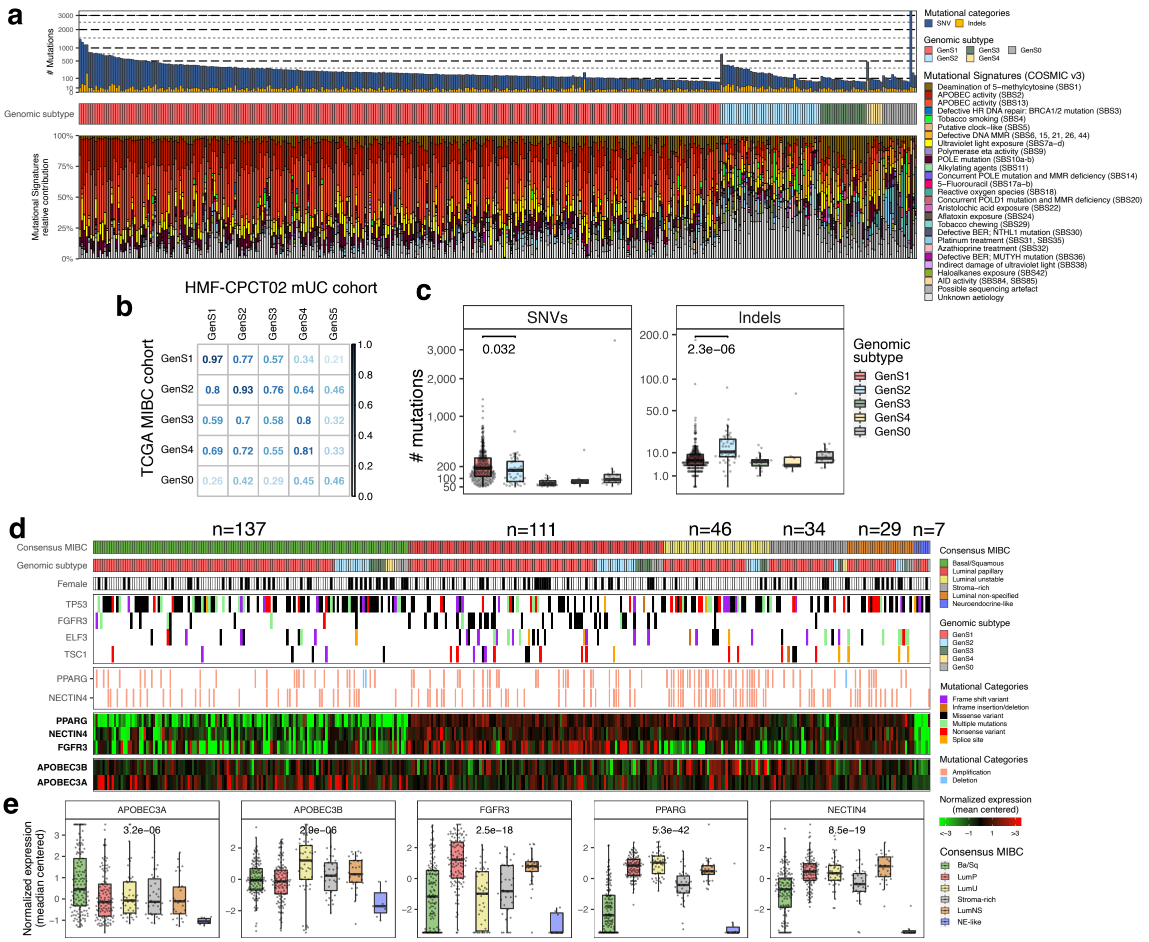


**Figure S9. Genomic and transcriptomic subtypes of the TCGA primary bladder cancer cohort**

(a) Publicly available WES data (n = 367) of the TCGA bladder cancer cohort were analyzed following the same method used in this study. ConsensusClusterPlus [4] was applied on the etiology of mutational signatures COSMIC v3 to divide the cohort into five genomic subtypes (GenS0-4). Tumor mutational burden (single nucleotide variants (SNVs) and insertions/deletions (Indels)) and mutational signatures are shown per tumor.

(b) Cosine similarity matrix of genomic subtypes of muscle invasive bladder cancer (MIBC) from the TCGA cohort with the metastatic urothelial carcinoma cohort analyzed in this study.

(c) Number of SNVs and indels across genomic subtypes. P-values are shown for Wilcoxon rank-sum test between GenS1 and GenS2.

(d) 364 tumors from the TCGA cohort were stratified using the consensus MIBC classifier on RNA-seq data [5]. Genomic subtypes, tumors from female patients (in black), genomic alterations of selected driver genes (*TP53, FGFR3, ELF3, TSC1, PPARG* and *NECTIN4*) and expression of selected genes (*PPARG, NECTIN4, FGFR3, APOBEC3A* and *APOBEC3B*) are displayed.

(e) Comparison of the expressions of selected genes between the transcriptomic consensus MIBC subtypes. Kruskal-Wallis test p-values were BH corrected.

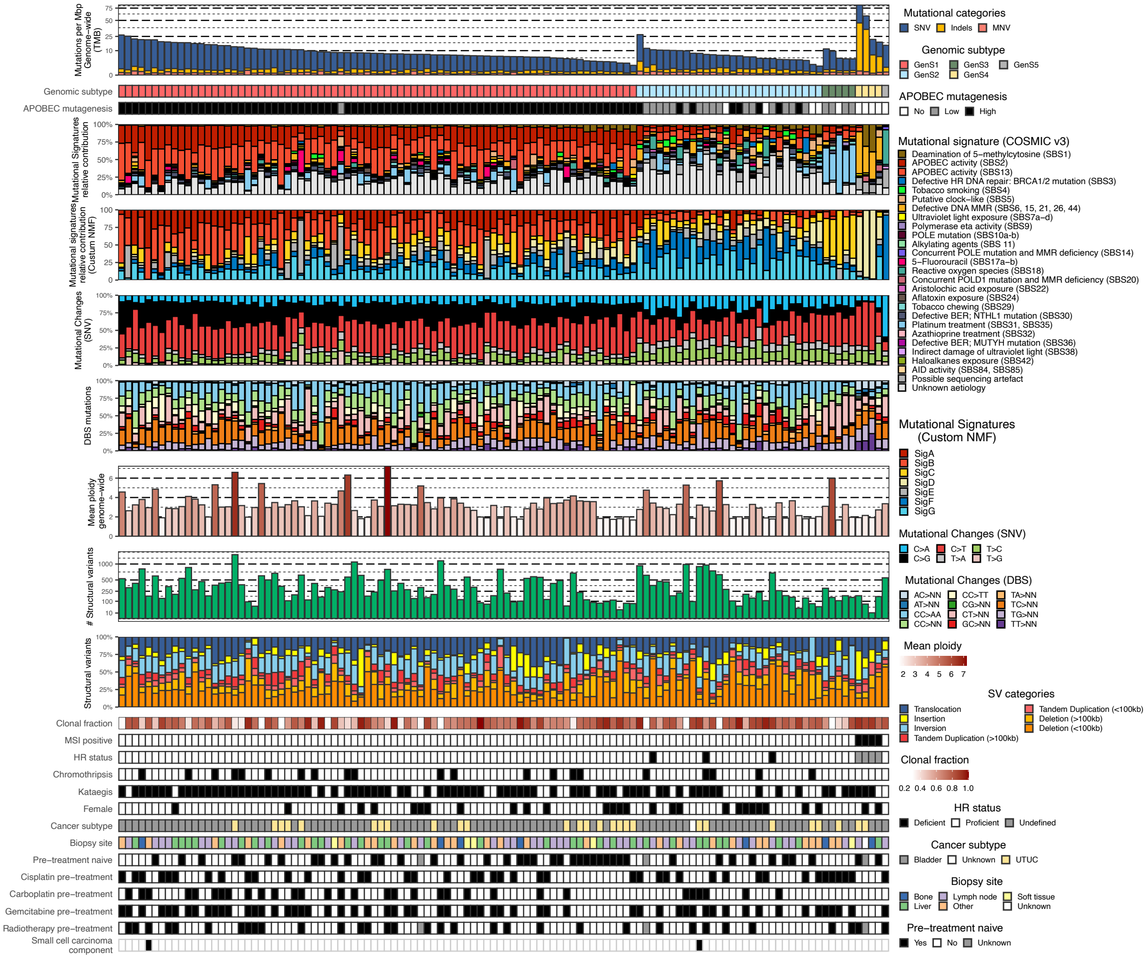


**Figure S10. Complete genomic landscape of the metastatic Urothelial Carcinoma cohort**

Patients were stratified based on the etiology of mutational signatures. The genomic features are displayed from top to bottom: Genome-wide tumor mutation burden (TMB; mutations per Mbp); Five genomic subtypes (GenS); APOBEC enrichment analysis showing tumors with APOBEC mutagenesis high, low and no APOBEC mutagenesis; mutational signatures COSMIC v3 grouped by etiology; Relative contribution of seven custom mutational signatures; Relative frequency of types of SNV; Relative frequency of double base substitution events (DBS); Genome-wide mean ploidy; Number of structural variants; Relative frequency of different types of structural variants; Relative amount of SNVs that are clonal; Tumors with MicroSatellite Instability (MSI); Homologous Recombination (HR) deficiency; Chromothripsis detected in tumors; Kataegis events detected in tumor; Female patients; Primary origin of metastatic sample (UTUC, UCB or unknown); Location of biopsy sample; Pre-treatment naïve patients; Cisplatin pre-treatment; Carboplatin pre-treatment; Gemcitabine pre-treatment; Radiotherapy pre-treatment; Small cell carcinoma component.

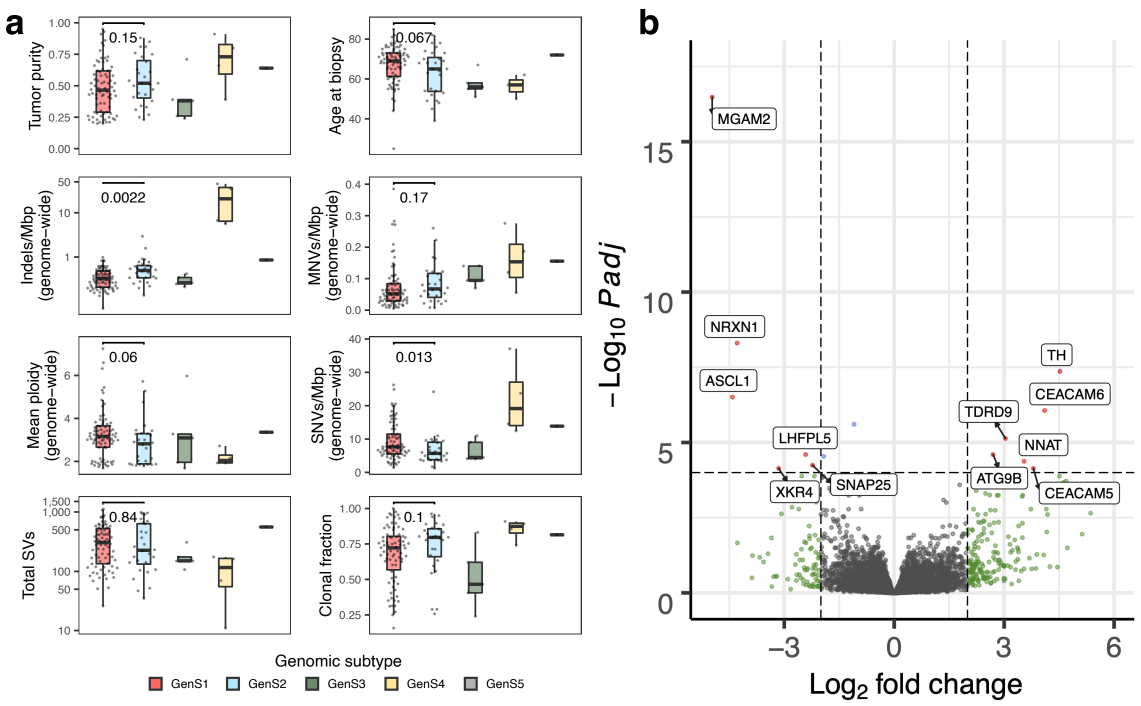


**Figure S11. Comparing the genomics and transcriptomics features between all genomic subtypes**

(a) Distribution of some genomic and transcriptomic features among genomic subtypes. Age at biopsy and tumor purity are also shown. Wilcoxon rank-sum test p-values are shown for comparisons of GenS1 with GenS2.

(b) Volcano plot of differential gene expression analysis between GenS1 and GenS2. Fold changes were log2 transformed and represent expression values of genes in GenS1 over GenS2.

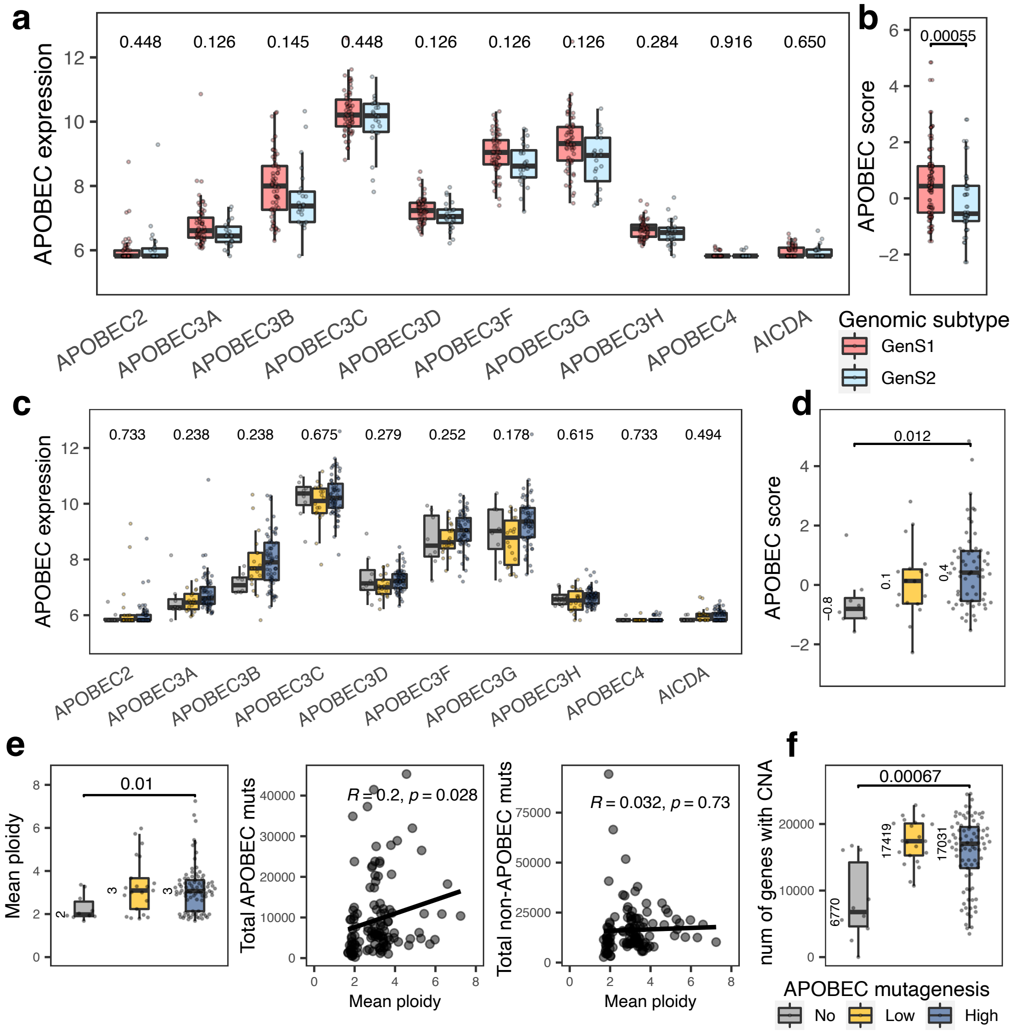


**Figure S12. Molecular differences between tumors with and without enriched APOBEC mutagenesis**

(a) Normalized mRNA-expression of enzymes of the APOBEC family of genomic subtypes GenS1 (mostly APOBEC-high mutation enrichment tumors) and GenS2 (mostly APOBEC-low mutation enrichment tumors). Wilcoxon rank-sum test was applied, and p-values were BH corrected for multiple hypothesis testing.

(b) APOBEC score calculated as the sum of the mean centered normalized mRNA-expression of *APOBEC3A* and *APOBEC3B*. P-value is shown for Wilcoxon rank-sum test.

(c) RNA-expression of different APOBEC enzymes compared between APOBEC and non-APOBEC mediated mutagenesis tumors. Kruskal-Wallis test p-values were BH corrected.

(d) APOBEC score across groups of tumors with distinct level of enriched APOBEC mutagenesis. Wilcoxon rank-sum test p-value is shown for non-APOBEC and high APOBEC mediated mutagenesis tumors.

(e) Ploidy differences between APOBEC and non-APOBEC mediated mutagenesis tumors. P-value of Wilcoxon rank-sum test is shown for APOBEC-high and non-APOBEC enriched tumors. Spearman's correlation coefficient was estimated on the number of APOBEC and non-APOBEC associated mutations with genome-wide ploidy.

(f) The number of genes affected by shallow or deep copy number changes (calculated with GISTIC2) is higher in tumors with enriched APOBEC mutagenesis. Wilcoxon rank-sum test p-value is shown for the comparison between APOBEC-high mutagenesis tumors and tumors with no evidence of APOBEC mutagenesis.

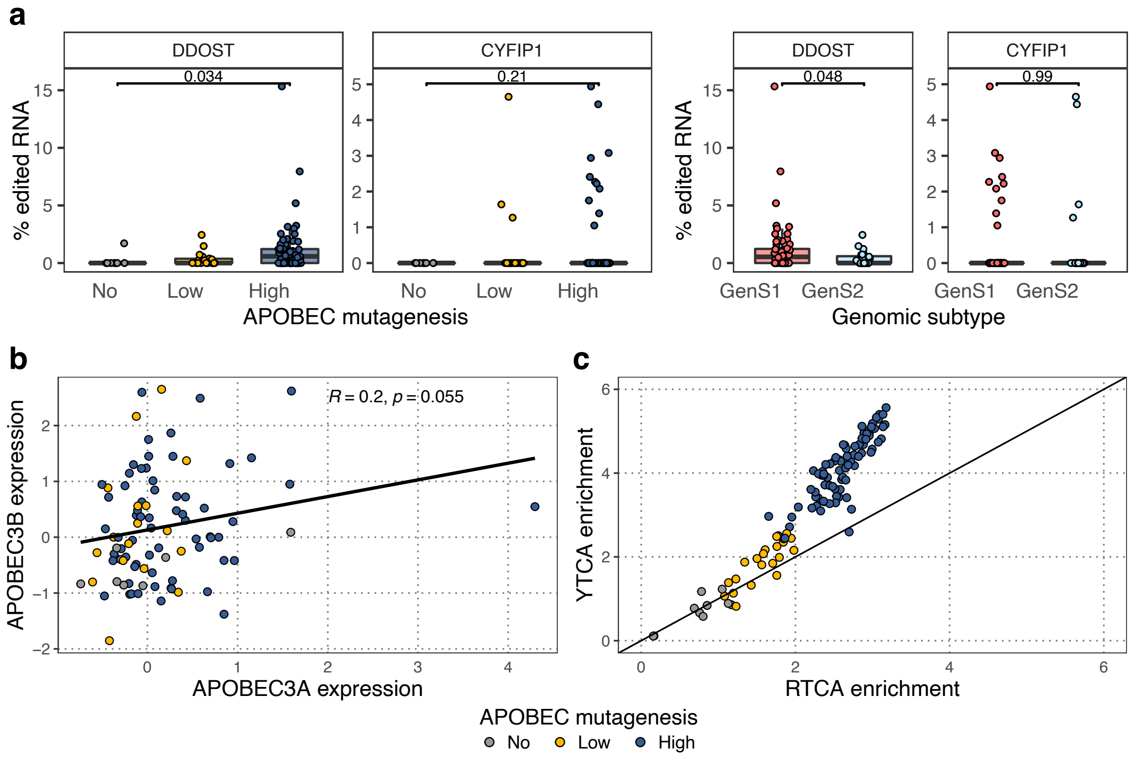


**Figure S13. APOBEC3A and APOBEC3B activity**
(a) Percentage of hotspot mutations in mRNA of *DDOST* (chr1:20981977) and *CYFIP1* (chr15:22999350). Wilcoxon rank-sum test was applied.
(b) Linear correlation of APOBEC3A and APOBEC3B expression. Pearson correlation coefficient was estimated. Expression was estimated by normalizing raw counts with DESeq2 [6] and final values were median centred.
(c) Fold enrichment of C>G and C>T mutations in YTCA (Y = T or C) and RTCA (R = G or A) context. Samples with equal enrichment of YTCA and RTCA mutations should coincide with the black line.

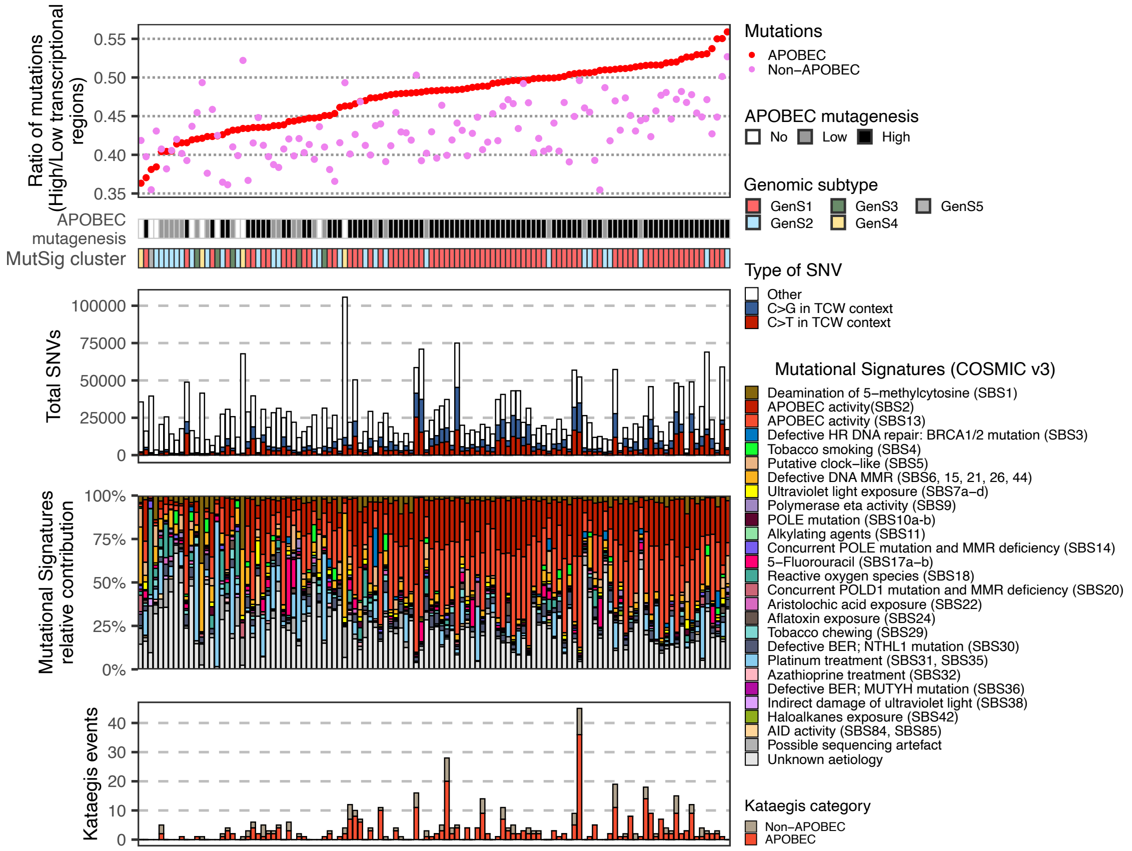


**Figure S14. Ratio of APOBEC mutations between high and low transcriptional regions**

The ratio of APOBEC associated mutations between high and low transcriptional regions, APOBEC mutagenesis, genomic subtype, total number of single nucleotide variants (SNVs), mutational signatures and number of kataegis events are displayed per sample. Samples were stratified according to the ratio of APOBEC mutations. Most tumors with high APOBEC mutagenesis have a ratio close to 0.5 (0.45-0.55). This ratio is, in the majority of tumors, lower for non-APOBEC mutations. When the number of APOBEC associated mutations is low (tumors with low APOBEC mutagenesis or tumors with no APOBEC mutagenesis), the ratio becomes smaller and in most cases is bellow 0.45, which is expected as non-APOBEC mutations dominate.

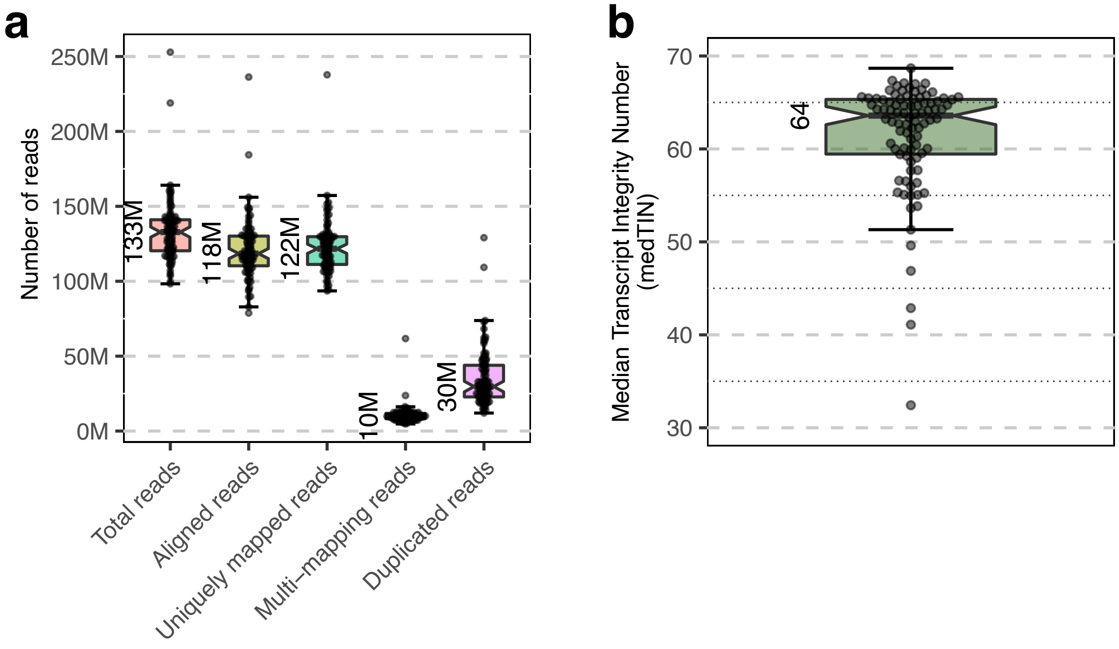


**Figure S15. Different metrics showing quality of RNA sequencing**
(a) Alignment results from 90 RNA-seq samples (excluding the second biopsy of seven tumors). Median values of the entire cohort are shown for each metric.
(b) The median transcript integrity number was estimated for each sample (excluding the second biopsy of seven tumors).

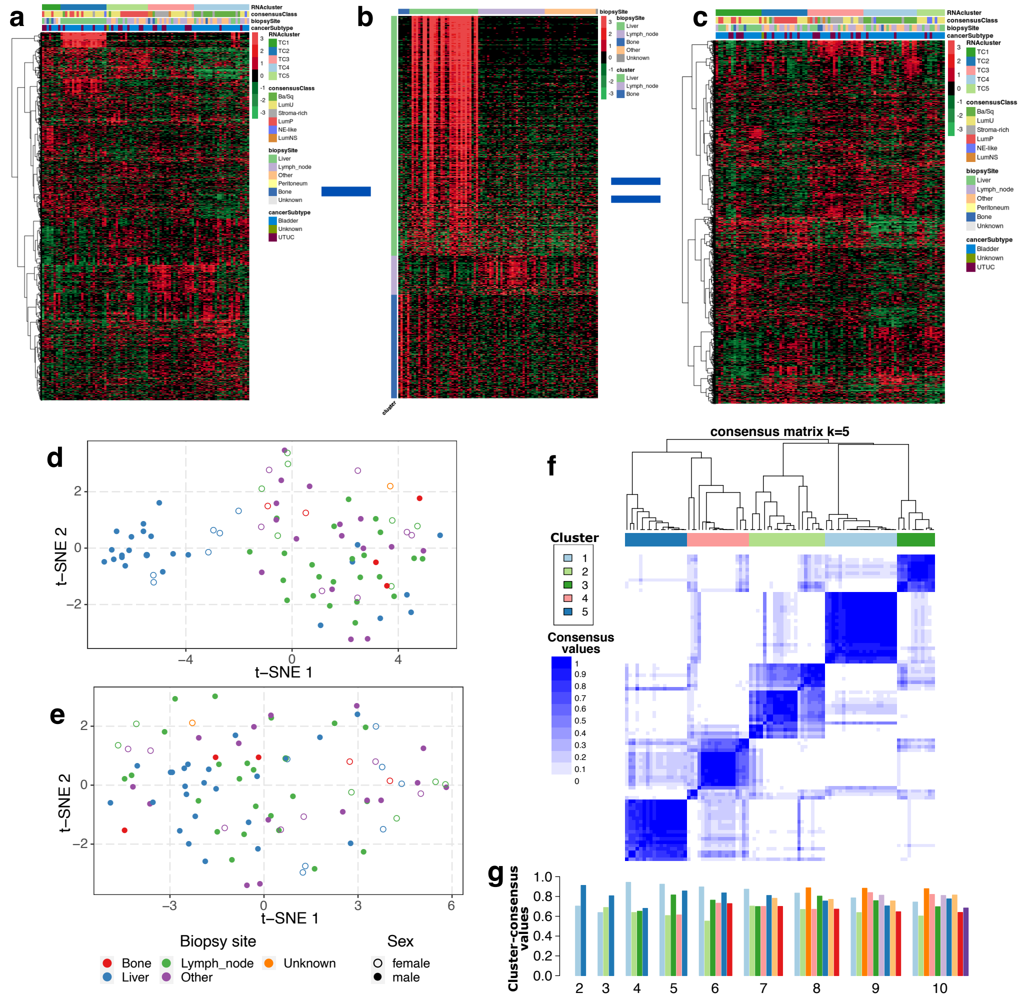

**Figure S16. Strategy followed to discard organ-specific transcripts of metastatic urothelial carcinoma**

(a) All transcripts were normalized using DESeq2 [6]; only the 50% most variable (highest variance) transcripts with base mean > 100 were used for downstream analysis. Normalized counts of transcripts were median centered; the transcriptomic profiles were used to cluster tumors with ConsensusClusterPlus [4] This panel shows the influence of organ-specific transcripts on clustering analysis. Note that liver specific transcripts form one cluster. The consensus MIBC classifier [5] was applied to the same RNA-seq data. Biopsy site and cancer subtype (bladder, UTUC = upper tract urinary cancer and unknown) are indicated per tumor.

(b) Organ-specific transcripts were identified through differential gene expression analysis with DESeq2 [6]. Transcripts that contributed most to each organ (biopsy site) were identified (log2 fold change > 1 and BH corrected p-values < 0.05). Note that liver is the organ with most differentially expressed transcripts.

(c) Organ-specific transcripts were removed from the RNA-seq data; clustering was performed with ConsensusClusterPlus on the remaining transcripts.

(d) t-SNE from panel (a) showing the influence of organ specific transcripts (mainly liver).

(e) t-SNE from panel (c) showing how the effect of organ-specific transcripts had disappeared after removing these transcripts from the RNA-seq data.

(f) Consensus matrix with transcripts from panel (c).

(g) Cluster stability in each cluster. From the clustering analysis, a mean cluster consensus value was obtained as a measure of cluster stability. Increasing the number of clusters increases the stability but creates smaller clusters as well. Thus, the criterium for selecting five clusters was based on cluster stability and the size of the clusters (not allowing clusters with <5 samples).

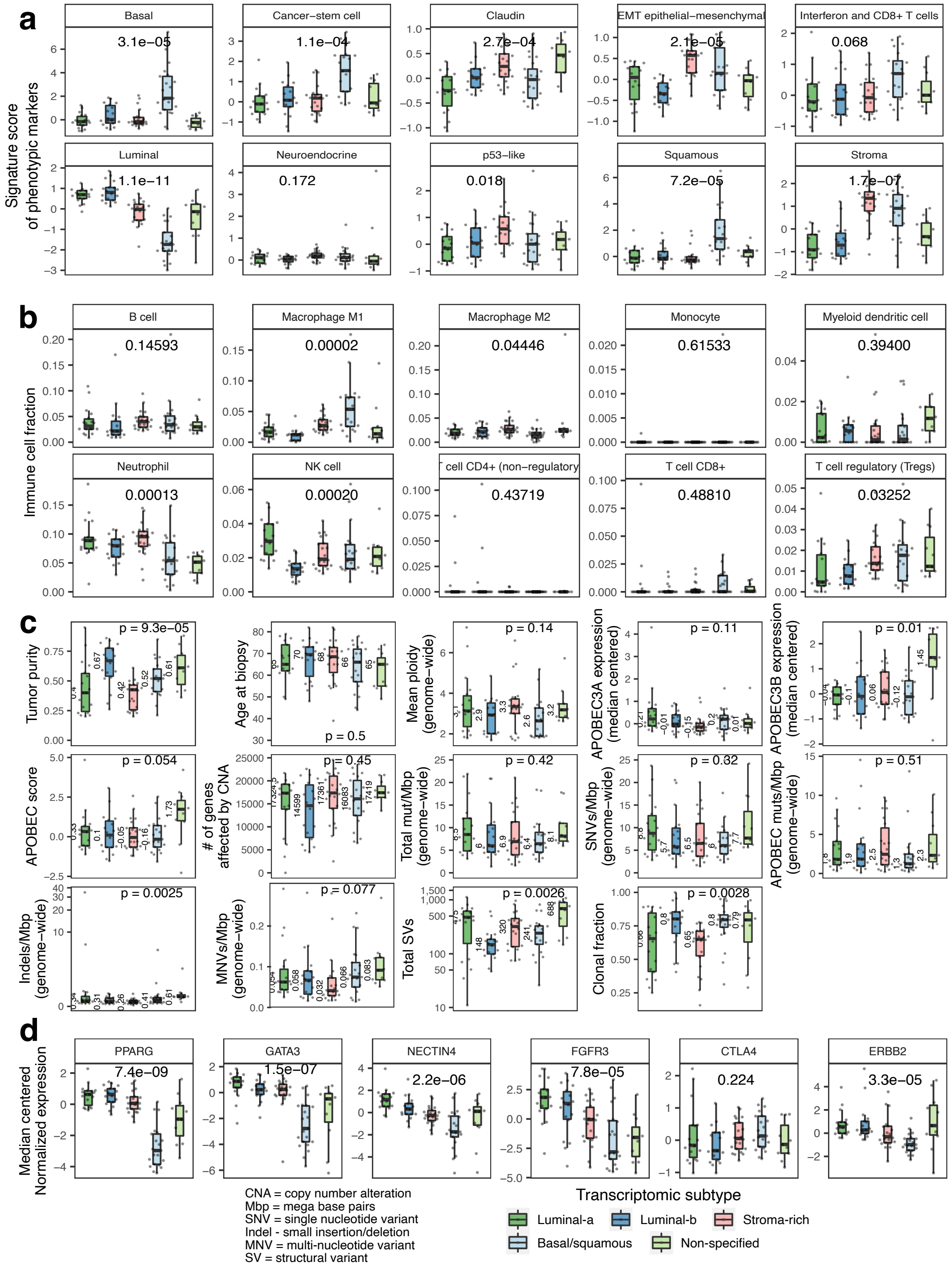


**Figure S17. Molecular differences between the five transcriptomic subtypes of metastatic urothelial carcinoma**

(a) Signature score of phenotypic markers estimated as the mean expression of genes associated with each phenotype. Kruskal-Wallis test p-values were BH corrected.

(b) Immune cell fraction estimated with immunedeconv [7], using the quanTIseq method [8]. Kruskal-Wallis test p-values were BH corrected.

(c) Tumor purity, genomic (mean ploidy, number of genes affected by copy number alterations, mutational load, clonal fraction), transcriptomic (APOBEC enzymes expression) and clinical data (age at biopsy) were compared. Kruskal-Wallis test p-value for each comparison is shown.

(d) Expression of selected genes. Kruskal-Wallis test p-values were BH corrected.

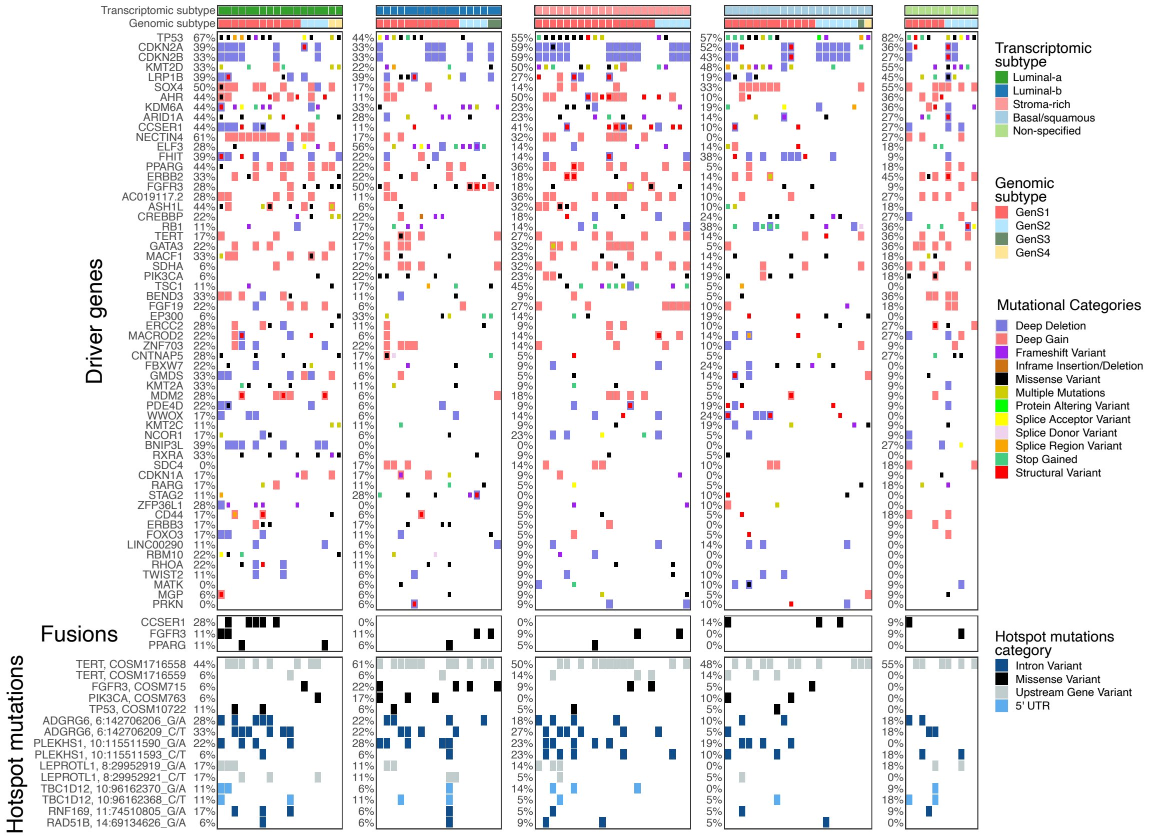


**Figure S18. Mutational landscape per transcriptomic subtype of metastatic urothelial carcinoma**

Overview of mutations in driver genes, gene fusions and hotspot mutations described in the metastatic urothelial carcinoma cohort across transcriptomic subtypes. Driver genes estimated from dNdScv [1], GISTIC2 [2] and frequently mutated genes known to be drivers in urothelial carcinoma are displayed.

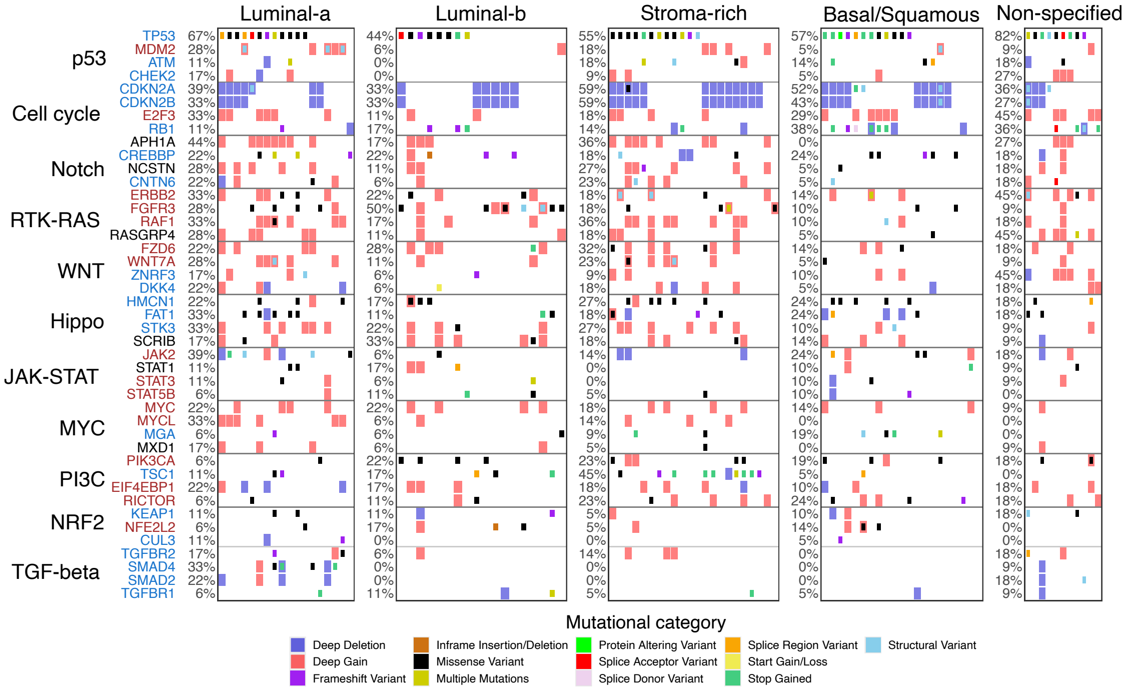


**Figure S19. Top four mutated genes in eleven oncogenic pathways distributed across transcriptomic subtypes**

Genes of eleven oncogenic pathways were assessed for genomic alterations. The top four altered genes are shown here. Names of tumor suppressor genes and oncogenes are colored in blue and red, respectively.

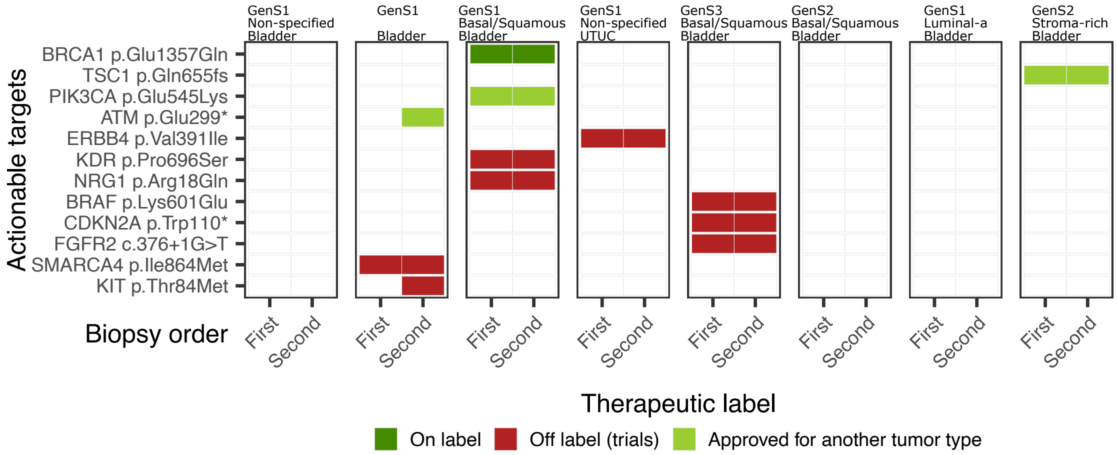


**Figure S20. Actionable targets identified in tumors with multiple biopsy samples** Actionable targets in patients with multiple biopsies were identified and compared. Targets found in the first biopsy are found back again in the second biopsy. In one patient (HMF001286), new targets were identified. The therapy for one of these targets is approved by the Food and Drug Administration for another tumor type. Only targets for single nucleotide variants are shown.

### Supplementary references


[1] Martincorena I, et al. (2017) Universal patterns of selection in cancer and somatic tissues. *Cell* 171(5):1041.e21.

[2] Mermel CH, et al. (2011) Gistic2.0 facilitates sensitive and confident localization of the targets of focal somatic copy-number alteration in human cancers. *Genome Biology* 12(4):R41.

[3] Buisson R, et al. (2019) Passenger hotspot mutations in cancer driven by apobec3a and mesoscale genomic features. *Science* 364(6447).

[4] Wilkerson MD, Hayes DN (2010) Consensusclusterplus: a class discovery tool with confidence assessments and item tracking. *Bioinformatics* 26:1572–1573.

[5] Kamoun A, et al. (2020) A consensus molecular classification of muscle-invasive bladder cancer. *European Urology* 77(4):420–433.

[6] Love MI, Huber W, Anders S (2014) Moderated estimation of fold change and dispersion for rna-seq data with deseq2. *Genome Biology* 15(12):550.

[7] Sturm G, et al. (2019) Comprehensive evaluation of transcriptome-based cell-type quantification methods for immuno-oncology. *Bioinformatics* 35(14):i436–i445.

[8] Finotello F, et al. (2019) Molecular and pharmacological modulators of the tumor immune contexture revealed by deconvolution of rna-seq data. *Genome Medicine* 11(1):34.
